# Quantifying target antigen-dependent CAR T-cell performance against AML

**DOI:** 10.1101/2024.12.16.628628

**Authors:** Saumil Shah, Jan Mueller, Emanuel Vogel, Michael Raatz, Steffen Boettcher, Arne Traulsen, Markus G. Manz, Philipp M. Altrock

## Abstract

Chimeric Antigen Receptor (CAR) T-cell therapy has transformed cancer immunotherapy by genetically engineering T-cells to target tumor antigens. Acute myeloid leukemia (AML) presents unique challenges due to resistance mechanisms, especially in patients with TP53 loss mutations. The complex dynamics of CAR T-cell expansion remain poorly understood. The field lacks validated quantitative frameworks to systematically evaluate different CAR T-cell target constructs, such as CD33, CD123, and CD371, against resistant AML variants. We address this gap by combining mathematical modeling with *in vitro* assay data and Bayesian inference. We select, train, and validate a two-compartment deterministic mathematical model that describes the nonlinear dynamics of target AML and CAR T cells, accounting for expansion, killing, and exhaustion. Using Bayesian inference, we train and select the best-performing functional form for CAR T expansion and then validate it on unseen data. Our framework selects a CAR T-cell expansion model that accounts for handling time and T-cell self-interference, highlighting that expansion is a dynamic process in which target-cell handling time and T-cell crowding negatively affect T-cell expansion. Analysis of posterior parameter distributions reveals target-antigen-specific responses against TP53-deficient AML. For instance, CD33-targeting CARs have reduced attack rates against TP53-deficient cells, while CD123- and CD371-targeting CARs show moderately increased attack rates; however, the former exhibit higher death rates, and the latter have increased handling times, impeding efficacy. This target-dependent form of resistance challenges the assumption of uniform performance and reveals a unifying nonlinear expansion model for integrated, yet antigen-specific, preclinical predictions of efficacy.

**Short Summary:** We identified the target antigen-specific expansion properties and the efficacy of CAR T cells against *TP53* wild-type and deficient AML by integrating experimental data from cytolytic assays, mechanistic mathematical modeling, and Bayesian inference.

## Introduction

Chimeric antigen receptor (CAR) T-cell therapy, a type of adoptive T-cell therapy, has significantly advanced personalized cancer immunotherapy for many hematologic malignancies^1^. Patients’ T-cells genetically modified to express a synthetic receptor with an extracellular antibody-derived fragment that binds to tumor antigens in an MHC-independent way^2^. These CAR T-cells are then usually expanded *ex vivo* and infused back, where they recognize their target, expand, and attack target cells. The recent successes against B-cell acute lymphoblastic leukemia (B-ALL), B-cell non-Hodgkin lymphoma (B-NHL) have not yet been replicated in myeloid cancers or many solid tumors^3^.

Myeloid malignancies, chiefly acute myeloid leukemia (AML) and myelodysplastic syndrome (MDS), are neoplastic diseases of early myeloid progenitors^4^. The observation that allogeneic stem cell transplants are effective in AML via graft-versus-leukemia effects suggests that AML blast cells are susceptible to T-cell predation, providing a rationale for CAR T-cell therapy^5^. Unfortunately, AML has been less amenable to CAR T-cell therapy, primarily due to a lack of suitable target antigens^6,7^. Multiple distinct myeloid surface antigens are typically expressed by leukemic blast cells and have been targeted by CAR T-cells in preclinical and clinical studies, including CD33, CD117, CD371, and CD123^8–10^. Despite these extensive efforts, a suitable target antigen for CAR T-cells in AML/MDS has not been described^6^. The choice of target antigen for an engineered T-cell therapy can profoundly impact the expansion of CAR T-cells^11^.

*TP53* is a well-studied tumor suppressor gene that acts as a transcription factor in response to cellular stress. *TP53* loss in AML and MDS is associated with therapy resistance and dismal prognosis^12,13^. *TP53* mutations affect the microenvironment, inhibit cancer cell apoptosis, promote immune evasion of cancer cells^14^, and have been shown to confer resistance to CAR T-cell cytotoxicity ^8^. Patients suffering from AML/MDS, especially the aggressive and largely incurable *TP53*-deficient variant, represent an unmet clinical need for CAR T-cell therapy^15^.

Insufficient dose and expansion negatively correlate with treatment efficiency and clinical response rates^16^. Prolonged expansion of cytotoxic CAR T-cells is also associated with T-cell exhaustion and poor treatment outcomes^17^. Thus, understanding the non-monotonic behavior of T-cell expansion is critical for the development and design of novel CAR T-cell therapies and treatment protocol^18^. Further, the importance of achieving favorable E:T ratios has been observed in different clinical contexts^19^. An integrated quantitative understanding of T-cell expansion, cytotoxicity, and exhaustion across E:T ratios is key to preclinical and translational studies of novel CAR T-cell therapy approaches. These investigations benefit from mathematical models that relate mechanistic cell population dynamics to outcome^20^. A challenge is associating realistic assumptions with parameter estimates^21^, particularly CAR T expansion, killing, and contraction^22,23^. Functional response is a classical concept that describes these processes^24–29^.

Here, we use functional response modeling to investigate the dynamics of T or CAR T-cells engaging acute myeloid leukemia blasts. We examine a pool of candidate models combining *in vitro* cytotoxic assays, mathematical modeling, and Bayesian inference^30^. We provide a framework for obtaining parameter distributions and CAR T-leukemia interaction models with nested or competing assumptions from longitudinal data, elucidating how *TP53* status affects the functional response of the proportion of killed target leukemia cells as a function of T-cell density. While the methods presented here cannot reflect clinical diversity in its entirety. Our primary goal is to use rigorous Bayesian inference to train, select, and validate the best-performing CAR T expansion function in the context of novel killing assays and, as such, to demonstrate that this approach is a feasible strategy for establishing preclinical knowledge. Our approach and results demonstrate a viable method for quantitative systems modeling of the performance of novel CAR T-cell constructs, powered by state-of-the-art statistical inference.

## Methods

We aimed to quantify the dynamic performance of T-cells against acute myeloid leukemia cells observed *in vitro* (leveraging four unique experimental settings, **Figure 1A**) using a mathematical model (**Figure 1B**). The overall workflow for assessing cell population growth longitudinally is shown in **Figure 1C**. The applied suite of Ordinary Differential Equations (ODE) models differ in the assumed T or CAR T-cell expansion rate function (**Figure 1D**). For all resulting submodels, we performed identifiability analyses, implemented Bayesian inference integrating pre-existing knowledge (priors of parameter value distributions), obtained posterior parameter distributions, and performed model selection and validation (**Figure 1E**).

**Figure 1:**
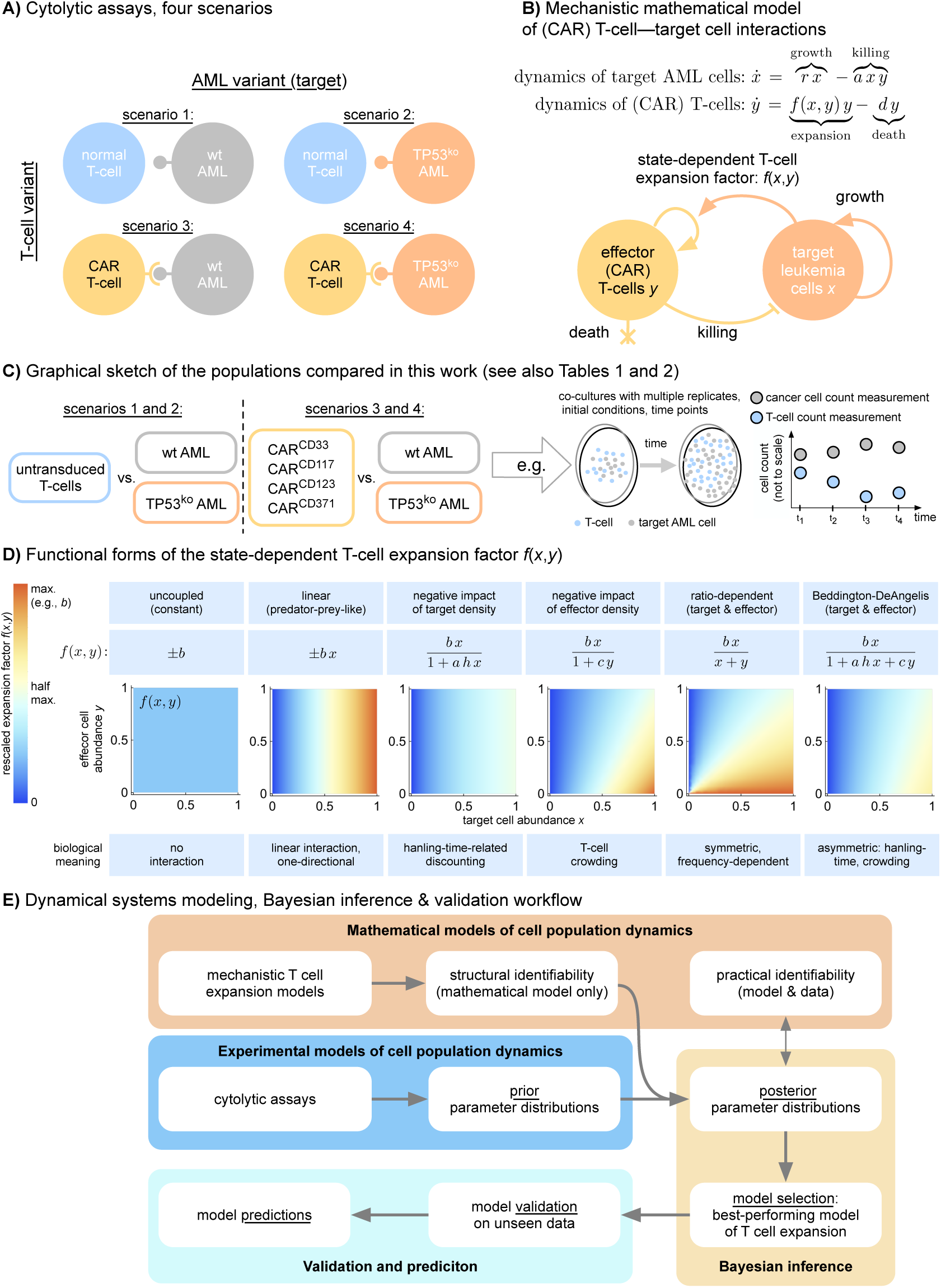
Identifiability and Bayesian inference workflow, candidate expansion functions, and structural identifiability of parameters. **A**) The cytolytic assay employed four experimental scenarios. **B**) We used a two-compartment mathematical model to study the performance of the CAR T-cell expansion function *f*(*x*,*y*). We use the abbreviation *dx*⁄*dt* = *ẋ* (*x*: target cell abundance, *y*: T-cell abundance). **C**) In the four scenarios, longitudinal data of matching untransduced T- and CAR T-cells were compared with one of the two AML variants at various E:T ratios (see Methods); we show a cartoon of one such co-culture experiment comparing untransduced T-cells and wt AML cells over time. **D**) CAR T expansion can depend on *x* and *y*. We studied the performance of multiple candidate expansion functions, each with a unique biological meaning. **E**) Statistical and modeling workflow illustration. The goodness of fit was evaluated using the negative log-likelihood, Akaike, and Bayesian information criteria (AIC, BIC). Validation was performed on previously unseen assay data using a correlation function (Methods). Predictions were made by implementing the model, inferring parameter posteriors, and using realistic conditions.

### Cell-culture experiments

Using a CRISPR/Cas9-based gene editing approach, the *TP53* gene was knocked out from acute myeloid leukemia (MOLM13) tumor cell line^31^. Lentiviral transduction of green fluorescent protein (GFP) and luciferase followed by fluorescence-activated cell sorting yielded *TP53*-wildtype (MOLM13-TP53^+/+ *GFP* + *Luc* +^), and *TP53*-knockout (MOLM13-*TP53*^−/− *GFP* + *Luc* +^) tumor cell populations. The target cells were cultured in Roswell-Park-Memorial-Institute (RPMI) medium supplemented with 100 U⁄*ml* penicillin, 100 *μg*⁄*ml* streptomycin, and 10% fetal bovine serum (FBS) at 37°C, 5% CO_+_. Human effector cells were isolated from healthy donor buffy coats and transduced with lentiviral vectors produced in HEK-293 cells to obtain second-generation CAR T-cells. Published sequences of single-chain variable-fragment (scFv) targeting CD33 (clone SGN-33, Lintuzumab^32^), CD117 (clone D79^33^), CD123 strong (CD123s, clone H9^34^), CD123 weak (CD123w, clone CSL362 ^35^), and CD371 (clone M26 ^36^) were used for the transduction together with a CD8 stalk and transmembrane domain and a 4-1BB intracellular activating domain. After transduction, CAR T-cells were purified by magnetic-activated cell sorting (MACS) and expanded in the presence of IL-2. After two weeks of expansion, aliquots of 10^7^ pure CAR T-cells were frozen in liquid nitrogen (LN2) until use in experiments. The target and effector cells were co-cultured in variable ratios, seeded with 60,000 total cells for the cytotoxic assay, repeated twice to obtain two technical replicates per measurement. The reagents and antibodies in **Supplementary Table 1** were used for flow cytometry-based analysis of co-incubation assays. Data were acquired on an LSRFortessa™ 4L equipped with a high-throughput sampler (HTS) (Becton Dickinson, Franklin Lakes, NJ, USA) and analyzed with the FlowJo® v9x software (Becton Dickinson, Franklin Lakes, NJ, USA). The total target and effector cells were measured using the described flow-based approach on days 1, 3, 6, and 10 for one experimental model setting (LT15) and on days 1, 3, 6, 10, and 16 for another (smaller) experimental model setting (termed LT6).

We verified that the MOLM-13 TP53-wildtype and TP53-knockout lines proliferate comparably over the timescale of the cytotoxic co-culture assays. In the absence of effector cells, the two genotypes showed overlapping cumulative fold expansion from day 7 through day 19 (**Supplementary Figure 3**), spanning the full duration of both the training (LT15, day 10) and validation (LT6, day 16) datasets. A modest divergence between WT and KO emerged only at longer culture times, beyond the experimental window of this study. Consistent with the within-window equivalence, the posterior distributions of the target growth rate inferred from the uT-vs-WT and uT-vs-KO co-culture conditions overlapped substantially across all five construct experiments (**Supplementary Figure 9**). Together, these observations rule out intrinsic differences in target-cell growth as a driver of the target-antigen-dependent CAR T kinetic signatures reported below.

### Data integration

The dataset contained two types of target cells, *TP53*-wildtype and *TP53*-knockout, and two effector cell types: untransduced T-cells and transduced CAR T effector cells. This resulted in 4 treatment combinations: untransduced T vs. *TP53*-wildtype tumor, untransduced T vs. *TP53*-knockout tumor, chimeric antigen receptor T vs. *TP53*-wildtype tumor, and chimeric antigen receptor T vs.*TP53*-knockout tumor. Aliquots from the cytotoxic assay were analyzed using fluorescence-activated cell sorting for tumor marker CD33+ and T-cell marker CD3+ counts. The counts were then multiplied by the dilution factor of the aliquot to approximate the total number of cells in the co-culture. Each chimeric antigen receptor target single-chain variable fragment (scFv) had a unique set of four initial E:T ratios shown in **Table 1**. The data frame is presented in **Table 2**. To avoid differences-in-magnitude numerical problems across E:T ratios, we divided the cellular abundance data of the entire data frame by 10^5^ for data fitting, bringing all cell counts to order 1.

**Table 1:**
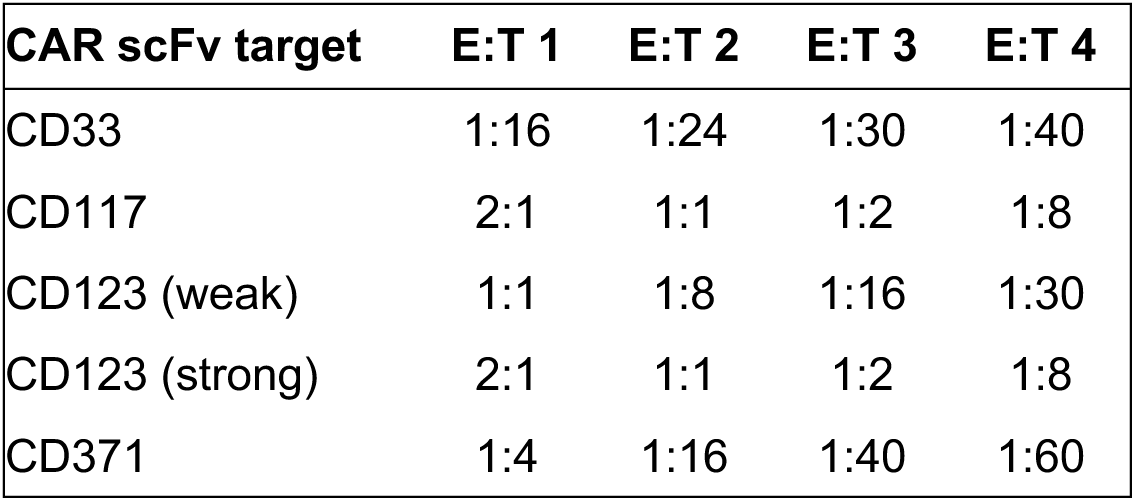
Initial E:T ratio for each chimeric antigen receptor construct cytotoxic assay. Each experiment was seeded with 60000 cells and the four listed E:T ratios. For example, the cytotoxic assay using CD123-targeting (weak) CAR T-cells (first initial condition) was seeded with 40000 effector and 20000 target cells.

| CAR scFv target | E:T 1 | E:T 2 | E:T 3 | E:T 4 |
| --- | --- | --- | --- | --- |
| CD33 | 1:16 | 1:24 | 1:30 | 1:40 |
| CD117 | 2:1 | 1:1 | 1:2 | 1:8 |
| CD123 (weak) | 1:1 | 1:8 | 1:16 | 1:30 |
| CD123 (strong) | 2:1 | 1:1 | 1:2 | 1:8 |
| CD371 | 1:4 | 1:16 | 1:40 | 1:60 |

**Table 2:**
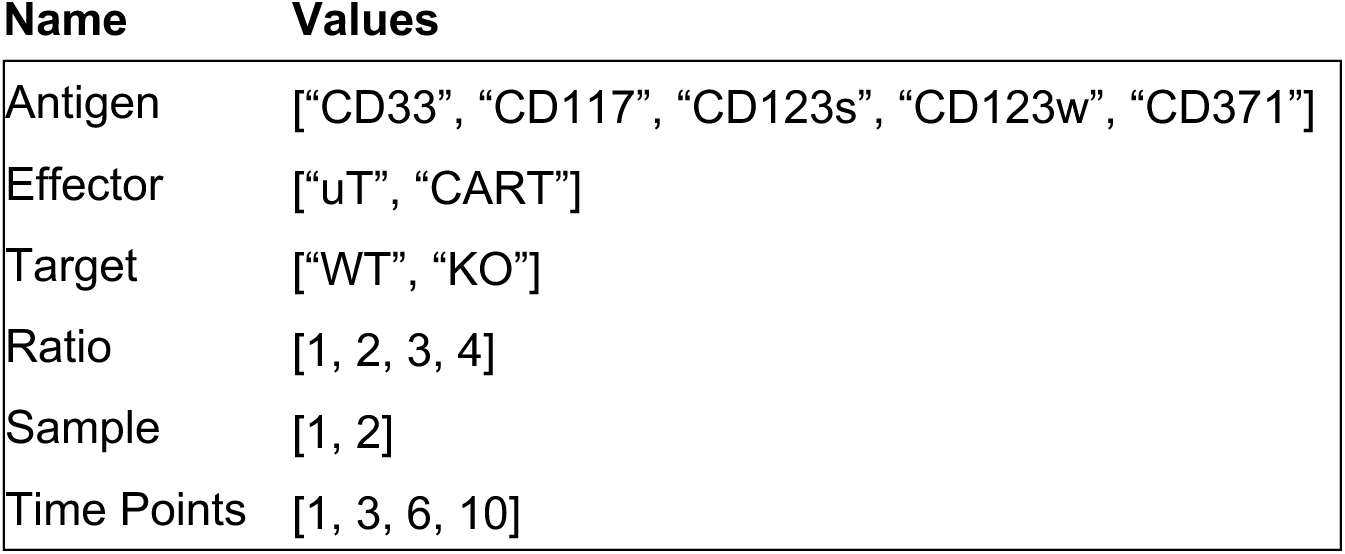
Structure of the experimental data. The first level consists of the five constructs, and the level elements are the names of the target receptors. The second and third levels are the type of effector and target cells, in order, in the co-culture cytotoxic assay. “uT” and “CART” refer to the untransduced and chimeric antigen receptor T-cells. “WT” and “KO” refer to the *TP53*-wildtype and -knockout tumor genotypes. The fourth level specifies the index of initial conditions of experiments (Table 1). The fifth level indicates the index of the biological replication of the co-cultures. The sixth level consists of the time, reported in days, of the cell abundance measurements. The dataset, thus, consists of 5 *×* 2 *×* 2 *×* 4 *×* 2 *×* 4 = 640 measured data points.

### Mathematical modeling of T-cell expansion and killing

We compared the performance of distinct chimeric antigen receptor effector constructs against *TP53*-wildtype and *TP53*-knockout leukemia target cells. To model the dynamics of (target) tumor cells, *x*, and the cytotoxic (effector) T lymphocytes, *y*, we described cancer growth, target cell killing, effector cell expansion, exhaustion, and death. The net rate of change in a cell population’s abundance or density is determined by the target growth function minus the target killing function, resulting in a system of ODEs,

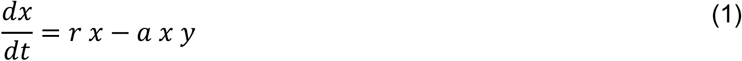

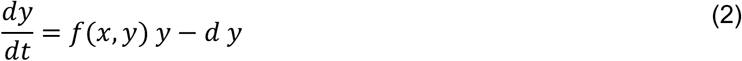

The parameter *r* models the target cell population’s constant per capita growth rate. A mass-action kinetics term describes target cell killing, *a x y*, and effector T-cell death (exhaustion) occurs at a constant rate *d*. Effector T-cell expansion is encoded in the function *f*(*x*, *y*). Equations (1) and (2) were solved numerically in Julia 1.9 using *DiffEqBayes.jl* or in Wolfram Mathematica 14 (*NDSolve*).

### Candidate functional responses for T-cell expansion

We assumed that target cells followed exponential growth and investigated the functional form of effector expansion *f*(*x*, *y*). We also assumed that the target-killing function is unrestricted by any other process, following mass-action kinetics. Effector cells expand in the presence of the target cells and possibly interact, encoded in the function of effector cell expansion *f*(*x*, *y*), **Figure 1B,D**. These expansion laws can be uncoupled, of Lotka-Volterra type ^37,38^, Holling type 2^24,25^, ratio-dependent ^29^, or of a more complicated fractional form^28^, e.g., the Beddington-De Angelis model^26,27^. All model parameters are described and summarized in **Table 3**.

**Table 3:**
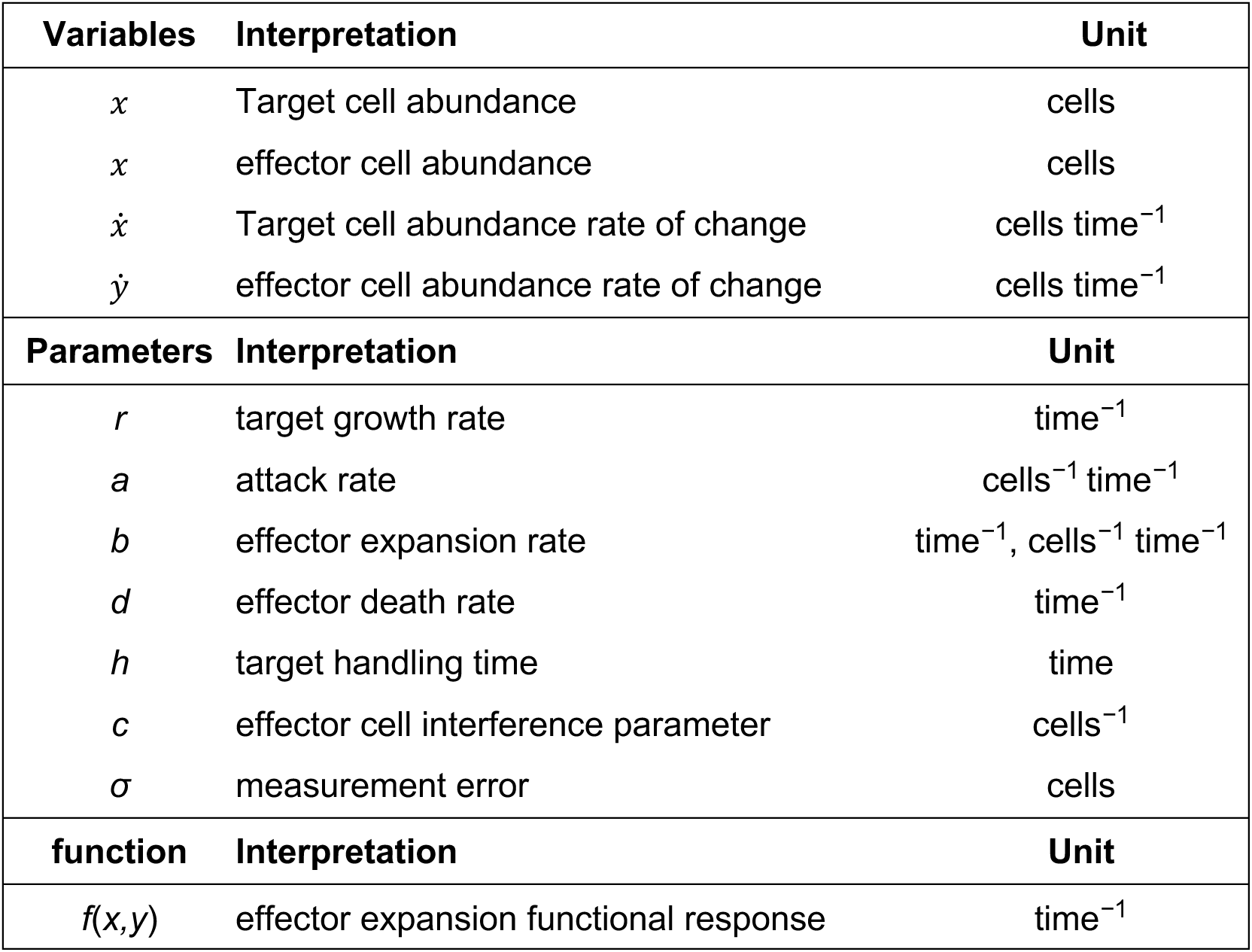
Symbols used in the study with their interpretations and units. We use the shorthand notation *ẋ* = *dx*⁄*dt* for the derivative in time. The measurement error, *σ*, is a non-mechanistic hyperparameter of the Bayesian parameter inference approach (see **Supplementary material**).

| Variables | Interpretation | Unit |
| --- | --- | --- |
| $x$ | Target cell abundance | cells |
| $x$ | effector cell abundance | cells |
| $\dot{x}$ | Target cell abundance rate of change | cells time<sup>-1</sup> |
| $\dot{y}$ | effector cell abundance rate of change | cells time<sup>-1</sup> |
| Parameters | Interpretation | Unit |
| $r$ | target growth rate | time<sup>-1</sup> |
| $a$ | attack rate | cells<sup>-1</sup> time<sup>-1</sup> |
| $b$ | effector expansion rate | time<sup>-1</sup>, cells<sup>-1</sup> time<sup>-1</sup> |
| $d$ | effector death rate | time<sup>-1</sup> |
| $h$ | target handling time | time |
| $c$ | effector cell interference parameter | cells<sup>-1</sup> |
| $\sigma$ | measurement error | cells |
| function | Interpretation | Unit |
| $f(x,y)$ | effector expansion functional response | time<sup>-1</sup> |

### Parameter identifiability, Bayesian inference, and model selection

The first step of parameter inference is assessing whether the parameters can, in principle, be inferred and whether the data is informative for the question at hand. We analyzed *a priori* structural identifiability and *a posteriori* practical identifiability for our candidate models.

Structural identifiability is data-independent. We used the input-output equation method based on the minimal differential polynomial related to the differential equation model ^39^. We used the Julia package *StructuralIdentifiability.jl* to evaluate the a priori identifiability of candidate models.

Practical identifiability can only be evaluated in the context of data. Instead of the profile likelihood approach ^40^, we used pairwise joint posterior distribution analysis ^30^. When the negative log-likelihood or the pairwise joint Bayesian posterior has a peak and forms closed contours in parameter space, the respective parameter combination is uncorrelated and identifiable. When the highest peak region extends infinitely along a particular dimension in parameter space, the parameter cannot be reliably inferred from the data (practical non-identifiability).

Bayesian parameter estimation (see **Supplementary Material**) updates the likelihood of a parameter value by considering prior beliefs and evidence via Bayes’ rule. Based on other experiments, dimensional analysis, and physical constraints, we decided on lower and upper bounds for the parameter values and converted them into Log-Normal priors for each parameter. We assumed a multiplicative measurement error *σ* to derive the corresponding likelihood. We used the Julia package *DiffEqBayes.jl* to estimate the parameters of ordinary differential equation models using Bayesian inference^41^, which provides an interface for *Turing.jl*, a framework for Bayesian inference. We used the No-U-Turn-Sampling variant of the Hamiltonian Monte Carlo algorithm with acceptance probability *δ*=0.65 for inference^42^. Each inference was repeated 5 times, yielding 5 independent chains with 10,000 samples each. We assessed within-chain and across-chain convergence, discarding chains that did not meet the appropriate convergence criteria. The loss from the negative log-likelihood function quantifies a model’s error on the data. We calculated the negative log-likelihood of all the posterior samples to obtain the Akaike Information Criterion and the Bayesian Information Criterion for model selection.

## Results and Discussion

We hypothesized that cell population interactions are key to CAR T expansion^20,22^, and that T-cell activity against resistant AML can be antigen-specific. We were able to select the most appropriate candidate model based on information criteria that account for fitting errors and penalize the number of parameters. Finally, we investigated the performance of CAR T-cells with different constructs and against wildtype or *TP53* knockout leukemia cells by examining the best-performing model’s parameter distributions and time-forward simulations.

### Identifiability, model training, and model selection

We assessed all candidate models for parameter identifiability prior to Bayesian parameter inference (see **Methods**). We found that all parameters in each candidate model are globally identifiable. Note that in candidate models with functional response, the attack rate appears in Equations (1) and (2), which renders the killing process’s handling time identifiable.

We trained the candidate models using an *in vitro* data set (LT15) that measured T or CAR T (always using one of the five constructs) vs. *TP53*-wildtype or *TP53*-knockout AML dynamics up to day 10, to obtain parameter posterior distributions. Model fitting results are shown in **Supplementary Figures 4-9**. We inferred parameter distributions for each CAR construct, effector, target, and initial E:T ratios (**Table 2**), individually pooling over time and experimental replicates. We repeated this inference scheme for all candidate models. To rank candidate models, we summed the negative log-likelihoods over the data frame (**Table 4**). The model selection across candidate models (**Table 4**) preferred the Beddington-DeAngelis model of T and CAR T-cell expansion.

**Table 4:** Differences in corrected Akaike and Bayesian information criteria across candidate models. The independent cytotoxicity assays over five chimeric antigen receptor constructs, four effector-target combinations, and four E:T ratios. Therefore, the parameter log-likelihoods were added. A lower corrected Akaike and Bayesian information criteria score indicates a better model performance. The differences from the lowest criterion value are reported for the candidate models. The Beddington-DeAngelis expansion performs the best, i.e., has the lowest information criterion, amongst candidate models overall, i.e., ignoring CAR constructs, effector-target combination, and the E:T ratio. AICc: *corrected* Akaike information criterion, BIC: Bayesian information criterion. The Δ indicates that values are shown *relative* to the best-performing candidate model. The measurement error, *σ*, is a non-mechanistic hyperparameter of the Bayesian parameter inference approach.

| Candidate model | Number of parameters | $\Delta\text{AICc}$ | $\Delta\text{BIC}$ |
| --- | --- | --- | --- |
| uncoupled, T growth | (5) $x_0, y_0, r, b, \sigma$ | $5.65 \times 10^4$ | $5.65 \times 10^4$ |
| uncoupled, T death | (5) $x_0, y_0, r, d, \sigma$ | $7.54 \times 10^{18}$ | $7.54 \times 10^{18}$ |
| Lotka-Volterra expansion | (7) $x_0, y_0, r, a, b, d, \sigma$ | $3.08 \times 10^2$ | $3.22 \times 10^2$ |
| Lotka-Volterra exhaustion | (7) $x_0, y_0, r, a, b, d, \sigma$ | $6.24 \times 10^{14}$ | $6.24 \times 10^{14}$ |
| Holling type 2 | (8) $x_0, y_0, r, a, d, h, c, \sigma$ | $2.35 \times 10^2$ | $2.44 \times 10^2$ |
| Ratio-dependent expansion | (7) $x_0, y_0, r, a, d, b, \sigma$ | $1.21 \times 10^2$ | $1.35 \times 10^2$ |
| Bazykin-Harrison | (8) $x_0, y_0, r, a, d, b, c, \sigma$ | $6.74 \times 10^5$ | $6.74 \times 10^5$ |
| Beddington-DeAngelis | (9) $x_0, y_0, r, a, d, b, c, h, \sigma$ | 0.0 | 0.0 |

To obtain best-fit equivalents, as exemplified in **Figure 2A-C** and detailed in the **Supplementary Material**, we considered the numerical solution with maximal distance correlation (solid lines). For 95%-confidence prediction intervals (shaded areas), we numerically solved the best-performing model using 250 initial abundances and parameter values randomly sampled from the posterior for each experimental condition, calculated the standard deviation, and multiplied it by the standard factor of 1.96.

**Figure 2:**
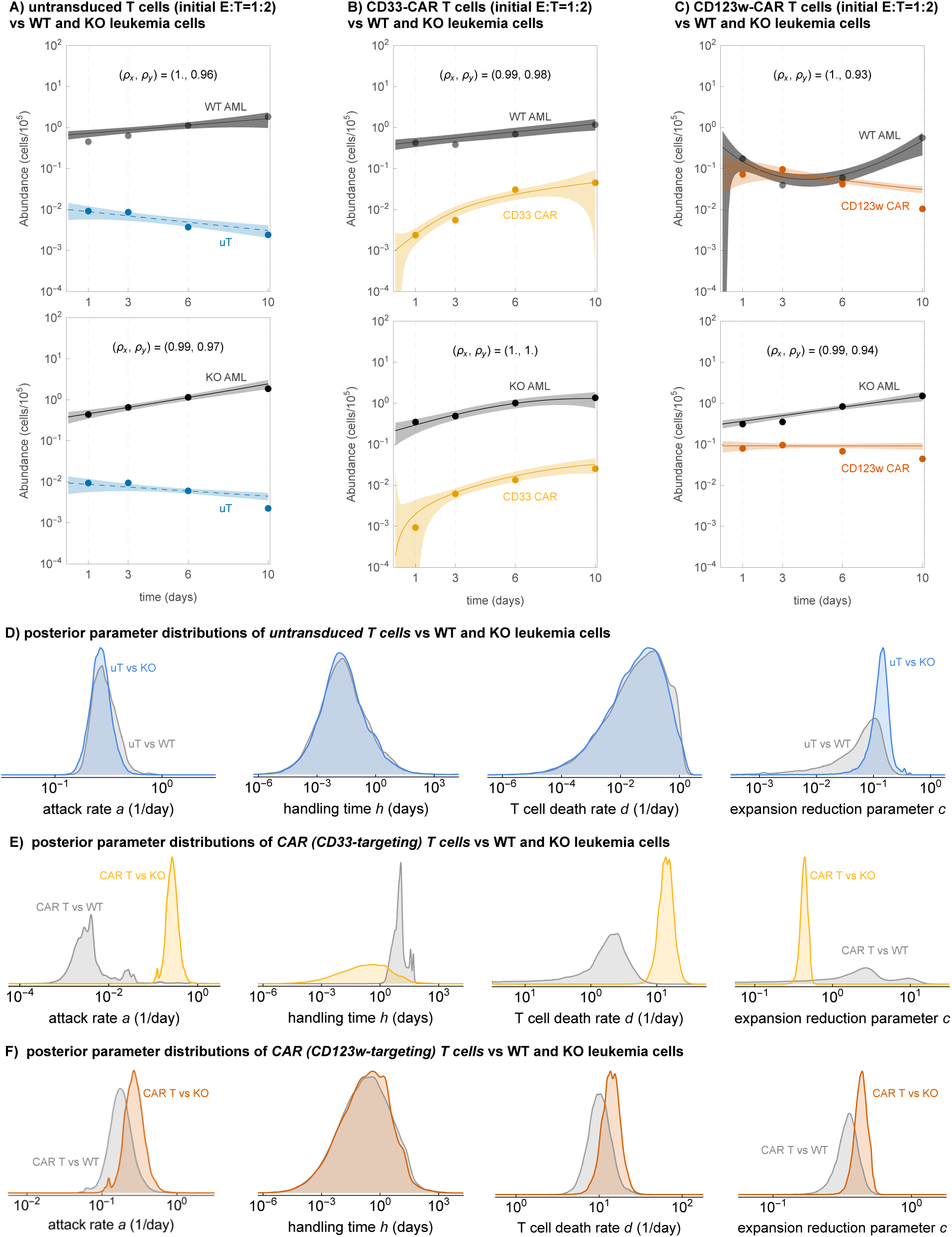
***E*xample Beddington-DeAngelis time series fits and example posterior parameter distributions. A)**-**C)** Example model training results for the best-performing (Beddington-DeAngelis CAR T expansion) model, each comparing the performance of untransduced or CAR T cells in co-culture with either WT or KO AML cells. The the goodness of fit was assessed using custom correlation functions (see **Supplement**) in each modeling compartment, *ρx* and *ρy*, which are also shown in each panel. **D)** Specific posterior parameter distributions of uT-cells vs. WT and KO AML cells. **E)** and **F)** Same posterior parameter distributions for two target antigen CARs (CD33 and CD123w). WT: *TP53*-wildtype target, KO: *TP53*-knockout target, uT: untransduced T-cell, CART: chimeric antigen receptor T-cell.

Specifically, to evaluate trained model performance, we employed a distance correlation statistic between the model and data for target and effector abundances *x* and *y*, *ρ*, and *ρ*_-_, respectively (see **Supplementary Material**). Notably, uT-cells typically performed worse, while the CAR T-cells continued to expand. Meanwhile, KO leukemia cells typically reached higher abundances, indicating decreased cancer control.

Each untransduced or target-antigen setting yielded its own combination of posterior parameter distributions, as every scenario within each CAR T-AML setting contributed independently to model assessment and selection. E.g., CD33 and CD123w, showed different posterior parameter distributions when co-cultured with different AML cells. These distributions were also different compared to untransduced T cell scenarios, as shown in **Figure 2D, E, F**. Thus, a combination of factors works in a complex way to explain the longitudinal co-culture experiments for various target antigens.

Untransduced T cells showed remarkably robust behavior in low attack rates and high death rates across AML subtypes, but differences in the self-inhibitory dynamics (parameter *c*) are already present. In contrast, CD33 showed lower attack rates and elevated death rates. Overall, they did expand better than untransduced cells. Furthermore, CD123w CAR T-cells showed comparable attack and death rates (and handling times) but a lower crowding parameter, which would negatively impact expansion, thus potentially indicating greater efficacy. Taken together, these observations show that our modeling identifies (dynamic) differences in CAR T-cell expansion, lifespan, and attack between constructs.

### CAR T efficacy against wildtype and *TP53*-knockout leukemia

Although variable across target antigen and *TP53*-status conditions (**Supplementary Figures 9**), untransduced T (uT) cells showed little difference when facing wild-type (WT) or *TP53*-knockout (KO) leukemia cells (**Figure 2D**). In contrast, CAR T-cells had an increased attack rate but also an increased death rate (**Figure 2E,F**). CD33 CAR T-cells showed a more favorable attack rate, a reduced handling time, and less T-cell crowding (parameter *c*), but had a markedly increased death rate (**Figure 2E**). CD123w CAR Ts showed less increase in attack rate, unchanged handling time, slight change in death rate, but more T-cell crowding (parameter *c*, **Figure 2F**). These differences highlight target antigen-specific mechanisms of CAR T evasion in *TP53*-knockout leukemia. These findings indicate how the target antigen shapes the complex pharmacokinetics against leukemia.

### Dynamic features of CAR T expansion

The Beddington-DeAngelis model assumes that T-cell expansion depends on both target cell and effector abundance; expansion decreases with the number of target cells due to the fraction of T-cells engaged in the killing (handling time) and with the number of other T-cells due to possible crowding and respective saturation. Interestingly, the ratio-dependent T-cell expansion model also has this property, but with less nuance; it performed second-best (**Table 4**). A common feature of both models is that effector expansion depends on both target and effector abundances, increasing with target abundance but decreasing with effector abundance.

It is important to note that corrected AIC and BIC both penalize for the number of parameters. Yet the Beddington-DeAngelis expansion, the model with the highest number of parameters, delivered the best scores across all conditions.

### Model validation

To validate the best-performing model, we used a separate set of longitudinal data of the same cell combinations, different initial E:T ratios, and compared model trajectories generated from 250 random samples of the parameter posteriors to the data. This second data set (LT6) went up to day 16 and was not used for model training or selection. To evaluate model performance on this unseen data, we again calculated the distance correlations *ρ*, and *ρ*_-_. Distance correlations above 0.6 are considered decent, and above 0.9 indicate excellent statistical agreement between two samples (e.g., between predictions from trained parameter values and validation data). A schematic of our true out-of-sample validation test, using a previously unseen data set that followed the cell populations for a longer period, is shown in **Figure 3A**. The shaded bands in **Figure 3B** and **C** are 95% posterior predictive intervals obtained by propagating 250 joint draws of parameters from the trained model, reflecting the combined uncertainty in initial conditions, kinetic parameters, and multiplicative measurement error. **Figure 3D** shows the distance correlation values across multiple experimental scenarios. We sometimes observed a drastic widening of the 95% prediction interval (**Figures 3B,C)**. Yet, all model-data correlation values were above 0.6, suggesting that the Beddington-DeAngelis expansion model can be used to predict CAR T-leukemia kinetics.

**Figure 3:**
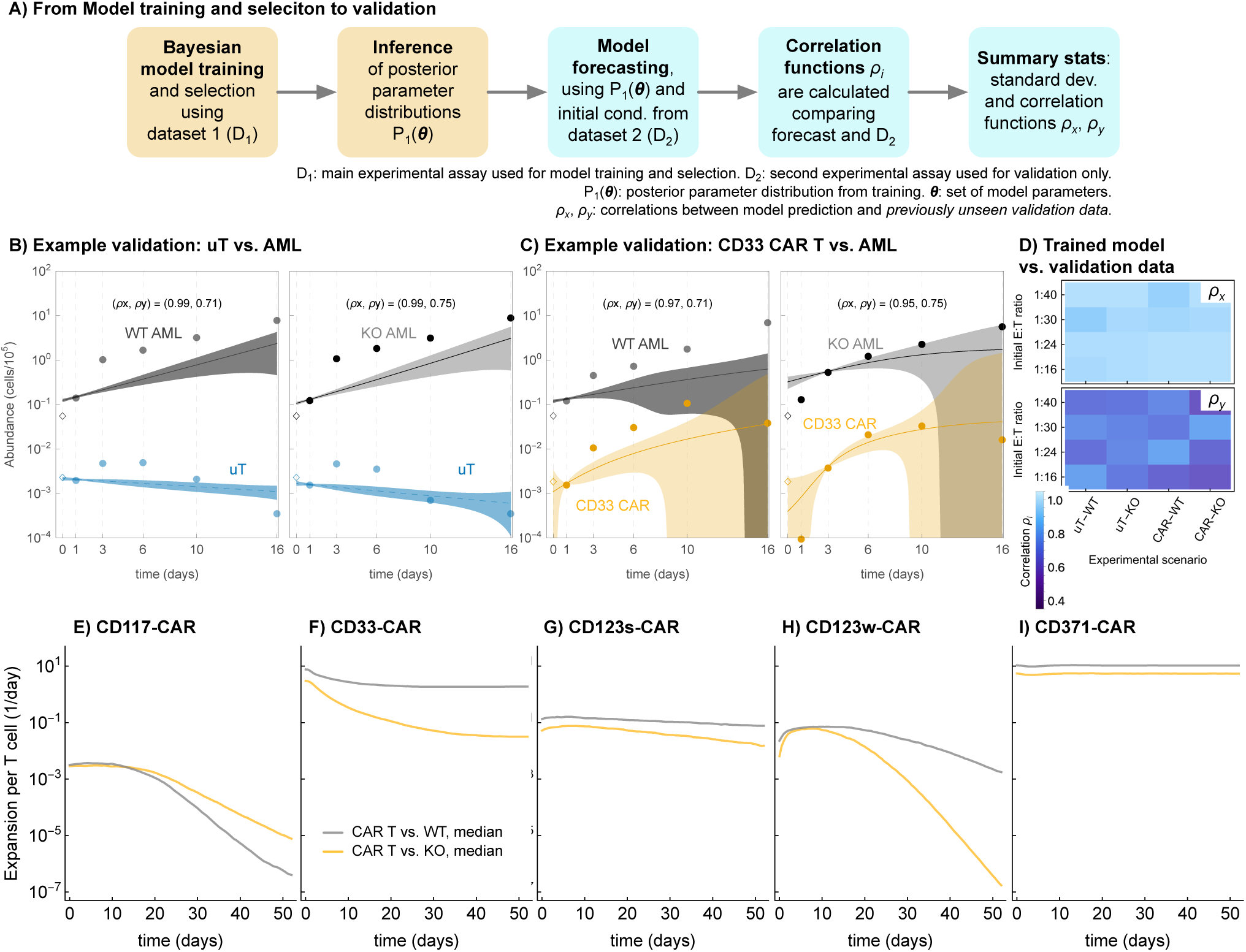
Validation of the best-performing model using initial conditions of previously unseen data. **A)** Schematic of the validation approach using two different assay experiments (dataset 1, dataset 2, see Methods). **B)**,**C)**. Example plots in which the model was parameterized using the posteriors from model training. The bands indicate the 95% posterior predictive interval, calculated from 1.96 times the standard deviation of 250 trajectories generated with randomly sampled parameter values from the posterior distributions. D) For goodness-of-fit of the validation, we calculated the distance correlations (see **Supplementary Material**) between prediction and validation, which were typically higher for the leukemia compartment *x* than for the (CAR) T-cell compartment *y.* **E)-I)** Median CAR T-cell expansion rates, *f*(*x*,*y*), over time, for the different constructs, sampled over 5000 values from the posterior parameter distributions of the best-performing validated expansion model. WT: *TP53*-wildtype target, KO: *TP53*-knockout target, uT: untransduced T-cell, CART: chimeric antigen receptor T-cell.

### An effector-target interaction model reliably captures T-cell performance

The posterior parameter distributions were varied across experimental conditions. **Supplementary Figures 9 (A-E)** shows the target antigen-specific posterior parameter distributions for the same four key parameters, comparing the performance of the differently constructed CAR T-cells with WT and KO leukemia cells. E:T ratios of ≈ 1 show prominent target killing with rapidly decreasing target abundance at first, followed by recovery, while the effector abundance constantly decreases. For E:T ratios ≈ 0.1, we observed little to no target killing, leading to target growth. Notably, CD123w-targeting chimeric antigen receptor T-cells exhibited more killing against *TP53* wildtype target cells than against the *TP53* knockout target while maintaining their handling time (see also **Figure 2E,F**). For target antigens CD117 and CD123s, the growth rate of the T-cells decreased with a decreasing initial E:T ratio. For CD33 and CD371, this trend was not apparent. While the Bayesian inference scheme selected the best model of CAR T-cell expansion, unresolved variation indicates underlying heterogeneity in both CAR T and target cell phenotypes.

### *TP53*-deficiency confers resistance, but in a target antigen-dependent manner

The preferred model of T- and CAR T-cell expansion implies a time dependence. Depending on the initial E:T ratios, the expansion rate is dynamic, as it depends on cell population feedback through the mechanisms of handling time required for killing (parameters *a* and *h*) and crowding due to T-cell competition (parameter *c*). To assess this complex process, we evaluated the Beddington-DeAngelis over time using 5000 parameter combinations drawn from the posterior distributions across the five target antigens and solving the coupled dynamical system; **Figure 3**, panels **E-I** show the expansion rate function *f*(*x*,*y*) in equation (2), per cell (medians taken over 5000 simulations of the system dynamics with random sampling from the posterior parameter distributions). Stark differences arise in how CAR T-cells expand vs. WT or *TP53* KO AML cells. All constructs except CD117 exhibit lower overall dynamic expansion rates per CAR T cell. For CD117 CARs, overall expansion is very low, as expected^8^. For CD33 and CD371, the expansion rate stabilized over the long term. These differences suggest that *TP53* knockout evasion occurs in a multifaceted, target-antigen-dependent fashion.

### Target-antigen-specific CAR T properties

The *in vitro* CAR T-cell response is typically short-lived compared to *in vivo* and potential clinical responses. The best-performing candidate model (**Figure 4A,B**) captures the duration of CAR T-target cell conjugation (handling time) and CAR T-cell crowding as possible detriments to expansion. The candidate models do not capture expansion through differentiation, e.g., by adding compartments of stem, naïve, or memory T-cells^22^ as we found no evidence that this added complexity contributed to the *in vitro* cytotoxicity assays.

**Figure 4:**
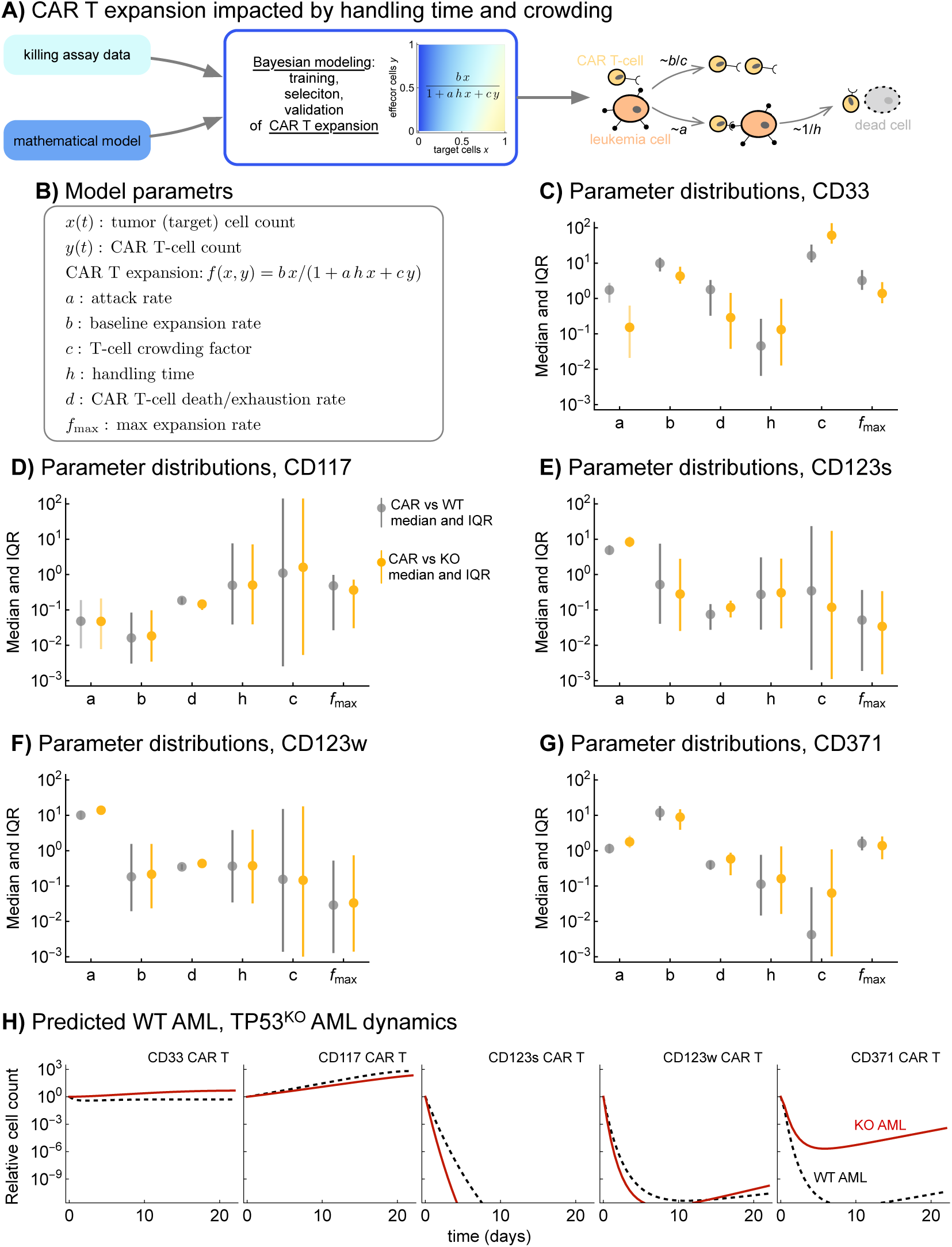
**Target antigen-specific CAR T kinetics**. **A)** Graphical summary of the main approach and results. **B)** Summary of the variables and parameters that determine the multi-parametric nonlinear functional form of CAR T expansion. In addition to the static parameters that affect the expansion function *f*(*x*,*y*), we also determined the maximum expansion over time by simulating the model using 5000 randomly sampled parameter combinations drawn from the validation model’s posteriors. **C)**-**G)** Parameter distributions (medians and interquartile range, IQR) of the five target antigens, comparing CAR T vs. WT (gray) and CAR T vs. KO (yellow). **H)** We used the median posterior parameter values to simulate the dynamical system *x*(*t*), *y*(*t*), and compare each CAR’s performance against WT (black) and KO AML (red). Only the CD123s targeting CAR T-cells are predicted to suppress KO AML. Chosen initial effector-to-target ratio=1. WT: *TP53*-wildtype target, KO: *TP53*-knockout target.

We analyzed the system’s two possible fixed points using linear stability analysis (**Supplementary Material**). In the case of tumor control, the stable target (leukemia) population is given by *d*⁄*a* × (*a* + *cr*)⁄(*b* − *a d* ℎ), and the steady-state of the effector population is *r*⁄*a*. This state is unstable if the attack rate meets the condition *a* > *b*⁄(*d* ℎ). Tumors could escape therapy pressure for long handling times, *h*, or high effector death rate, *d*. This calculation exemplifies the complex impact of the best-performing model’s parameters. Interestingly, a very high attack rate, unbalanced by a low handling time, can lead to tumor escape and possible progression because CAR T cells that constantly attack and form CAR T-target cell conjugates cannot contribute to expansion.

We hypothesized that CAR T antigen-targets behave differently when challenged with *TP53*-knockout cells. To this end, we analyzed the posterior parameter distributions of the best-performing model. Our primary interest was comparing the CAR vs. WT and CAR vs. KO scenarios, as these could indicate specific differences in how a particular CAR T construct handles the *TP53* KO challenge.

In **Figure 4C-G**, we compare the posterior median values of the CAR vs. WT and CAR vs. KO scenarios (full distributions shown in **Supplementary Figure 10**), focusing on the parameters important for killing (attack rate) and expansion (baseline expansion, handling time, crowding, T-cell death rate), as well as the derived quantity of maximal CAR expansion within the first 55 days.

CD33 showed a lower attack rate and baseline expansion rate, as well as a lower CAR T death rate (**Figure 4C**). CD117 showed relatively conserved parameter values, with a low overall attack rate (**Figure 4D**). CD123s, CD123w, and CD371 all showed increased attack rates against the KO cells, at the cost of reduced baseline expansion (CD123s) or increased handling time (CD371) (**Figure 4E,F,G**). Together, these observations underscore how our best-performing nonlinear model of CAR T cell expansion can capture a multifaceted, target-antigen-dependent spectrum of changes in kinetic parameters that determine efficacy.

The modeled dynamics of WT and KO AML, based on equations (1) and (2) and the median posterior parameter values, confirmed these observations (**Figure 4H**). CD117 was ineffective against the cancer cell population. CD33 had a moderate effect, while CD123s likely suppressed both WT and KO AML for extended times. In comparison, both CD123w and CD371 eventually lost their efficacy against the KO AML population.

These comparisons of parameter distributions and their impact on the target AML dynamics highlight how different sets of parameter values can produce similar overall kinetics. The underlying functional mechanisms of CAR T evasion can be quite different, such as a reduced attack rate or increased handling time, leading to lower killing efficacy or decreased CAR T cell expansion. In all but one target (CD33), the CAR T cells sought to attack more, but this effect was mainly muted by either an increased handling time (time spent killing rather than expanding) or decreased CAR T survival. CD123s’ relative success against both WT and KO AML may stem from their shorter handling time, thereby mitigating the negative effect of the target-killing process on CAR expansion.

## Conclusions

Computational and mathematical models can complement experimental approaches to explore mechanisms driving tumor-immune interactions, particularly CAR T-target cell kinetics. We quantified and compared the performance of untransduced T-cells and CAR T-cells expressing five different antigens against two leukemia genotypes. We inferred individual system-specific parameter distributions using a novel Bayesian estimation workflow. As a result, we inferred the functional relationship between cellular densities and CAR T-cell expansion. We evaluated how dynamic expansion, increased attack rate, and decreased CAR T-cell lifespan interact, highlighting target-antigen-specific complications in treating *TP53*-deficient leukemia cells. This analysis enabled the quantification of the dose dependence and the long-term effects on effector T-cell performance. By tuning CAR T performance parameters and dosing strategies, it is possible to optimize treatment outcomes based on theoretical considerations. Expansion, rather than killing, may be the ‘rate-limiting’ step in optimizing CAR T-cell therapy against AML.

We used a Bayesian approach for practical identifiability analysis and model selection. Other model selection procedures based on practical identifiability use the profile likelihood^21,40^. These methods rely on context-adapted optimization algorithms and are vulnerable to local optima. Bayesian inference replaces this optimization with parameter sampling via a Hamiltonian Monte Carlo approach^42^. We estimated posterior parameter distributions in this way and quantified uncertainty using credibility intervals^43^. While many other statistical methods require large sample sizes, the minimum sample size for Bayesian inference is one (given perfect priors)^44^: the Bayesian approach requires prior knowledge. We used knowledge of tumor growth rates from independent cancer growth experiments to eliminate regions of parameter space that are *a priori* unlikely. Experimental insights into the timescale of the relevant biological processes can similarly be incorporated.

Our study has several limitations. First, the experimental setting cannot account for clinical heterogeneity among patients and antigen diversity. Second, we do not present an analysis of antigen mixtures, antigen density, or the evolution of selection of *TP53*-depleted leukemia variants. These aspects might be crucial ingredients for future studies. Third, we did not account for antigen and CAR abundance per cell, as well as the potentially varying affinities of different CAR constructs. These variabilities are reflected in differences in posterior parameter estimates across constructs, particularly in baseline expansion and attack rate. Of note, observed differences in these dynamics may not be driven primarily by differential antigen availability on the target cell^8^, and we were only able to use previously unseen *in vitro* data of the same AML cell line for validation. Fourth, a fully mechanistic receptor-ligand description is beyond the scope of the current study, but could be the next step for specifically valuable CAR T constructs. Density-dependent antigen modulation was not measured in the current study, but antigen expression profiles were previously characterized^8^.

Specific investigations that combine antigen density with mathematical system characterizations of CAR T expansions are valuable avenues for future work; antigen density should be explicitly modeled as a state variable rather than implicitly influencing the inferred parameter distributions.

As reported in several clinical studies, cancer-T-cell dynamics and CAR T therapy outcomes depend on the achieved E:T ratio^45–48^. In particular, the CD123- and CD371-targeting CARs exhibited strong cytotoxicity, even against mutant AML. Elevated effector death rates often accompanied these high attack rate estimates, suggesting higher CAR T exhaustion and cell turnover. These findings imply that attack rate and effector turnover may be essential for eliminating the target leukemia, as CAR T expansion is limited by the time spent attacking (handling time) and by self-interference. Due to the handling time, the attack rate has a positive effect from immediate cancer killing and a negative impact on expansion because the killing process binds CAR T cells.

As recently shown and discussed, the pharmacodynamic process of CAR T-target cell conjugation can delay expansion^49^ and may impede efficacy^50^. Short handling times could lead to sustained CAR T-cell abundance, a form of persistence. Our results suggest that sustained effector expansion with reduced handling time could be a crucial CAR T-cell characteristic that translates into durable responses, but that different target antigens offer specific kinetic benefits. Future studies should leverage these complementary properties against AML and, if feasible, combine multiple target-antigen CAR T constructs into a single dose or exploit them for subsequent dosing.

## Data availability

The data and code of this project are publicly available through a zenodo repository 10.5281/zenodo.12823159.

## Ethics approval

Ethics approval was not required, as this study does not use data from human or animal subjects.

## Supplementary Material

The Supplementary Material can be found on 10.5281/zenodo.12823159.

## Acknowledgments

SaSh, MR, AT, and PMA acknowledge generous funding from the Max Planck Society. SaSh, MR, and AT acknowledge funding by Deutsche Forschungsgemeinschaft through the Research Training Group “Translational Evolutionary Research” (TransEvo, project number 400993799). MGM was supported by the Clinical Research Priority Program “ImmunoCure” of the University of Zurich. SB was supported by the Promedica Foundation Chur, the Swiss Cancer League (KFS-4885-08-2019), the Fondation Peter Anton & Anna Katharina Miescher pour la Recherche en Hématologie, and the Swiss Society of Hematology. JM was supported by the KRAK-Physician Scientist Fellowship. PMA is supported by the DFG Heisenberg Program (project number 525136051) and the DFG Clinical Research Unit 5010 CATCH ALL (project number 444949889).

## Author contributions

SaSh, JM, and PMA conceived the study and wrote the main manuscript text. JM and EV performed experiments. SaSh, JM, EV, MR, and PMA analyzed the data. SaSh, EV, and PMA performed statistical analyses, prepared the figures, and wrote the Supplement. AT, SB, MGM, and PMA supervised the study. All authors contributed to the final version and reviewed the manuscript.

