## Supplementary Methods, Tables, and Figures for "Quantifying target antigen-dependent CAR T-cell performance against AML"

—

June 12, 2026

#### Contents

**Bayesian inference and the statistical model** 2

**Model selection** 3

**Linear stability of Beddington-DeAngelis expansion model** 4

**Long-term effects of effector properties** 4

**Distance correlation** 5

**Negative log-likelihoods for CAR-model pair** 5

**References** 7

**Supplementary tables** 8

**Supplementary figures** 11

### Bayesian inference and the statistical model

Bayesian parameter estimation exploits the observation that the joint probability of two events is related to the likelihood of one of the events when the other has already happened Bayes and Price (1997); Ellison (1996). Consider the two events $\boldsymbol{\theta}$, and $\boldsymbol{y}$, the probability that both events occur equals the product of probability of one of the events and the conditional probability of the second event given the first, i.e., $\Pr(\boldsymbol{y})\Pr(\boldsymbol{\theta} \mid \boldsymbol{y}) = \Pr(\boldsymbol{y}, \boldsymbol{\theta}) = \Pr(\boldsymbol{\theta})\Pr(\boldsymbol{y} \mid \boldsymbol{\theta})$. Rearranging this expression using the two terminal terms, we get

$$\Pr(\boldsymbol{\theta} \mid \boldsymbol{y}) = \frac{\Pr(\boldsymbol{\theta})\Pr(\boldsymbol{y} \mid \boldsymbol{\theta})}{\Pr(\boldsymbol{y})}. \tag{S.1}$$

Here, $\boldsymbol{\theta}$ is the set of parameters, $\boldsymbol{y}$ are observations, and $\Pr(\boldsymbol{\theta})$ is the prior probability of parameters, i.e., investigator's expectation of how likely a value is. This prior probability can be informed from other experiments. The term $\Pr(\boldsymbol{x} \mid \boldsymbol{\theta})$ is known as the likelihood function for the parameter, and $\Pr(\boldsymbol{y})$ is the expected value of the likelihood function also known as 'evidence.' The Bayes theorem describes the posterior probability as proportional to the product of the likelihood and the prior probability, indicating that likelihood modifies prior expectations into the posterior. The ratio of the posterior probability of two alternate parameter sets $\boldsymbol{\theta}_1$, and $\boldsymbol{\theta}_2$ are called the odds ratio

$$\frac{\Pr(\boldsymbol{\theta}_2 \mid \boldsymbol{y})}{\Pr(\boldsymbol{\theta}_1 \mid \boldsymbol{y})} = \frac{\Pr(\boldsymbol{\theta}_2)\Pr(\boldsymbol{y} \mid \boldsymbol{\theta}_2)}{\Pr(\boldsymbol{\theta}_1)\Pr(\boldsymbol{y} \mid \boldsymbol{\theta}_1)}. \tag{S.2}$$

The odds ratio quantifies the relative expectations of parameter set $\boldsymbol{\theta}_2$ compared to $\boldsymbol{\theta}_1$. Hoffman and Gelman (2011) developed a class of Hamiltonian Monte Carlo algorithms, more efficient than randomly guessing, to find the posterior distribution using the odds ratio of parameter sets. An independent sequence of such parameter values is called a (Markov) chain, and the elements of this sequence are distributed identically to the posterior distribution.

Using this approach, prior distributions, and likelihoods, we obtain probability distributions over parameters for our model and data. Based on other experiments, dimensional analysis, and physical constraints, we decided lower bounds and upper bounds on the parameter values. The bounds for each parameter were turned into Log Normal priors for each parameter.

We considered measurement $\sigma$ error between the generative model and the experimental measurements. This measurement error $\sigma$ is multiplicative, i.e., the standard deviation of the replicate measurements is proportional to the abundances, and the corresponding likelihood is derived in the following section. There are different ways to combine the available data using statistical hierarchy. Complete pooling combines all the levels of the data frame (**Table 2**) to infer the parameters. The underlying assumption is that a unique set of parameters can reproduce all observed data. We assume that chimeric antigen receptor constructs, types of effectors, and types of target cells have distinct best-fit parameter sets, which we can use to compare their parameters. We also suspect that the tumor-immune dynamics are dose-dependent and differ at different T cell abundances. Thus, we do not pool over the following levels: Antigen, Effector, Target, and Ratios. Samples and time levels share a set of parameters; thus, they have parameter pooling. So, our hierarchy leads to a statistical model with partial pooling across dataset levels.

We use `DiffEqBayes.jl`, a Julia package, to estimate the parameters of the ordinary-differential-equation models using Bayesian inference (Dong et al., 2023). The package provides an interface for `Turing.jl`, a framework for Bayesian inference (Ge et al., 2018). The 5 Antigens $\times$ 2 Effectors $\times$ 2 Target $\times$ 4 initial effector-to-target ratios result in 80 unique inferences. We use the No-U-Turn-Sampling variant of the Hamiltonian Monte Carlo algorithm with acceptance probability $\delta = 0.65$ for each of the inferences. Each inference is repeated five times, yielding five independent chains with 10,000 samples each. To assess the reliability of sampled chains, we check the within-chain and across-chain convergence. Convergence of each chain is evaluated using the potential scale reduction factor $\hat{R}$ (Gelman and Rubin, 1992; Gelman et al., 2014). When $\hat{R} \approx 1$, we conclude the chains converged; and we discard the chains with $|\hat{R} - 1| > 0.03$.

The pairwise joint posterior distributions of 80 inferences are presented below; please see the individual figure title and Supplementary Fig. 11 for a joint description.

### Model selection

We infer the parameters of all identifiable candidate models and compare the performance of each model using the corrected Akaike information and Bayesian information criteria. The model comparison assesses how well a candidate model captures the data and how reasonable the corresponding assumptions are. The error of a model is quantified by the loss given by the negative log-likelihood function with the data as the reference. To derive the log-likelihood function, we begin with the assumption that the observed abundance $y_{i,t}$ is normally distributed around the true abundance $\tilde{y}_{i,t}$ of cell type $i$ at time $t$ with standard deviation $\sigma\ \tilde{y}_{i,t}$.

$$\begin{aligned} &y_{i,t} \sim \mathcal{N}(\tilde{y}_{i,t}, \sigma \tilde{y}_{i,t}), \\ &\Pr(y_{i,t}) = (2\pi)^{-\frac{1}{2}} \left(\sigma\ \tilde{y}_{i,t}\right)^{-1} \exp\left[\frac{1}{2}\left(\frac{y_{i,t} - \tilde{y}_{i,t}}{\sigma\ \tilde{y}_{i,t}}\right)^2\right]. \end{aligned} \tag{S.3}$$

With the assumption of independent and identically distributed observations, the likelihood is

$$\begin{aligned} \Pr(\boldsymbol{y}) &= \prod_{i,t}^{n,m} \left(2\pi\sigma^2\right)^{-\frac{1}{2}} \left(\tilde{y}_{i,t}\right)^{-1} \exp\left[\frac{1}{2}\left(\frac{y_{i,t} - \tilde{y}_{i,t}}{\sigma\ \tilde{y}_{i,t}}\right)^2\right]. \\ \Pr(\boldsymbol{y}) &= \left(2\pi\sigma^2\right)^{-\frac{n\cdot m}{2}} \left[\prod_{i,t}^{n,m} \left(\tilde{y}_{i,t}\right)^{-1}\right] \exp\left[\frac{1}{2}\sum_{i,t}^{n,m}\left(\frac{y_{i,t} - \tilde{y}_{i,t}}{\sigma\ \tilde{y}_{i,t}}\right)^2\right]. \\ \log \mathcal{L}(\hat{\boldsymbol{\theta}} \mid \boldsymbol{y}) = \log \Pr(\boldsymbol{y}) &= -\frac{n \cdot m}{2} \log\left(2\pi\sigma^2\right) - \sum_{i,t}^{n,m} \log\left(\tilde{y}_{i,t}\right) + \frac{1}{2}\sum_{i,t}^{n,m}\left(\frac{y_{i,t} - \tilde{y}_{i,t}}{\sigma\ \tilde{y}_{i,t}}\right)^2. \end{aligned} \tag{S.4}$$

Here, the quantity $\log \mathcal{L}(\hat{\boldsymbol{\theta}} \mid \boldsymbol{y})$ is the log likelihood for estimated set of parameters $\hat{\boldsymbol{\theta}}$ given the dataset $\boldsymbol{y}$. The last term of the negative log-likelihood is similar to the sum of squared error; analogously, the negative log-likelihood is an error function displaced by a constant value. We calculated the negative log-likelihood of all the posterior samples for model selection and used the median parameter value for further calculations. The corrected Akaike and Bayesian information criteria consider the negative log of the likelihood evaluated at the parameter value and penalize the number of parameters (Gelman et al., 2014). We use the following definition for information criteria:

$$\mathrm{AICc} = 2\kappa - 2\log(\mathcal{L}) + \frac{2\kappa(\kappa+1)}{\eta - (\kappa+1)}, \tag{S.5}$$

$$\mathrm{BIC} = \kappa \log(\eta) - 2\log(\mathcal{L}). \tag{S.6}$$

Here, $\kappa$ is the total number of unknowns, and $\eta$ is the number of samples used for fitting. The number of samples were fixed to the product of 4 time points, 2 observable, 2 samples, i.e. $\eta = 4 \times 2 \times 2 = 16$. The unknowns in each model are listed in the **Table 4**.

### Linear stability of Beddington-DeAngelis expansion model

We examine the linear stability of the candidate model with Beddington-DeAngelis expansion. The set of differential equation for this model is

$$\begin{aligned} \dot{x} &= r\ x - a\ x\ y \\ \dot{y} &= \frac{b\ x\ y}{1 + a\ h\ x + c\ y} - d\ y. \end{aligned} \tag{S.7}$$

At the equilibrium the target and effector cell abundances do not change, i.e. $\dot{x} = \dot{y} = 0$. Substituting this in the model, we get

$$\begin{aligned} 0 &= r\ x - a\ x\ y \\ 0 &= \frac{b\ x\ y}{1 + a\ h\ x + c\ y} - d\ y \end{aligned} \tag{S.8}$$

Note that $(x, y) = (0, 0)$ is a solution of this set of equations. Dividing the first equation by $x$, and the second equation by $y$, we get

$$\begin{aligned} y &= \frac{r}{a} \\ x &= \frac{d\ (1 + c\ y)}{b - a\ d\ h} \end{aligned} \tag{S.9}$$

Substituting the value of $y$ from first equation into the second, we obtain the second set of fixed points for the system.

$$(x, y) = \left( \frac{d\ (a + c\ r)}{a\ (b - a\ d\ h)}, \frac{r}{a} \right).$$

The eigenvalues of the Jacobian matrix evaluated at the 2nd equilibrium are a pair of imaginary numbers with real part $-\frac{r\ d\ c\ (b-a\ d\ h)}{2\ b\ (a-c\ r)}$. Since all the parameter values are positive, $\frac{b-a\ d\ h}{a-c\ r} > 0$ ensures that the equilibrium is stable.

### Long-term effects of effector properties

We examined the stability of the model with Beddington-DeAngelis expansion using linear stability analysis (see Supplementary Section Linear stability of Beddington-DeAngelis expansion model). To investigate how the performance of the effector cells affects the treatment outcome, we look at long-term target abundance as a function of the model parameters. The long-term target cell abundance changes proportional to the inverse of the expansion rate $b$ and the negative inverse of the handling time $h$. Thus, increasing the expansion rate $b$ improves the treatment outcome by decreasing the long-term target cell abundance. Conversely, the treatment outcome is improved by decreasing the handling time $h$ The long-term target cell abundance is proportional to effector interference $c$. Therefore, as the effector interference increases, the long-term target cell worsens the treatment outcome. The long-term target cell abundance depends non-linearly on the attack rate $a$. Thus, an intermediate attack rate $a \approx 1$ minimizes the long-term target cell abundance. The handling time determines the optimal range of attack rate, i.e., lower handling time leads to a broader range of optimal attack rate.

### Distance correlation

The distance correlation is a relatively new measure of joint (in)dependence between random samples. The distance correlation $\rho = 0$ if and only if the two samples are independent. Otherwise, the distance correlation $\rho$ is a positive real number between $0 < \rho \leq 1$. Let $(p_k, q_k), k = 1, 2, \ldots, n$ be a pair of real-valued samples. We compute $n \times n$ distance matrix containing pairwise distances between each $p_k$ (or each $q_k$) using

$$a_{j,k} = \parallel p_j - p_k \parallel, \quad j,k = 1,2,\ldots,n b_{j,k} \quad = \parallel q_j - q_k \parallel, \quad j,k = 1,2,\ldots,n, \tag{S.10}$$

where $\parallel \cdot \parallel$ denotes Euclidean norm. We, then, compute all doubly centered distances using

$$A_{j,k} = a_{j,k} - \bar{a_j} - \bar{a_k} + \bar{\bar{a}} B_{j,k} \quad = b_{j,k} - \bar{b_j} - \bar{b_k} + \bar{\bar{b}}, \tag{S.11}$$

where, $\bar{a}_j$, and $\bar{a}_k$ are $j$-th row and $k$-th column means of the distance matrix $a_{j,k}$. The $\bar{\bar{a}}$ is the mean of all pairwise distances in the distance matrix $a_{j,k}$. A similar notation is applied for the computed values from sample $q$. The squared sample distance covariance is then a scalar given by the mean of the element-wise matrix product of doubly centered distances

$$\mathrm{dCov}_n^2(p,q) = \frac{1}{n^2} \sum_{j=1}^{n} \sum_{k=1}^{n} A_{j,k} B_{j,k}. \tag{S.12}$$

Similar to ordinary variance, the distance variance is defined as the distance covariance of a sample with itself $\mathrm{dVar}^2(p) = \mathrm{dCov}^2(p,p)$. The distance correlation $\rho(p,q)$ of two samples $p$ and $q$ is obtained by dividing their distance covariance by the product of the distance standard deviations $\rho(p,q) = \mathrm{dCor}^2(p,q) = \frac{\mathrm{dCov}^2(p,q)}{\sqrt{\mathrm{dVar}^2(p)\ \mathrm{dVar}^2(q)}}$ Székely et al. (2007); Székely and Rizzo (2009). We used `EnergyStatistics.jl` package available for julia programming language that features an implementation of distance correlation.

### Negative log-likelihoods for CAR-model pair

The corrected Akaike and Bayesian information criteria can be calculated from the negative log-likelihood (see Supplementary Eqs. S.5 and S.6 in Methods section). These negative log-likelihood values reveal how well each candidate model fits to a specific chimeric antigen receptor co-culture data (Supplementary Table 4). Uncoupled T cell growth and uncoupled T cell death do not fit well in every case. This could be due to the fact in no case T cell monotonically grow or die exponentially. Lotka-Volterra exhaustion is a bad fit consistently for every chimeric-antigen-receptor construct. However, Lotka-Volterra expansion has a consistently good fit for all CAR T cell constructs. This observation, coupled with the previous observation, signifies that T cell abundances change non-monotonically and nonlinearly. The cytotoxic effector function is either low or occurs instantaneously when Holling type 2 expansion model shows a bad fit. The Bazykin expansion features T cell interference when this is not an appropriate assumption for the underlying activity, this model fails. Consequently, the models that consistently feature saturating effects in both target and effector abundance perform better than those without this feature. The Beddington-DeAngelis model features saturating effects in target and effector abundance tunable via two parameters. The improvement in the fit, i.e., the reduction in error, of the Beddington-DeAngelis expansion model is higher than the penalty of these parameters, resulting in the best performance (see **Table 4**).

### Supplementary tables

| Antibody | Dilution | Supplier |
|---|---|---|
| CD3(OKT3)-PE | 1:200 | BioLegend® |
| CD34(QBEND10)-FITC | 1:30 | ThermoFischer |
| CD33(WM53)-BV711 | 1:300 | BioLegend® |
| CD123(6H6)-APC | 1:400 | BioLegend® |
| Hoechst | 1:5000 | ThermoFischer |

**Supplementary Table 1:** Reagents and antibodies used for flow cytometry-based analysis of co-incubation assays in this study.

| **CAR scFv target** | **E:T 1** | **E:T 2** | **E:T 3** | **E:T 4** |
|---|---|---|---|---|
| CD33 | 1:16 | 1:24 | 1:30 | 1:40 |
| CD117 | 2:1 | 1:1 | 1:2 | 1:8 |
| CD123 (strong) | 2:1 | 1:1 | 1:2 | 1:8 |
| CD123 (weak) | 1:1 | 1:8 | 1:16 | 1:30 |
| CD371 | 1:4 | 1:16 | 1:40 | 1:60 |

**Supplementary Table 2:** Initial E:T ratio for each chimeric antigen receptor construct cytotoxic assay. Each experiment was seeded with 60000 cells and the four listed E:T ratios. For example, the cytotoxic assay using CD123-targeting (weak) CAR T-cells (first initial condition) was seeded with 40000 effector and 20000 target cells.

**Supplementary Table 3:** Structure of the experimental data. The first level consists of the five constructs, and the level elements are the names of the target receptors. The second and third levels are the type of effector and target cells, in order, in the co-culture cytotoxic assay. "uT" and "CART" refer to the untransduced and chimeric antigen receptor T-cells. "WT" and "KO" refer to the *TP53*-wildtype and -knockout tumor genotypes. The fourth level specifies the index of the initial conditions for the experiments (Supplementary Table 2). The fifth level indicates the index of the biological replication of the co-cultures. The sixth level consists of the time, reported in days, of the cell abundance measurements. The dataset, thus, consists of $5 \times 2 \times 2 \times 4 \times 2 \times 4 = 640$ measured data points.

| **Name** | **Values** |
|---|---|
| Antigen | ["CD33", "CD117", "CD123s", "CD123w", "CD371"] |
| Effector | ["uT", "CART"] |
| Target | ["WT", "KO"] |
| Ratio | [1, 2, 3, 4] |
| Sample | [1, 2] |
| Time Points | [1, 3, 6, 10] |

| | uncoupled T growth | uncoupled T death | Lotka-Volterra expansion | Lotka-Volterra exhaustion | Holling type 2 | Ratio-dependent expansion | Bazykin expansion | Beddington-DeAngelis |
|---|---|---|---|---|---|---|---|---|
| **CD33** | 21755.5 | -86.6034 | -242.624 | 4.35238e13 | 10000.9 | -305.584 | -340.857 | -340.27 |
| **CD117** | 14410.7 | 2.74009e18 | -179.122 | 9.88004e11 | -171.927 | -180.381 | -177.284 | -179.67 |
| **CD123s** | 2506.22 | 265.152 | -179.829 | 7.0961e10 | -176.06 | -190.404 | 529658.0 | -185.579 |
| **CD123w** | 7117.09 | -124.858 | -187.094 | 1.45765e14 | -186.412 | -191.215 | -188.986 | -189.182 |
| **CD371** | 11842.0 | -143.4 | -226.55 | 4.3724e13 | 5998.0 | -245.44 | -289.884 | -284.604 |

**Supplementary Table 4:** The negative log-likelihood summed over different effector and target combinations and initial effector-to-target ratios.

### Supplementary figures

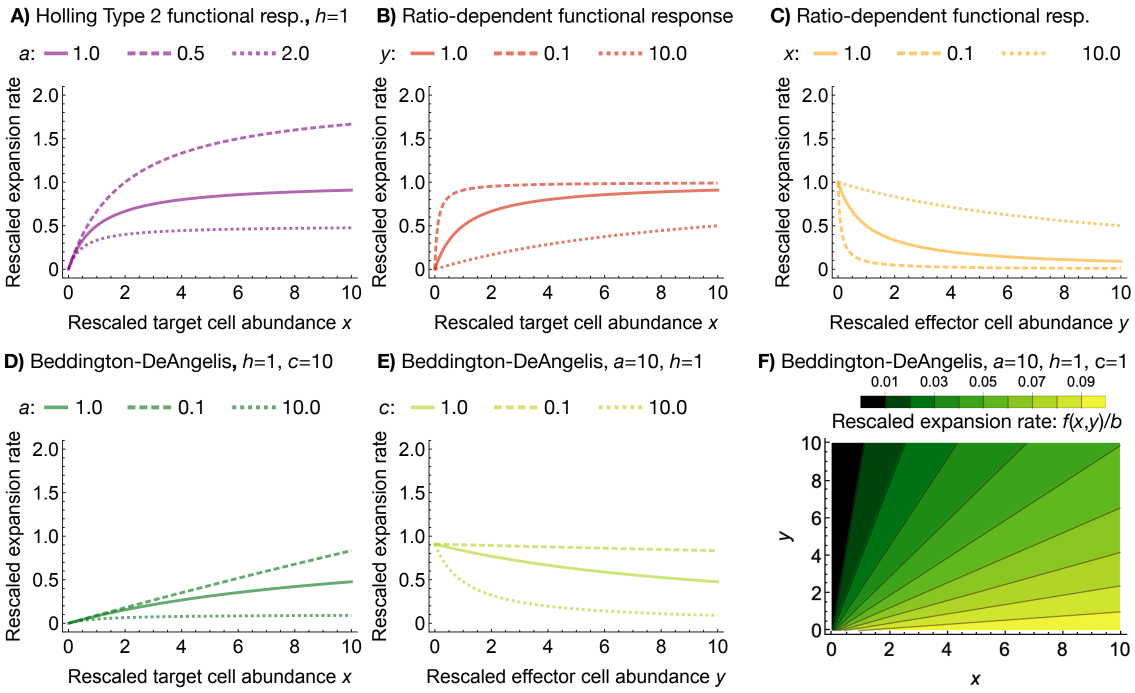


**Supplementary Figure 1: Behavior of key T cell expansion/activation functions. A**: Holling type 2 functional response example. **B**: and **C** Ratio-dependent functional response examples. **D**, **E**, and **F** Beddington-DeAngelis functional response examples. We rescaled the functional form in all plots using the baseline expansion rate parameter b. The other parameters are given in each panel.

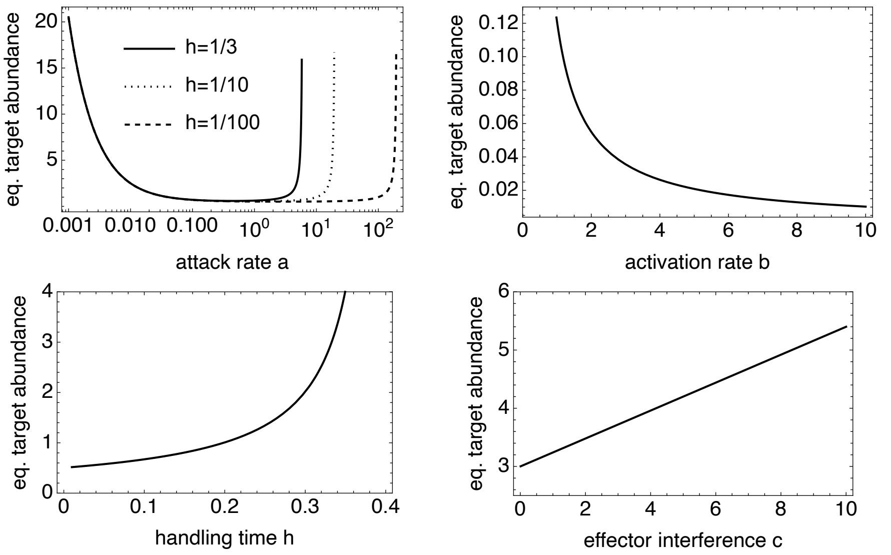


**Supplementary Figure 2: Long-term target abundance is affected by the performance of effector cells.** Each panel shows long-term target abundance as a function of one of the selected model's parameters (Beddington-DeAngelis). The target abundance decreases monotonically as the expansion rate $b$ increases. The increase in handling time $h$ and the effector interference $c$ increase the long-term target abundance. The range of attack rate $a$ that minimizes long-term target abundance depends on the handling time $h$. Attack rate $a$ lower or higher than this range increases long-term target abundance.

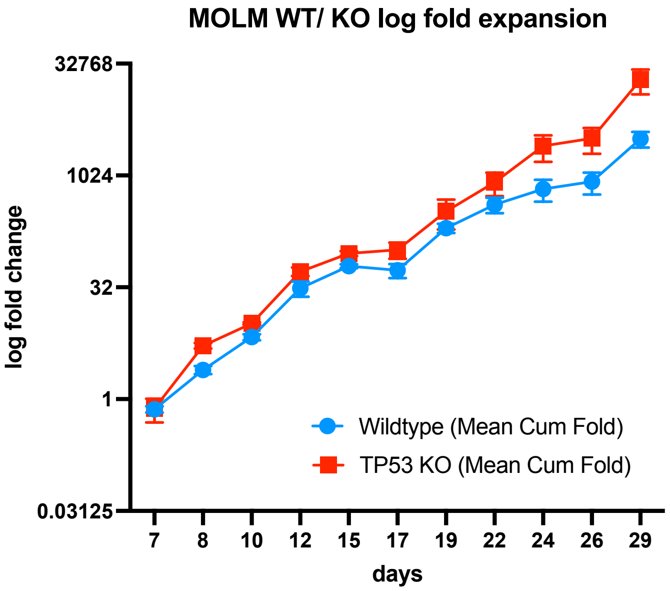


**Supplementary Figure 3: Experimental confirmation that wildtype (WT) and TP53 knockout (KO) MOLM13 cells do not differ in growth over the relevant time frame.** MOLM-13 TP53-wildtype and TP53-knockout target cells show comparable intrinsic proliferation over the timescale of the cytotoxic co-culture assays. Split-corrected cumulative fold expansion of MOLM-13-TP53+/+ (blue) and MOLM-13-TP53-/- (red) cells cultured in the absence of T or CAR T effector cells, normalized to day 7 (baseline $0.8\times10^6$ cells, $0.4\times10^6$/mL $\times$ 2 mL per well). Cells were maintained in parallel wells of a 6-well suspension plate ($n = 3$ wells per genotype) and re-seeded at $0.4 \times 10^6$/mL at each passage. Symbols show mean±SEM across wells. The two genotypes display overlapping expansion trajectories from day 7 through day 19, spanning the duration of the LT15 (day 10) and LT6 (day 16) cytotoxic co-culture assays used for model training and validation. A modest divergence emerges thereafter, with TP53-KO cells showing a higher long-term cumulative expansion than TP53-WT cells by day 29. Because the cytotoxic co-culture assays terminate well within the equivalence window, the target-antigen-dependent differences in CAR T kinetic parameters reported in the main text cannot be attributed to intrinsic target-cell growth differences over the relevant timescale.

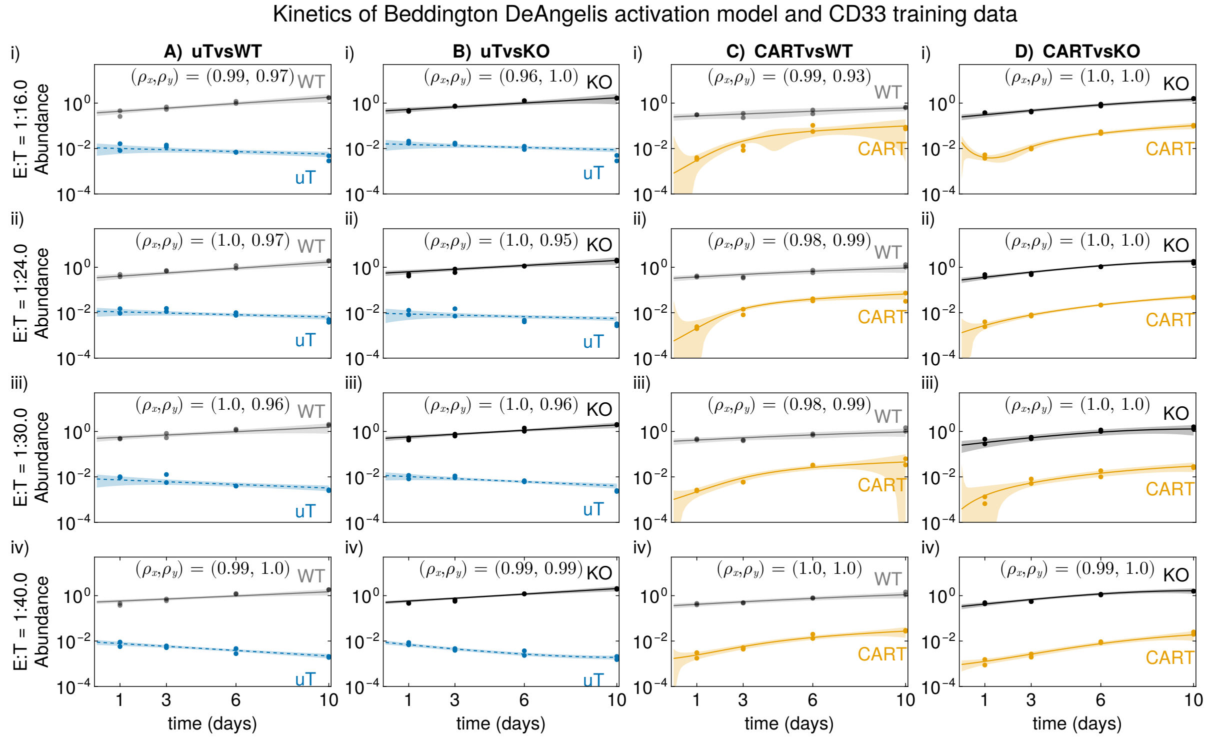


**Supplementary Figure 4: *In-vitro* cytolytic assay for CD33 targeting chimieric-antigen-receptor cells and numerical solution of the Beddington-DeAngelis expansion model.** Each column (**A-D**) uniquely combines an effector and a target cell type. Each row has a different initial effector-to-target ratio. The dots represent experimental measurements of cell abundance via fluorescence-activated cell sorting, and the solid lines are numerical solutions of the Beddington-DeAngelis expansion model. The black represents the target, and the other colors represent the effector cell abundances.

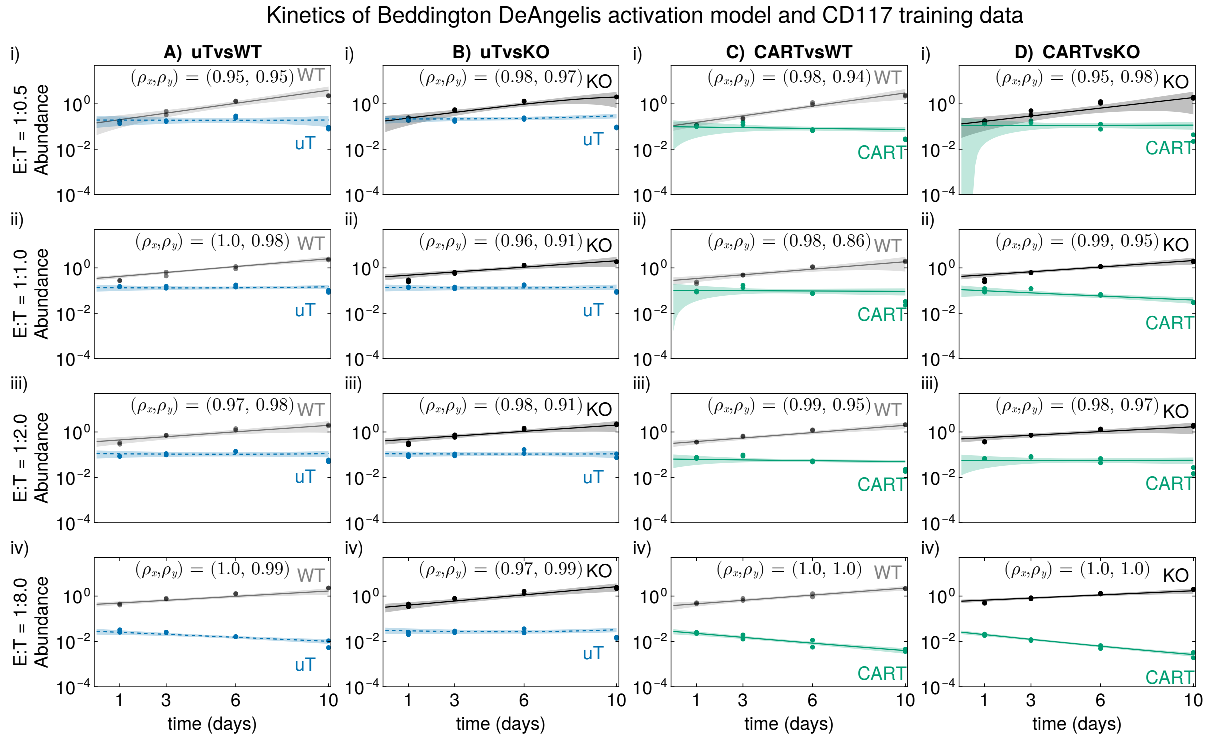


**Supplementary Figure 5: *In-vitro* cytolytic assay for CD117 targeting chimieric-antigen-receptor cells and numerical solution of the Beddington-DeAngelis expansion model.** Please see Supplementary Fig. 4 for panel description.

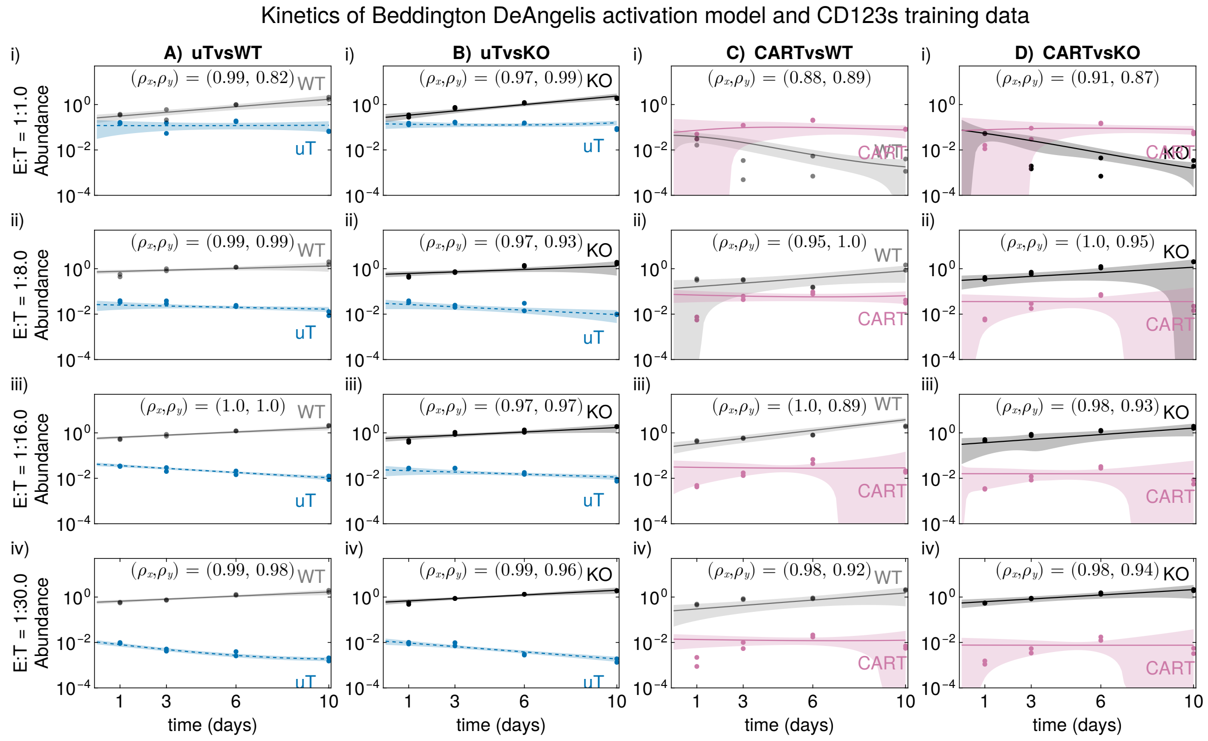


**Supplementary Figure 6: *In-vitro* cytolytic assay for CD123s targeting chimieric-antigen-receptor cells and numerical solution of the Beddington-DeAngelis expansion model.** Please see Supplementary Fig. 4 for panel description.

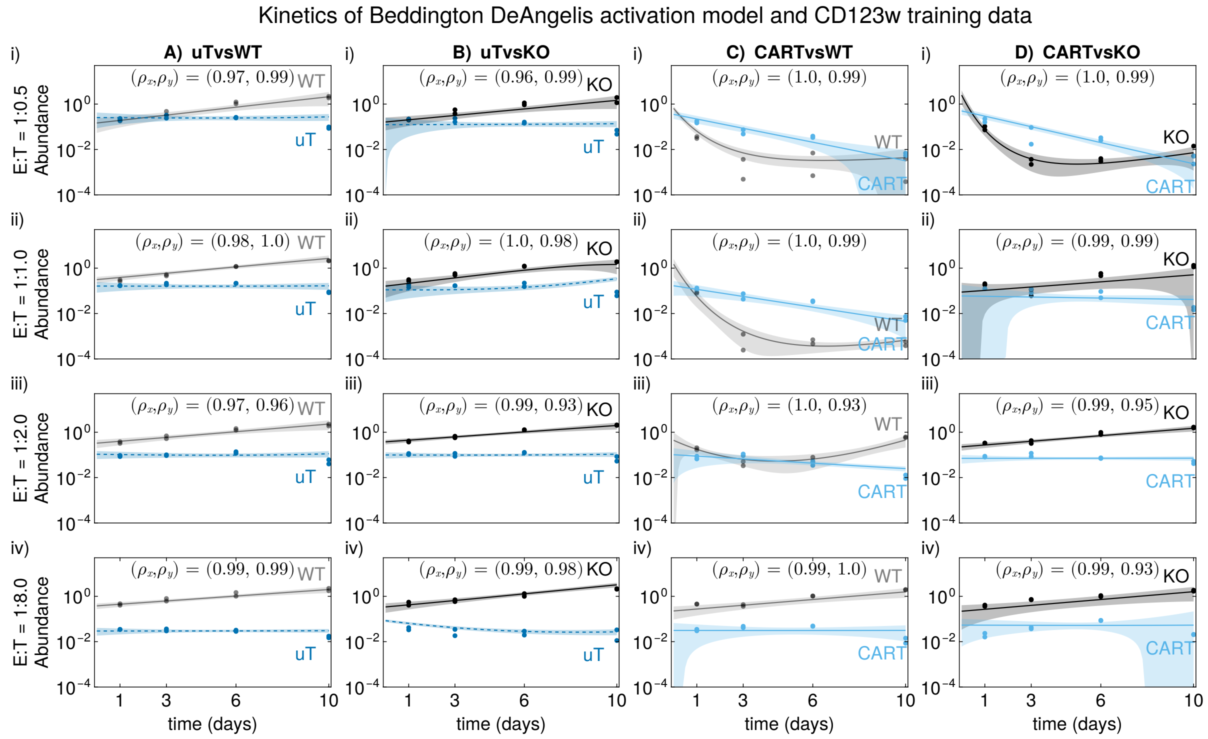


**Supplementary Figure 7: *In-vitro* cytolytic assay for CD123w targeting chimieric-antigen-receptor cells and numerical solution of the Beddington-DeAngelis expansion model.** Please see Supplementary Fig. 4 for panel description.

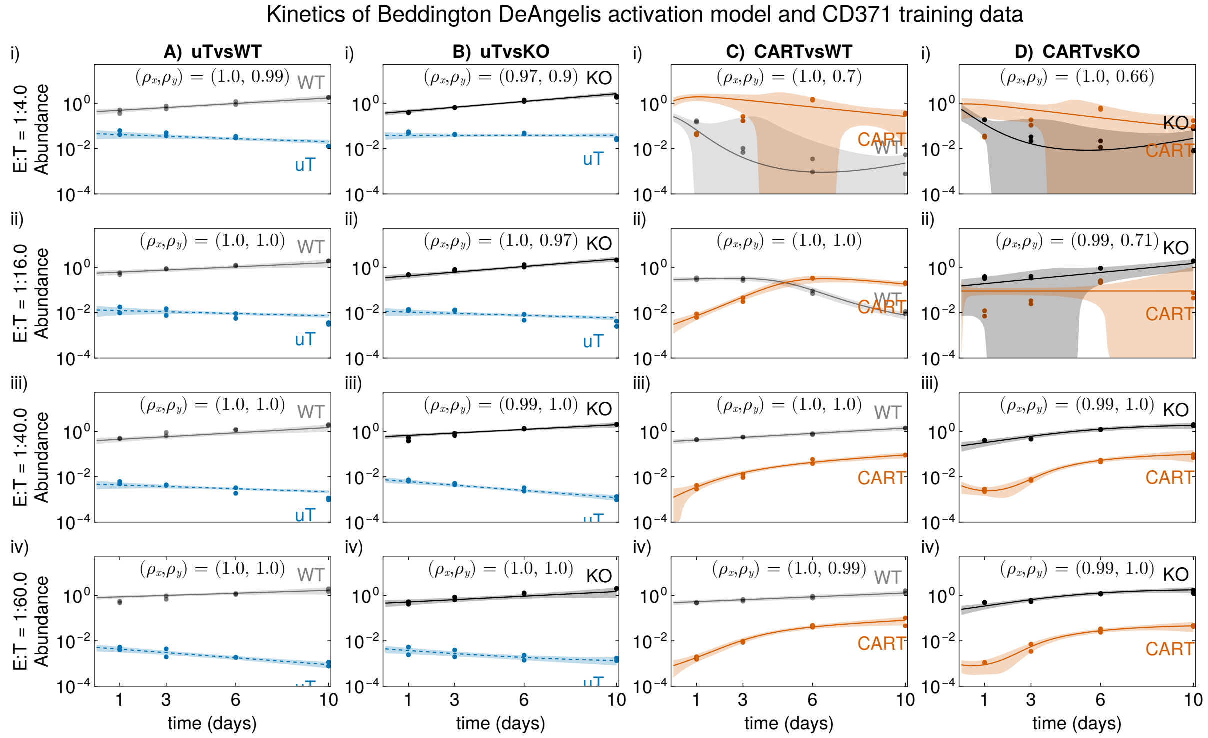


**Supplementary Figure 8: *In-vitro* cytolytic assay for CD371 targeting chimieric-antigen-receptor cells and numerical solution of the Beddington-DeAngelis expansion model.** Please see Supplementary Fig. 4 for panel description.

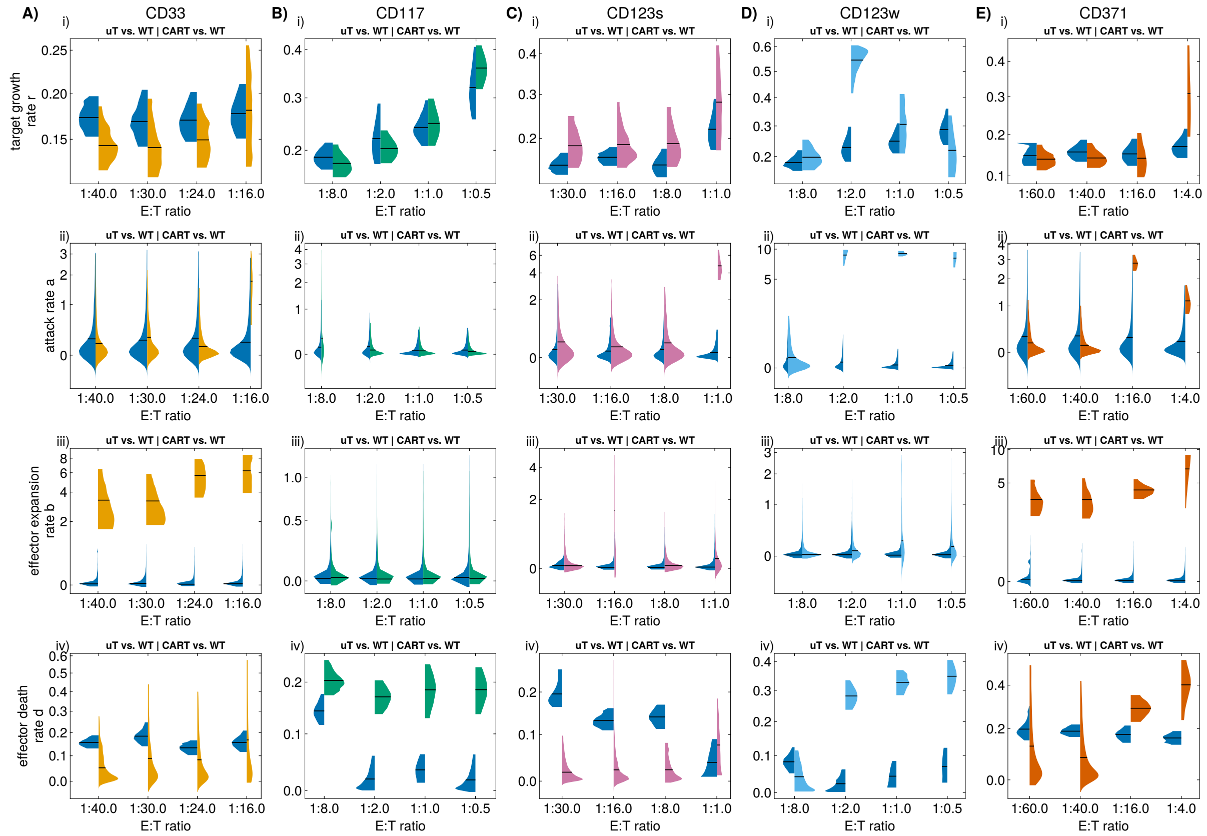


**Supplementary Figure 9: Quantification of heterogeneous effector performance**. Each column A)-E) represents a specific target construct. Each line, sub-panels i)-iv), shows posterior parameter distributions of four important parameters that characterize T cell performance. In each sub-panel, we compare the performance of uT cells (left halves of the violin) with CART cells (right halves) against WT target leukemia cells across experimentally realized initial conditions (E:T ratios). Sometimes, the parameter distributions change with initial E:T ratios. Overall, we see a markedly different CAR T cell performance against WT leukemia cells, mainly in the direction of better killing and persistence. WT: TP53-wildtype target, KO: TP53-knockout target, uT: untransduced T, CART: chimeric antigen receptor T cells.

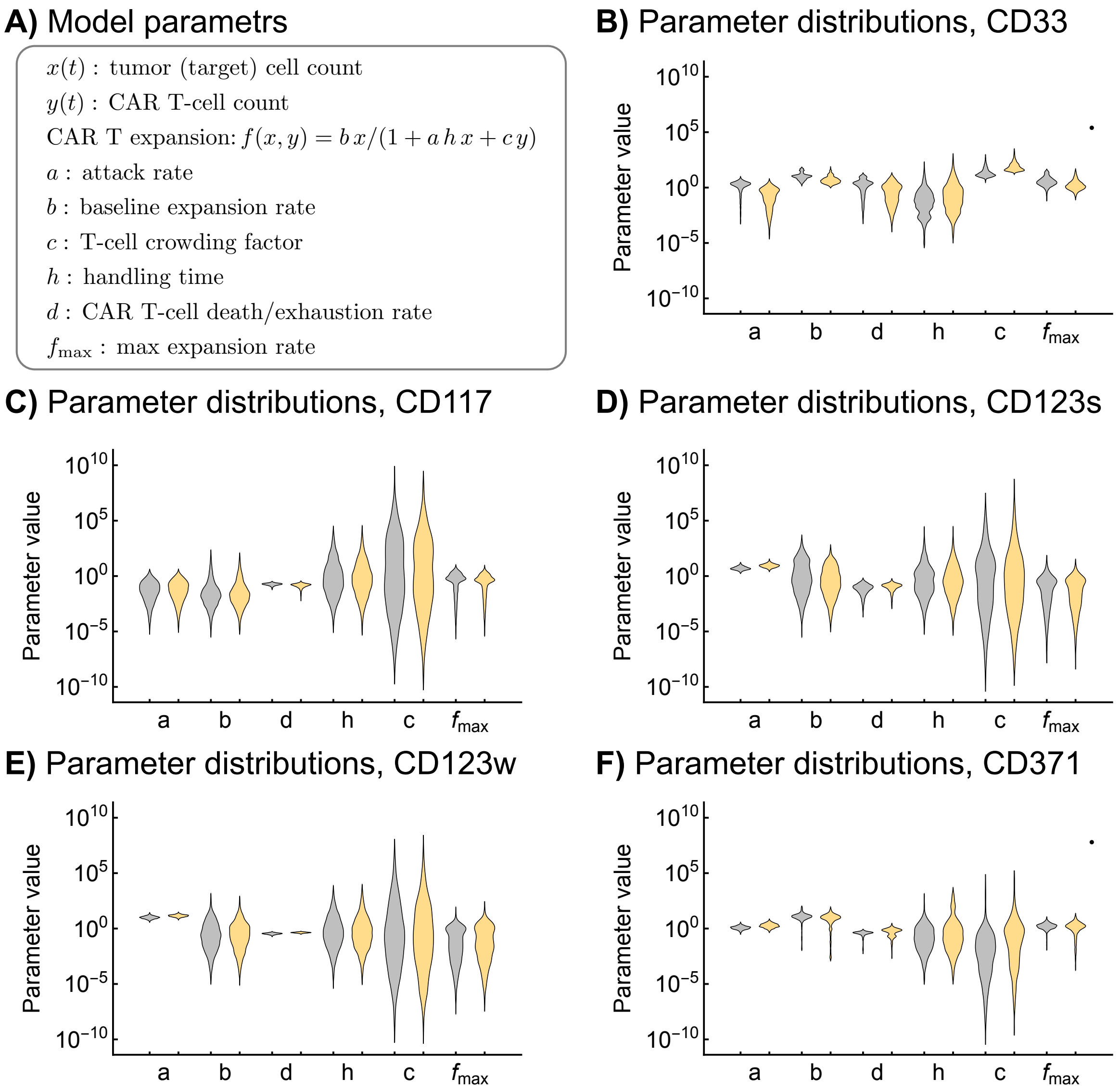


**Supplementary Figure 10: Violin plots of parameter distributions for all five CAR constructs.** A) Recap of the model parameters and the derived quantity of maximal expansion (over time, see main text). B)-F) Violin plots comparing the posterior parameter distributions against WT AML (gray) and against KO AML (yellow).

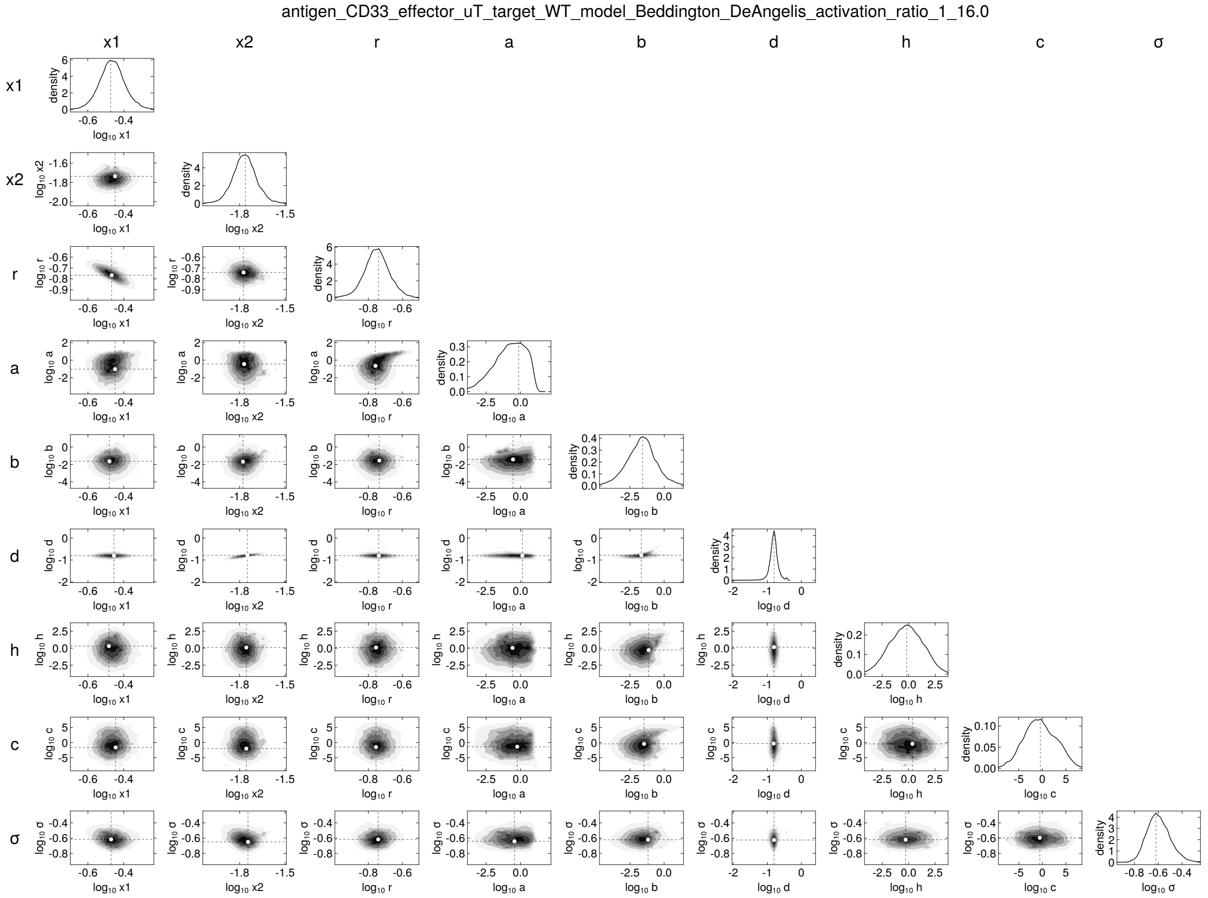


**Supplementary Figure 11: Posterior distributions over parameter values of Beddington-DeAngelis expansion model.** The title describes the antigen target of the chimeric-antigen-receptor, the type of effector and tumor cell, the mathematical model, and the initial effector-to-target ratio. Each diagonal panel shows marginal posterior distribution over one parameter. Each of the remaining panels is a contour plot of the unique pair of parameters of the Beddington-DeAngelis expansion model. The white dot depicts the mode of each distribution computed by finding the point with the highest posterior density. Closely placed contours, i.e., the degree of curvature, are related to the inferential precision. The oval closed contours in the joint posterior density suggest parameters are practically identifiable. Contour lines separated over one order of magnitude or lacking contour lines, i.e., a flat profile, indicate non-identifiability.s

antigen_CD33_effector_uT_target_WT_model_Beddington_DeAngelis_activation_ratio_1_24.0

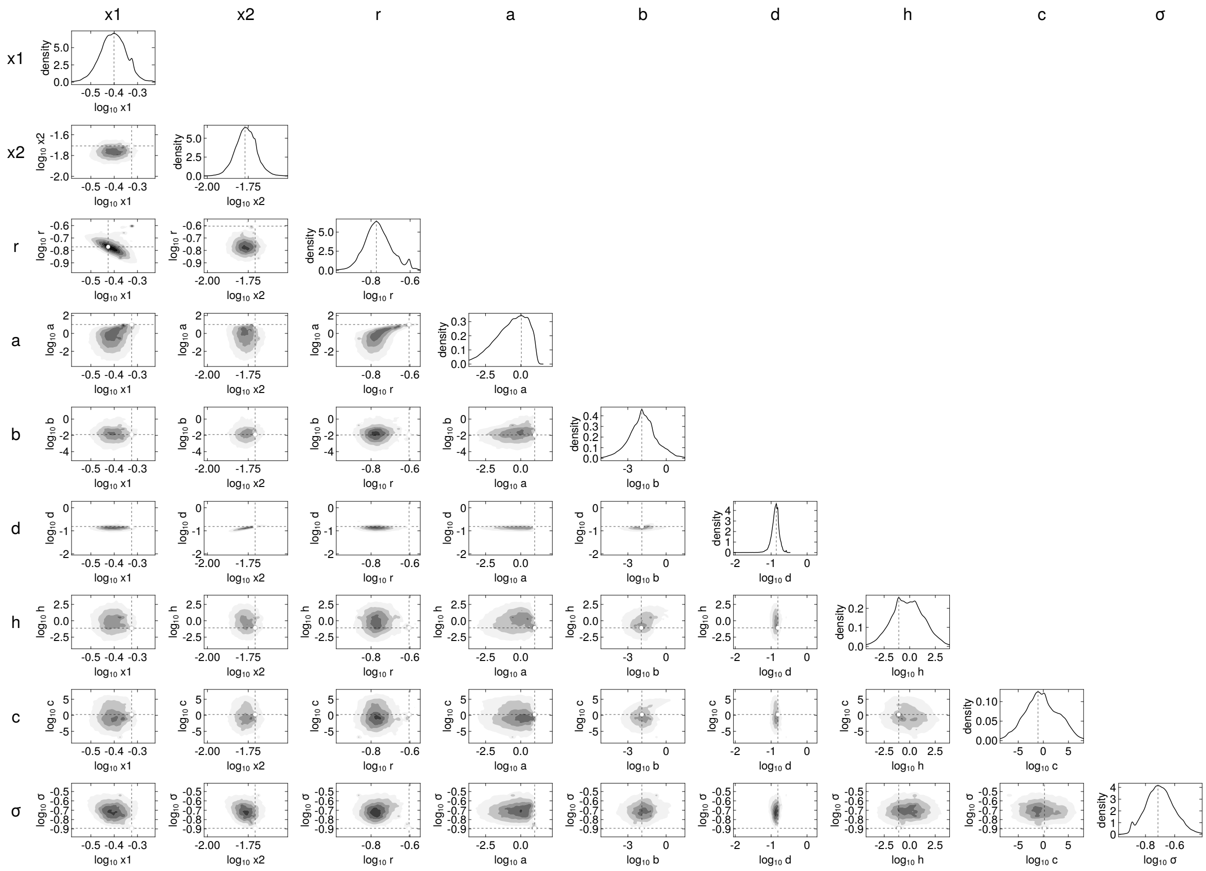


206

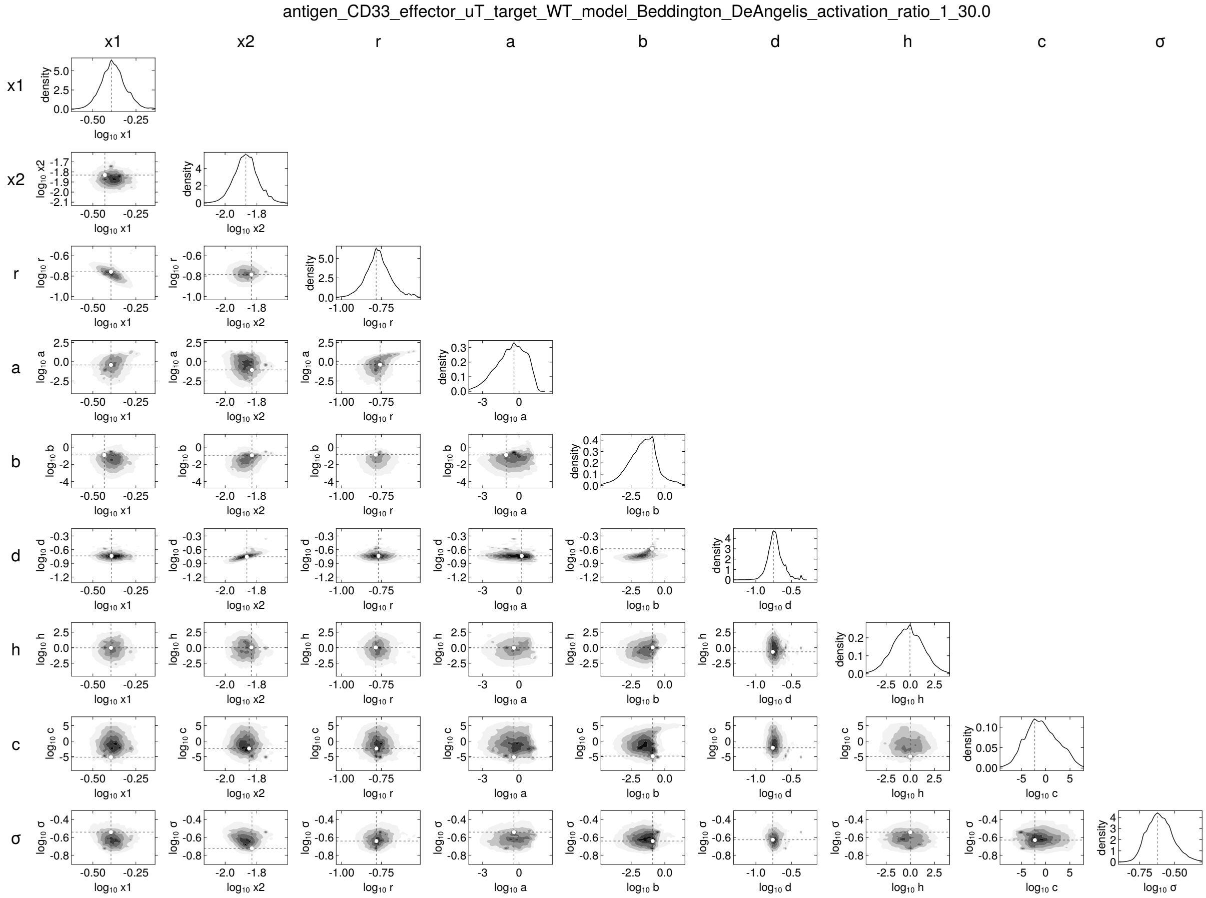
antigen_CD33_effector_uT_target_WT_model_Beddington_DeAngelis_activation_ratio_1_30.0
x1
x2
r
a
b
d
h
c
σ
density
log10 x1
log10 x2
log10 r
log10 a
log10 b
log10 d
log10 h
log10 c
log10 σ

antigen_CD33_effector_uT_target_WT_model_Beddington_DeAngelis_activation_ratio_1_40.0

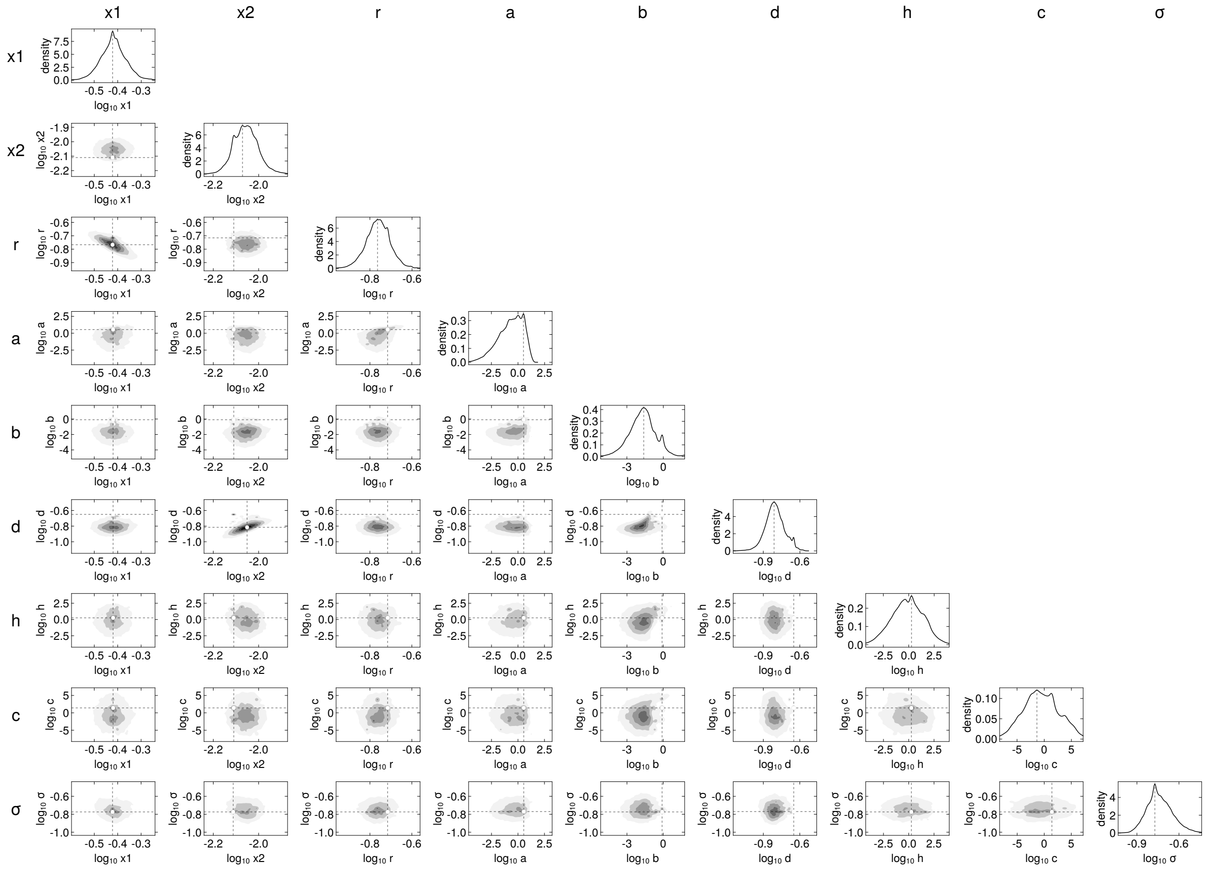


208

antigen_CD33_effector_uT_target_KO_model_Beddington_DeAngelis_activation_ratio_1_16.0

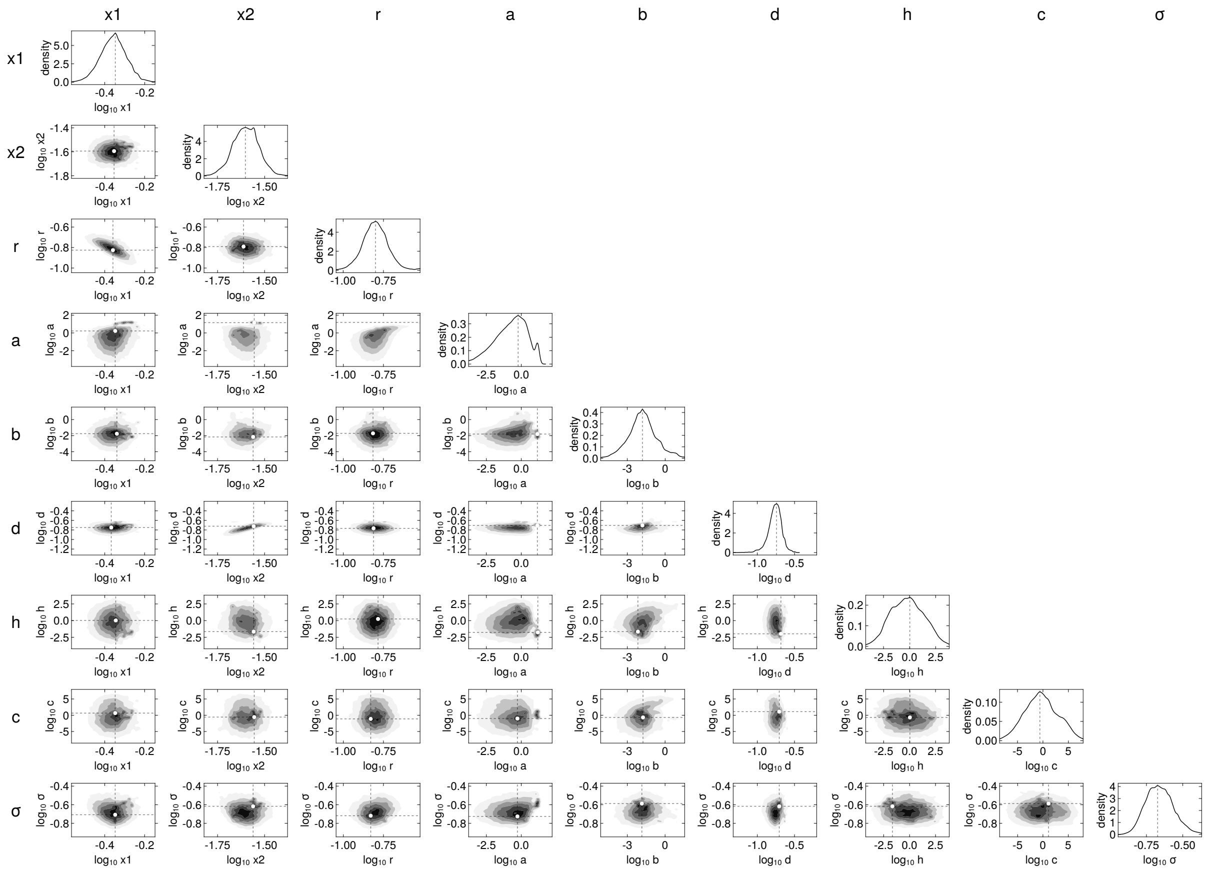

antigen_CD33_effector_uT_target_KO_model_Beddington_DeAngelis_activation_ratio_1_24.0

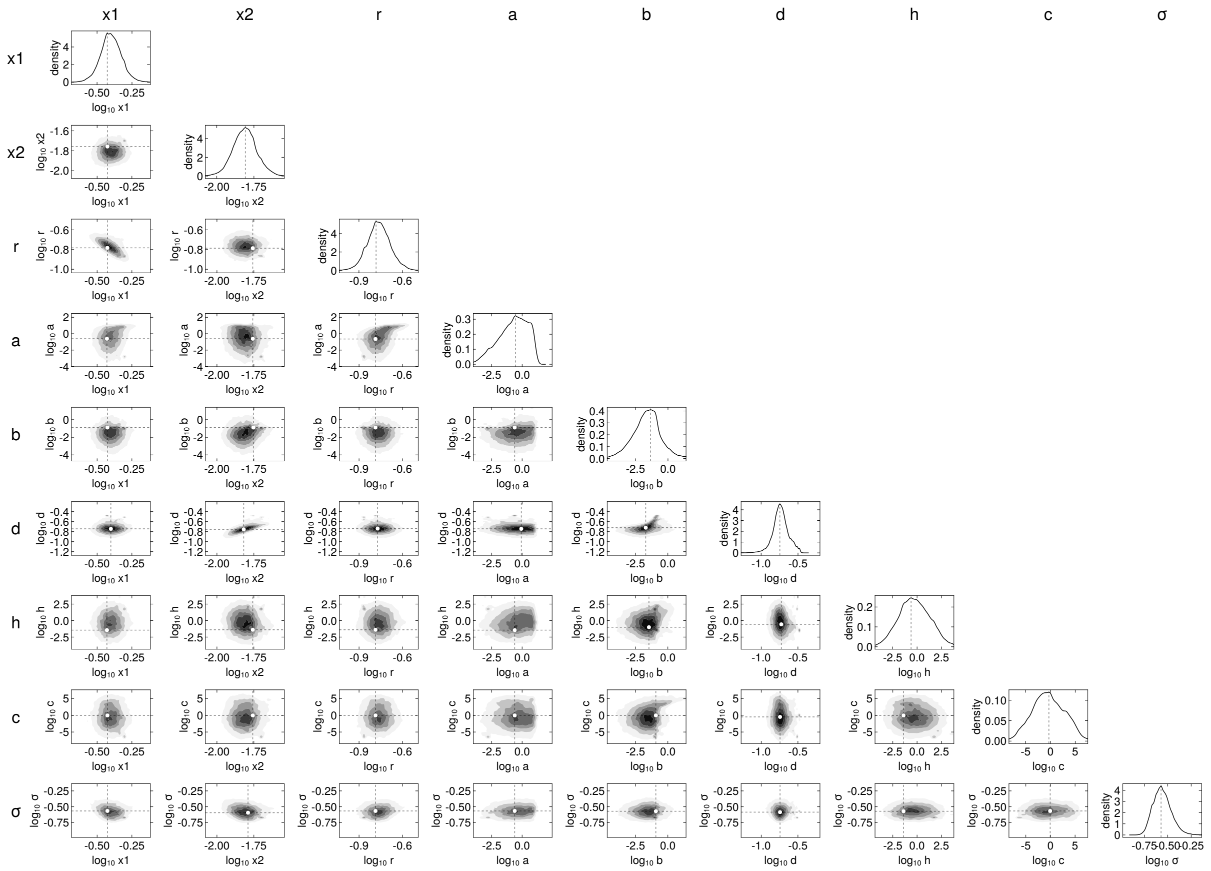

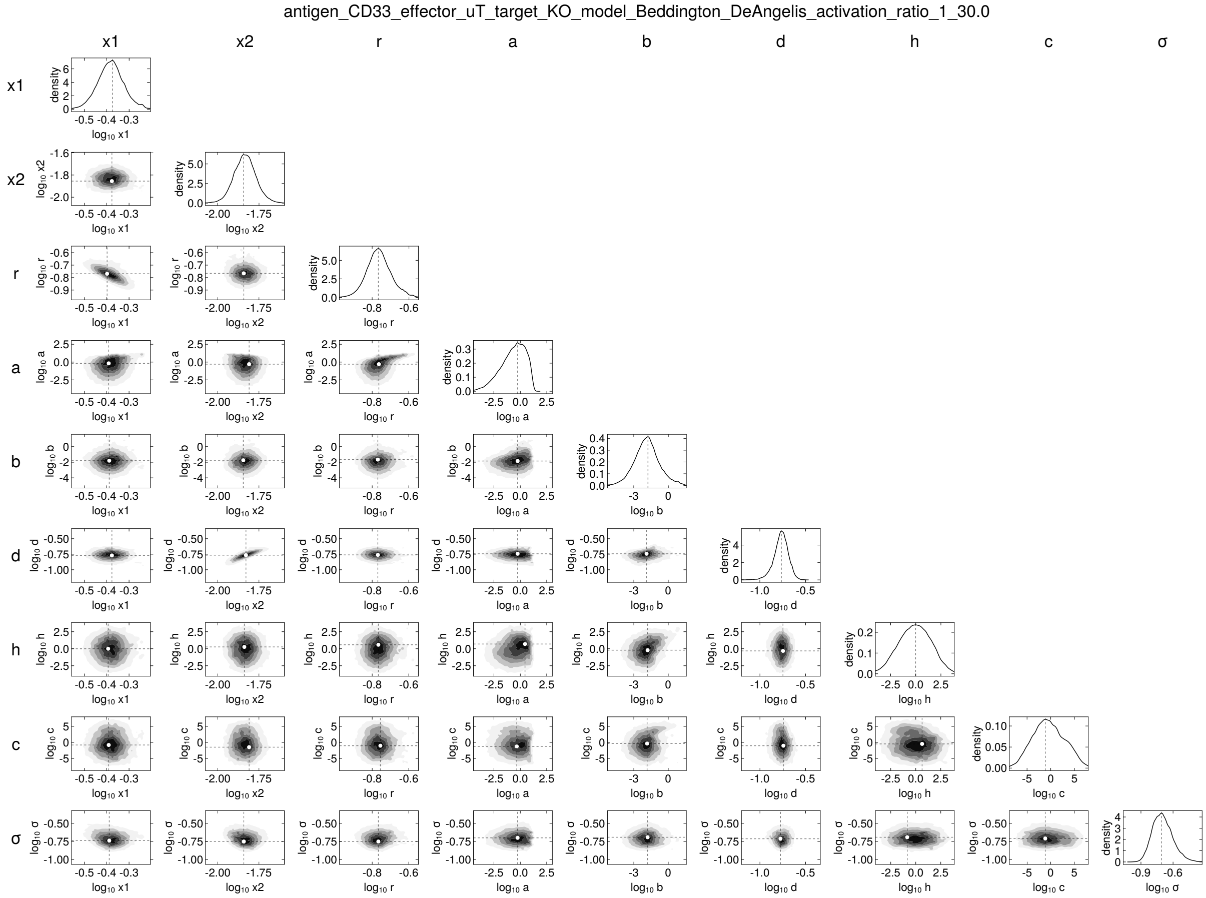
antigen_CD33_effector_uT_target_KO_model_Beddington_DeAngelis_activation_ratio_1_30.0
x1
x2
r
a
b
d
h
c
σ
density
log10 x1
log10 x2
log10 r
log10 a
log10 b
log10 d
log10 h
log10 c
log10 σ

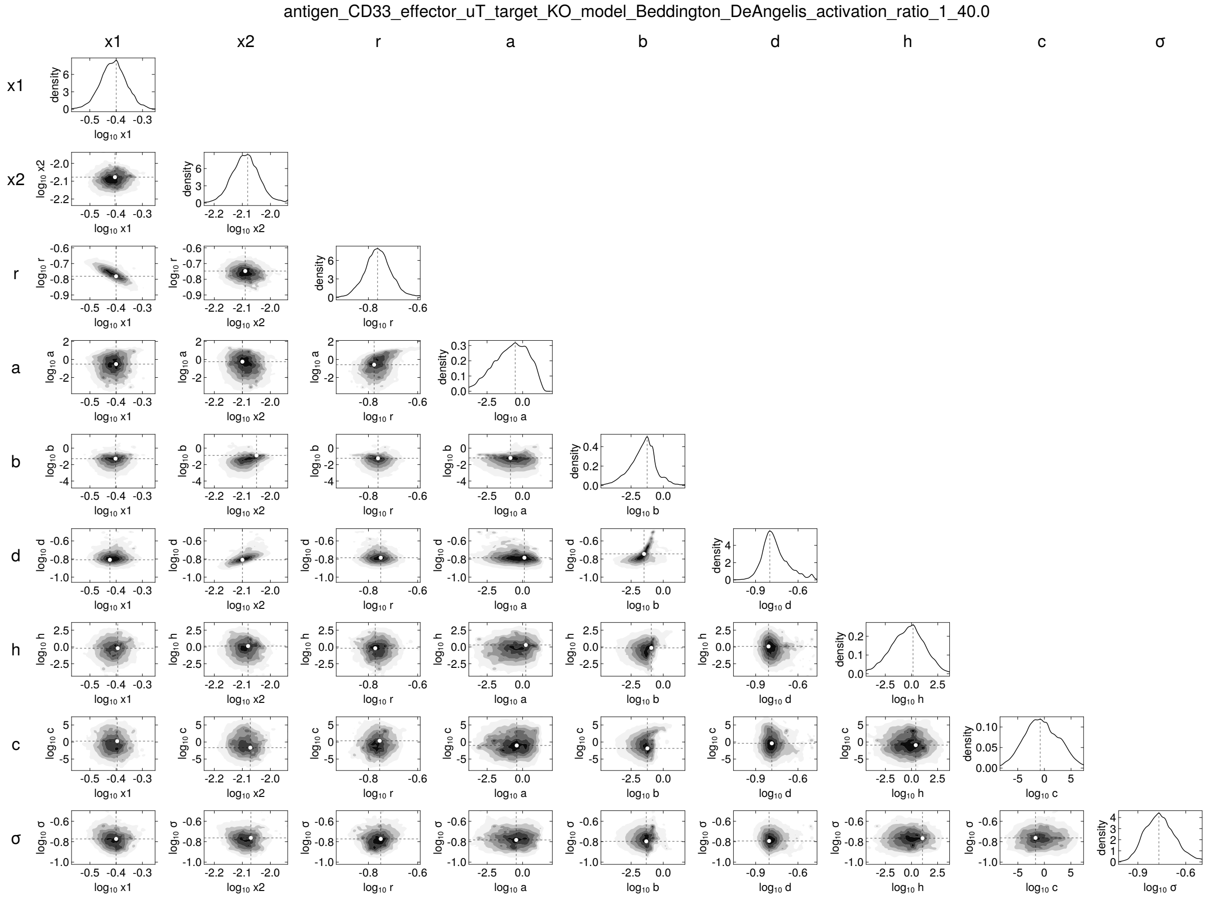
antigen_CD33_effector_uT_target_KO_model_Beddington_DeAngelis_activation_ratio_1_40.0
x1
x2
r
a
b
d
h
c
σ

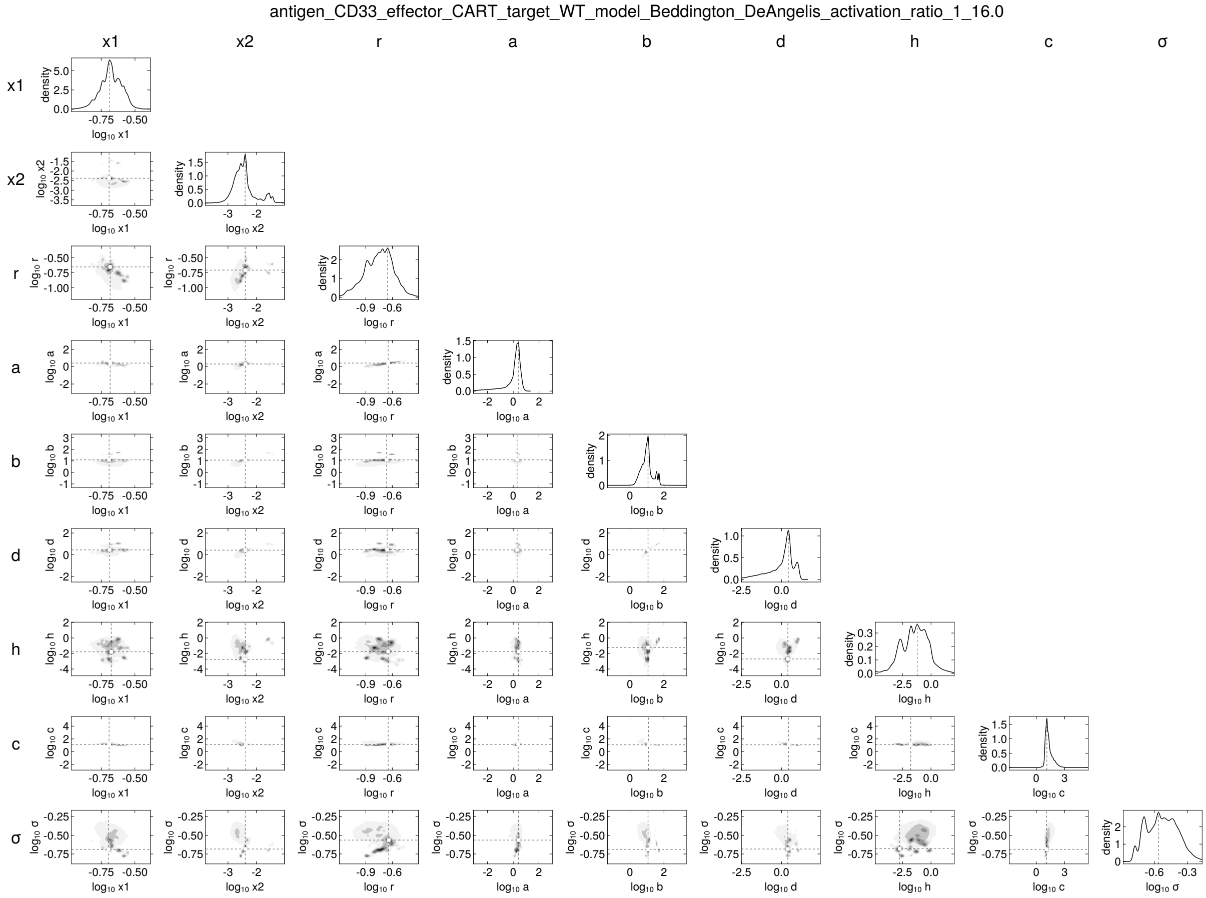
antigen_CD33_effector_CART_target_WT_model_Beddington_DeAngelis_activation_ratio_1_16.0
x1
x2
r
a
b
d
h
c
σ
density
log10 x1
log10 x2
log10 r
log10 a
log10 b
log10 d
log10 h
log10 c
log10 σ

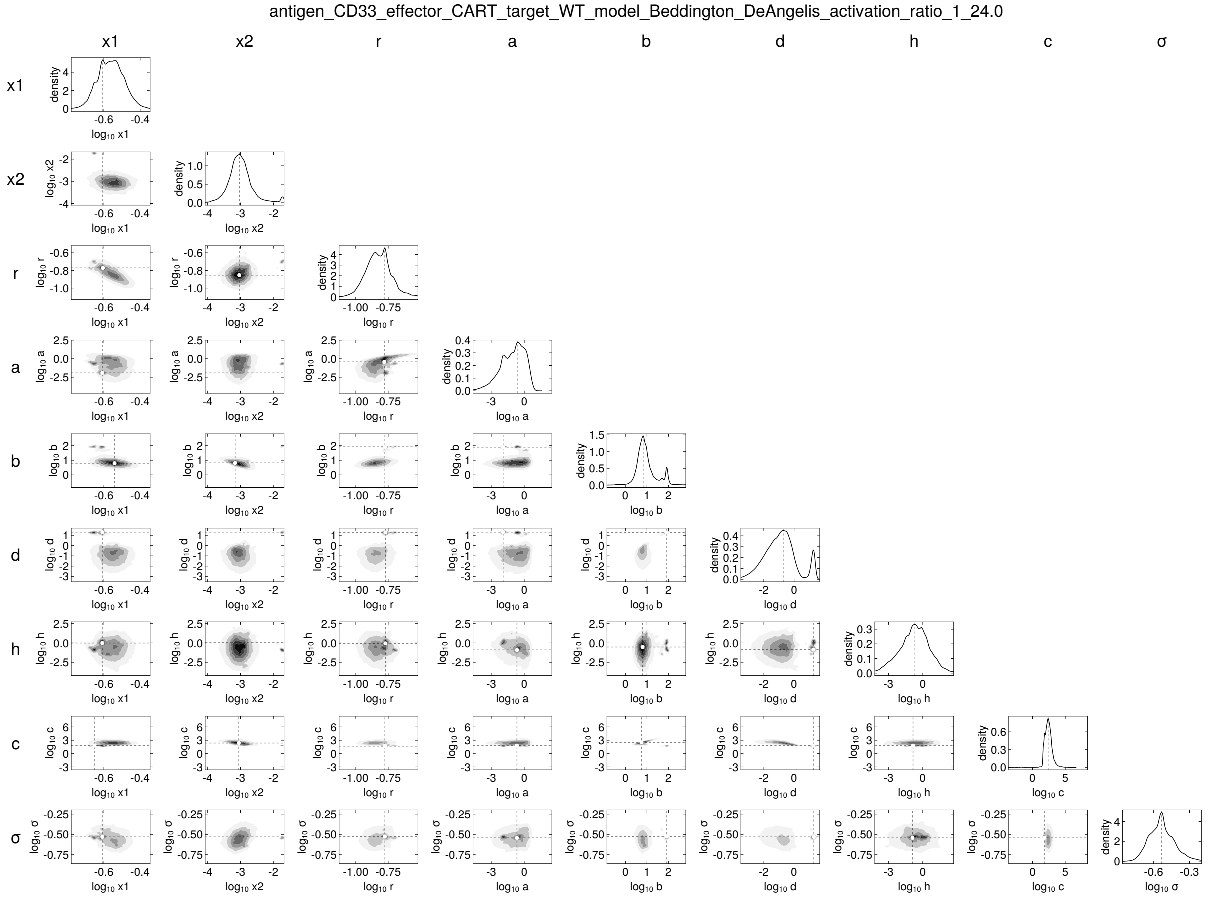
antigen_CD33_effector_CART_target_WT_model_Beddington_DeAngelis_activation_ratio_1_24.0
x1
x2
r
a
b
d
h
c
σ

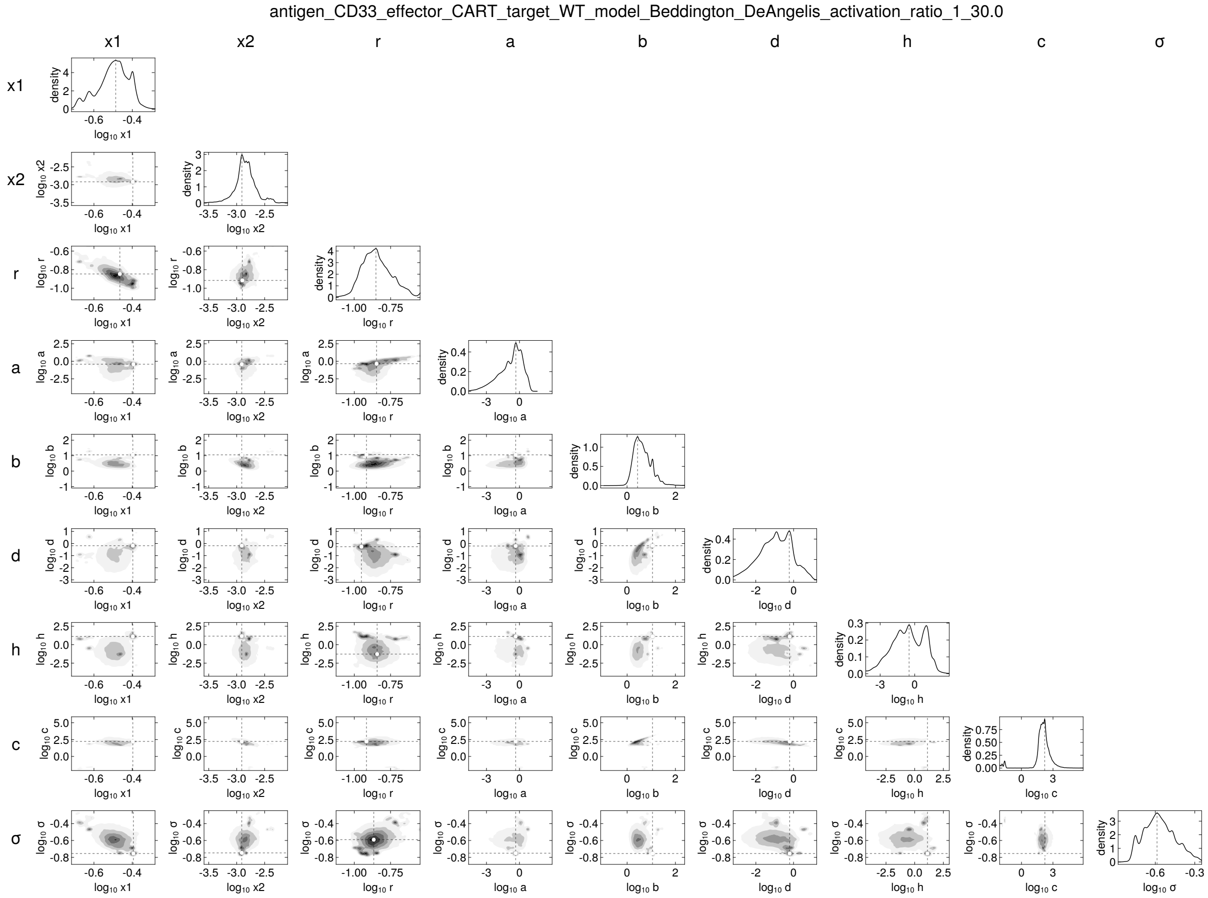
antigen_CD33_effector_CART_target_WT_model_Beddington_DeAngelis_activation_ratio_1_30.0
x1
x2
r
a
b
d
h
c
σ
density

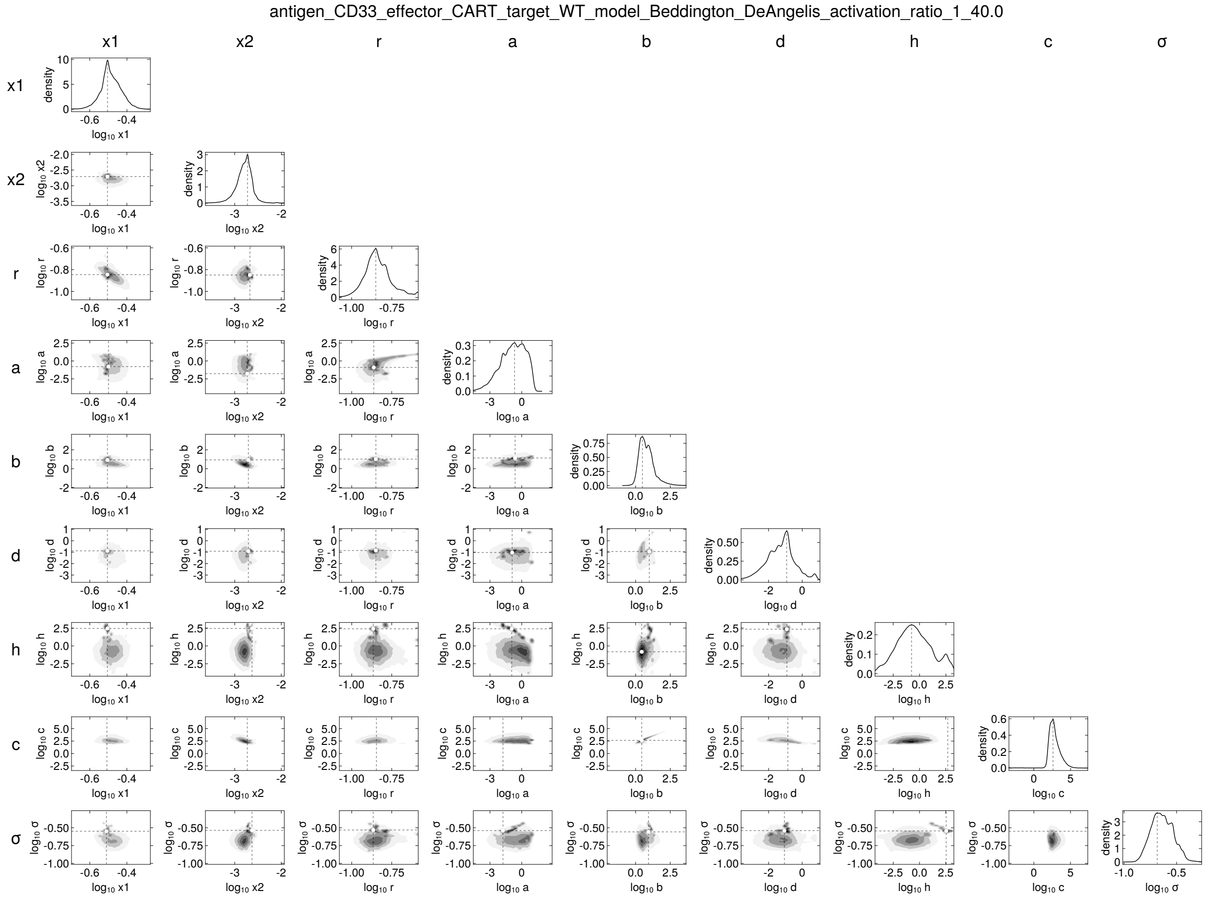
antigen_CD33_effector_CART_target_WT_model_Beddington_DeAngelis_activation_ratio_1_40.0
x1
x2
r
a
b
d
h
c
σ
density
log10 x1
log10 x2
log10 r
log10 a
log10 b
log10 d
log10 h
log10 c
log10 σ

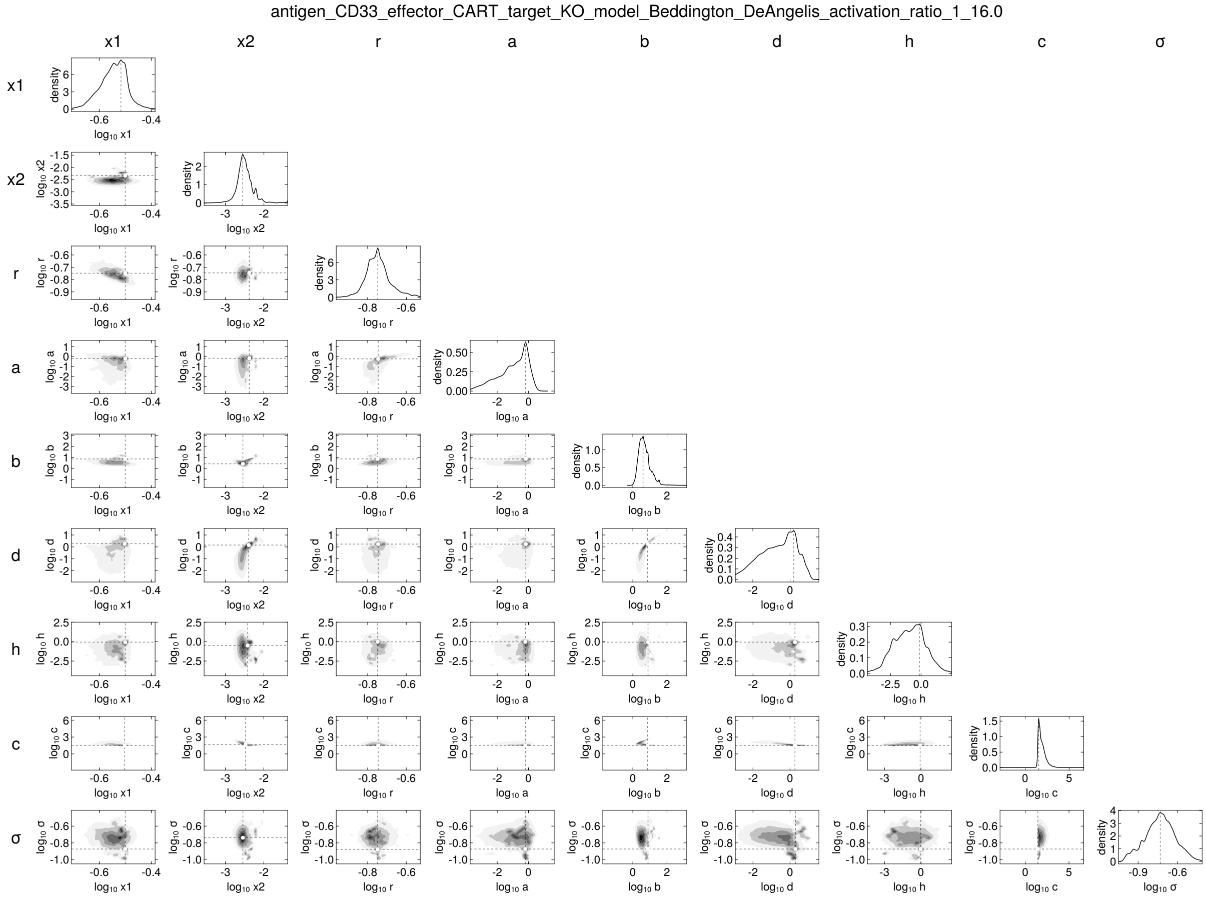
antigen_CD33_effector_CART_target_KO_model_Beddington_DeAngelis_activation_ratio_1_16.0
x1
x2
r
a
b
d
h
c
σ

217

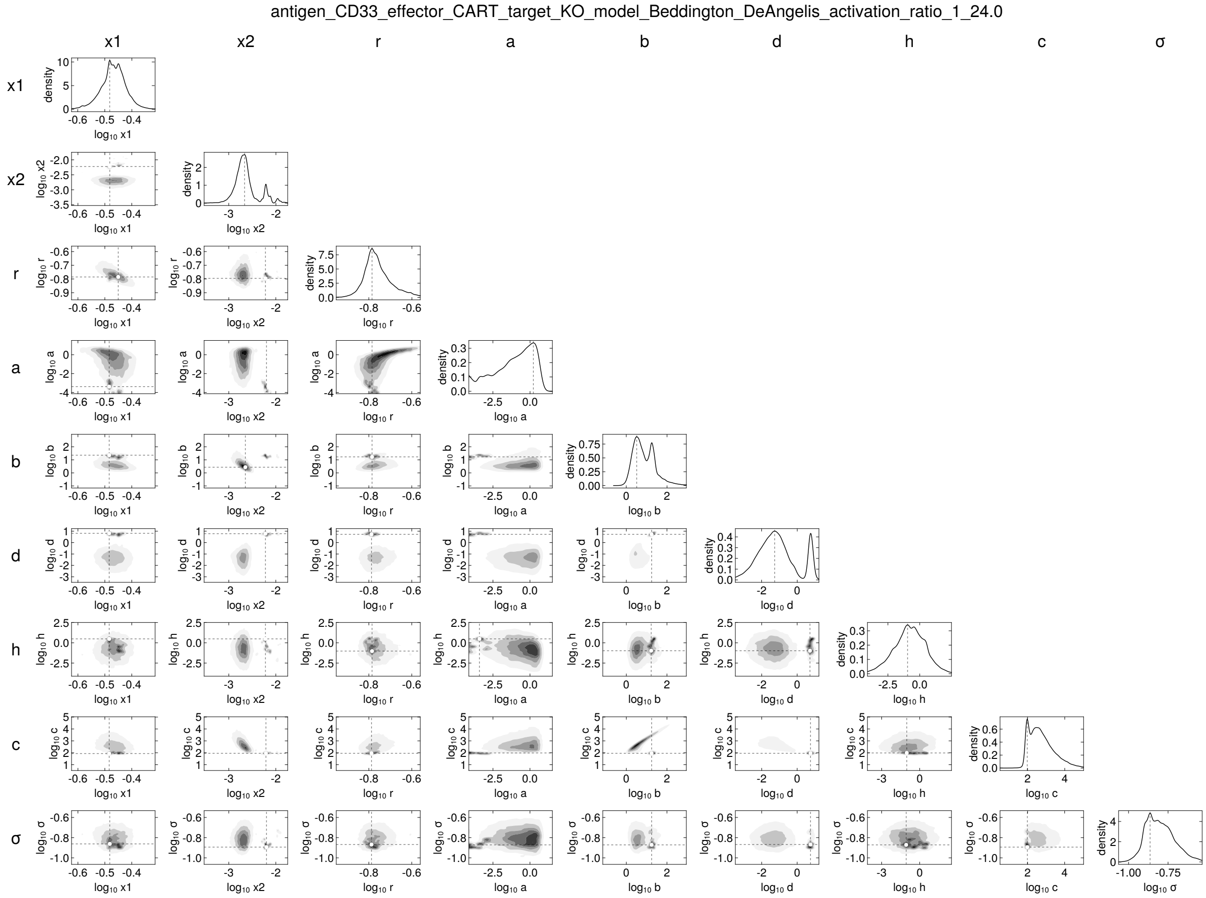
antigen_CD33_effector_CART_target_KO_model_Beddington_DeAngelis_activation_ratio_1_24.0
x1
x2
r
a
b
d
h
c
σ
density
$\log_{10}$ x1
$\log_{10}$ x2
$\log_{10}$ r
$\log_{10}$ a
$\log_{10}$ b
$\log_{10}$ d
$\log_{10}$ h
$\log_{10}$ c
$\log_{10}$ σ

218

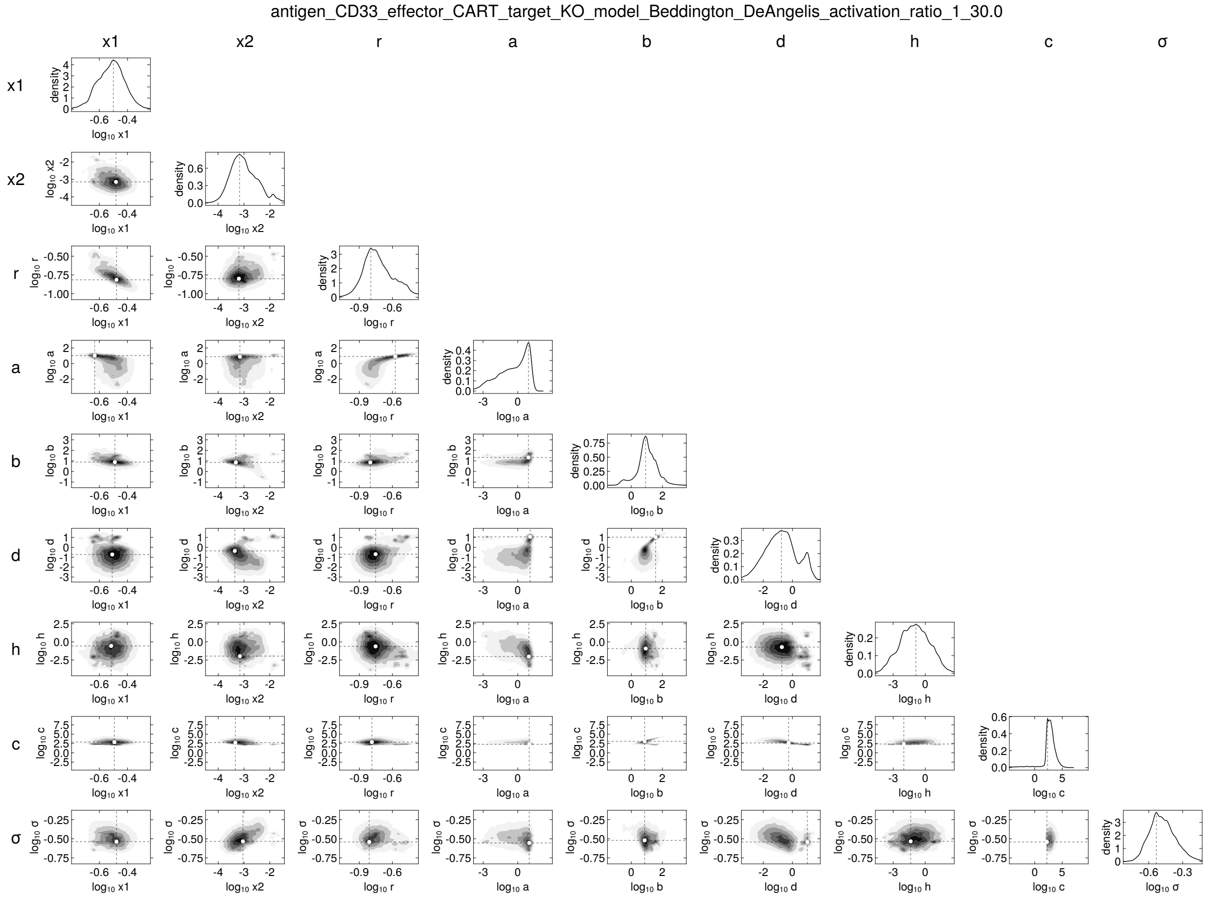
antigen_CD33_effector_CART_target_KO_model_Beddington_DeAngelis_activation_ratio_1_30.0
x1
x2
r
a
b
d
h
c
σ

antigen_CD33_effector_CART_target_KO_model_Beddington_DeAngelis_activation_ratio_1_40.0

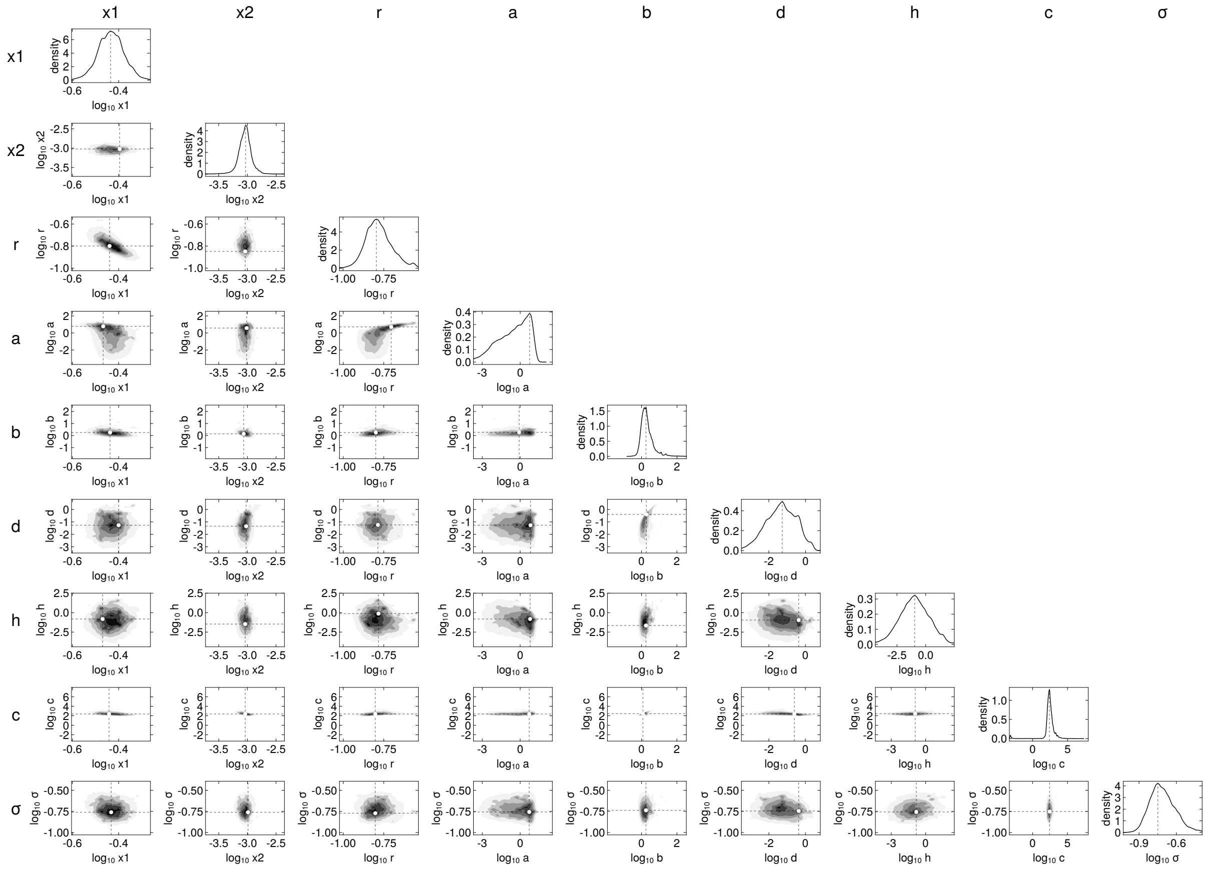


220

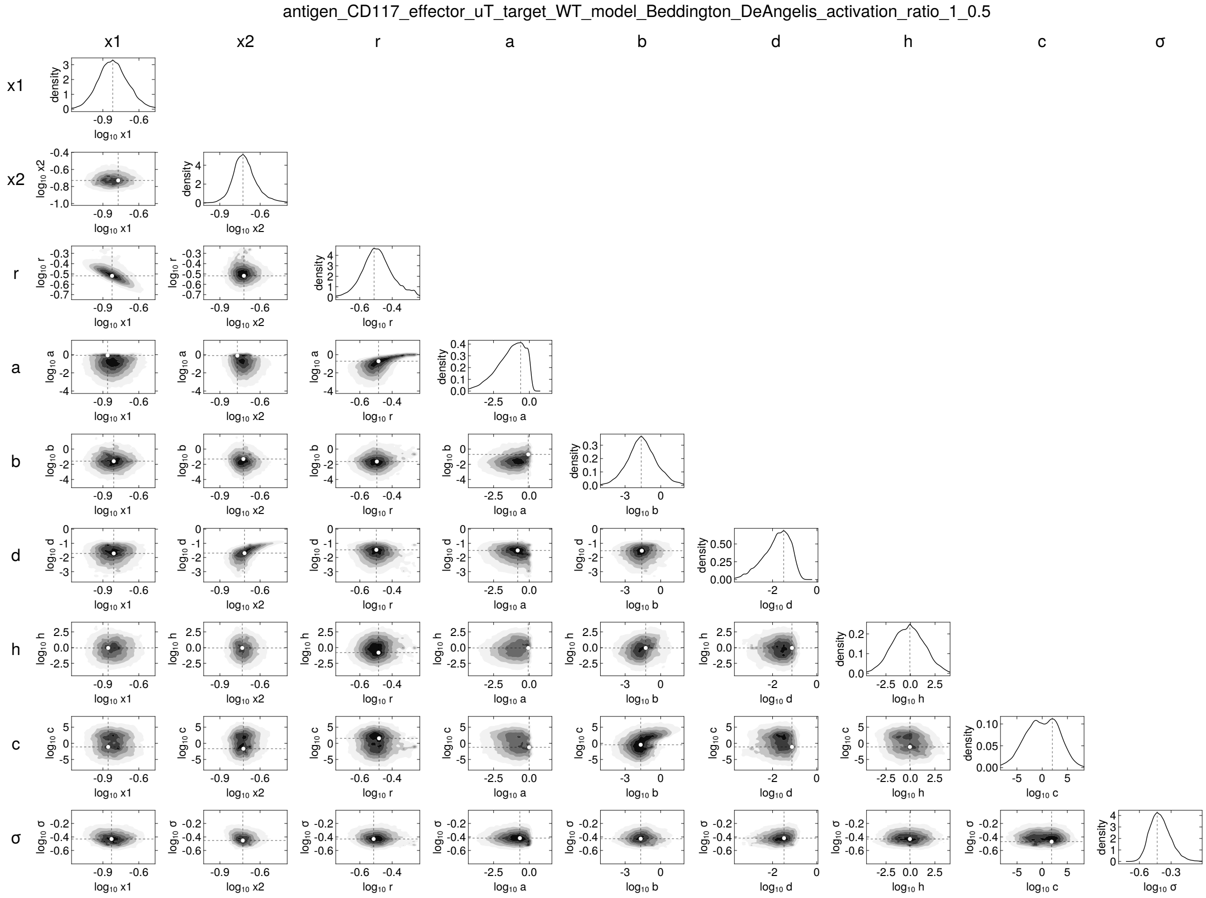
antigen_CD117_effector_uT_target_WT_model_Beddington_DeAngelis_activation_ratio_1_0.5
x1
x2
r
a
b
d
h
c
σ
density
log10 x1
log10 x2
log10 r
log10 a
log10 b
log10 d
log10 h
log10 c
log10 σ

antigen_CD117_effector_uT_target_WT_model_Beddington_DeAngelis_activation_ratio_1_1.0

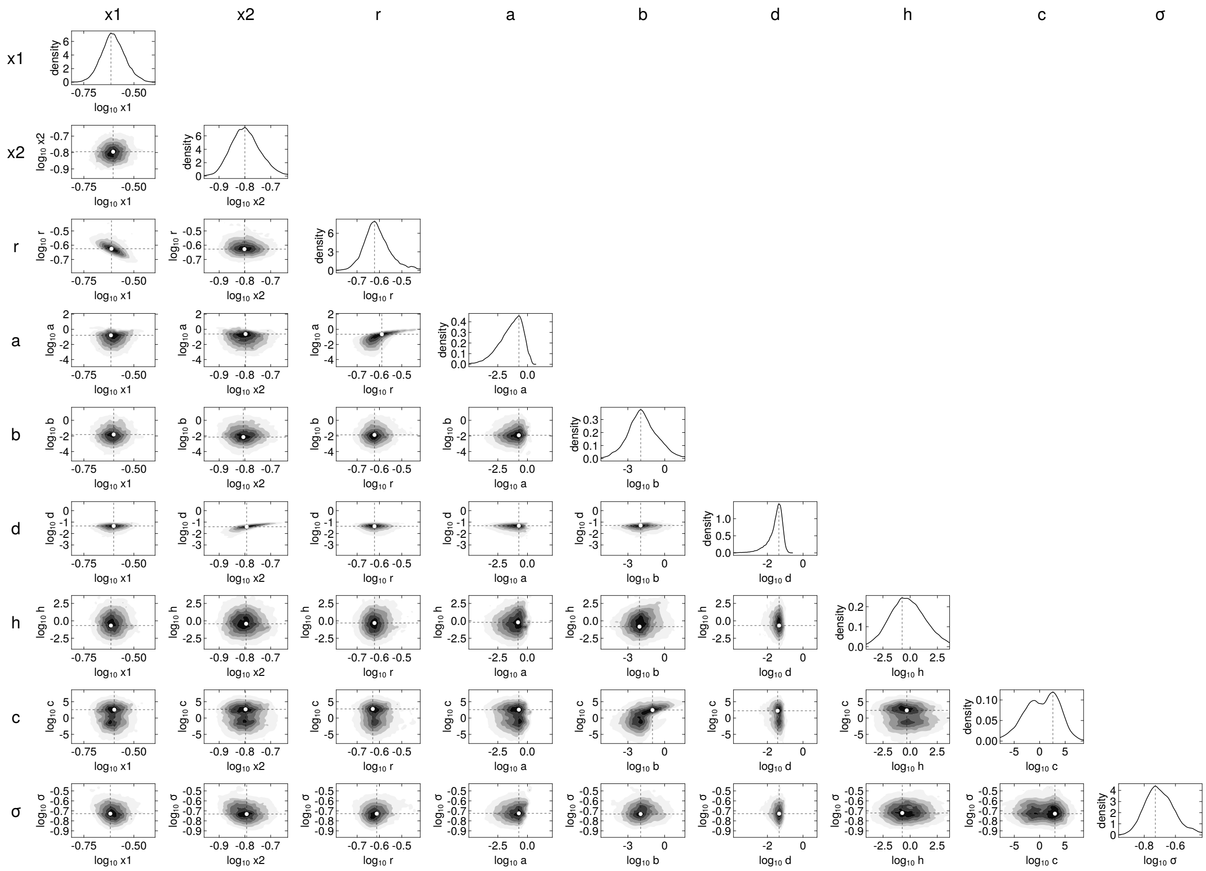

antigen_CD117_effector_uT_target_WT_model_Beddington_DeAngelis_activation_ratio_1_2.0

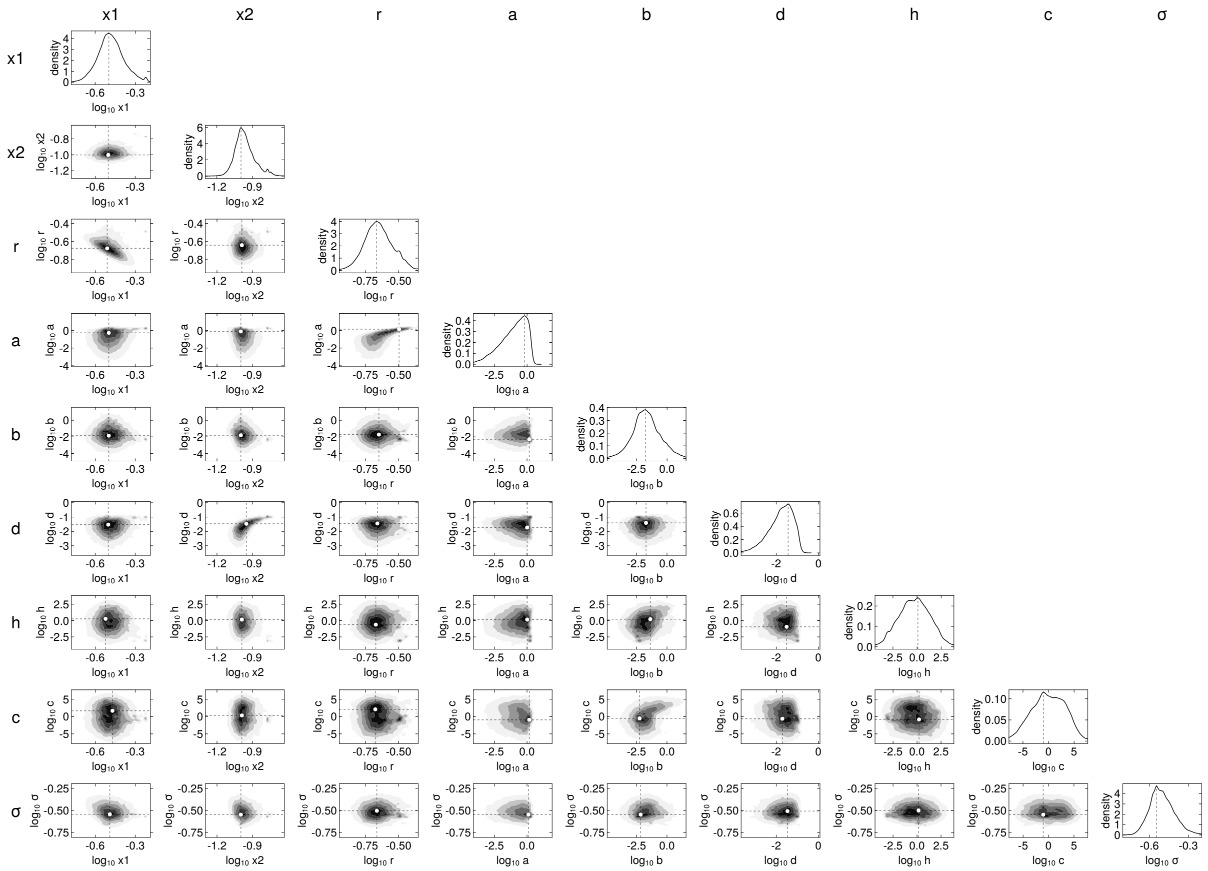

antigen_CD117_effector_uT_target_WT_model_Beddington_DeAngelis_activation_ratio_1_8.0

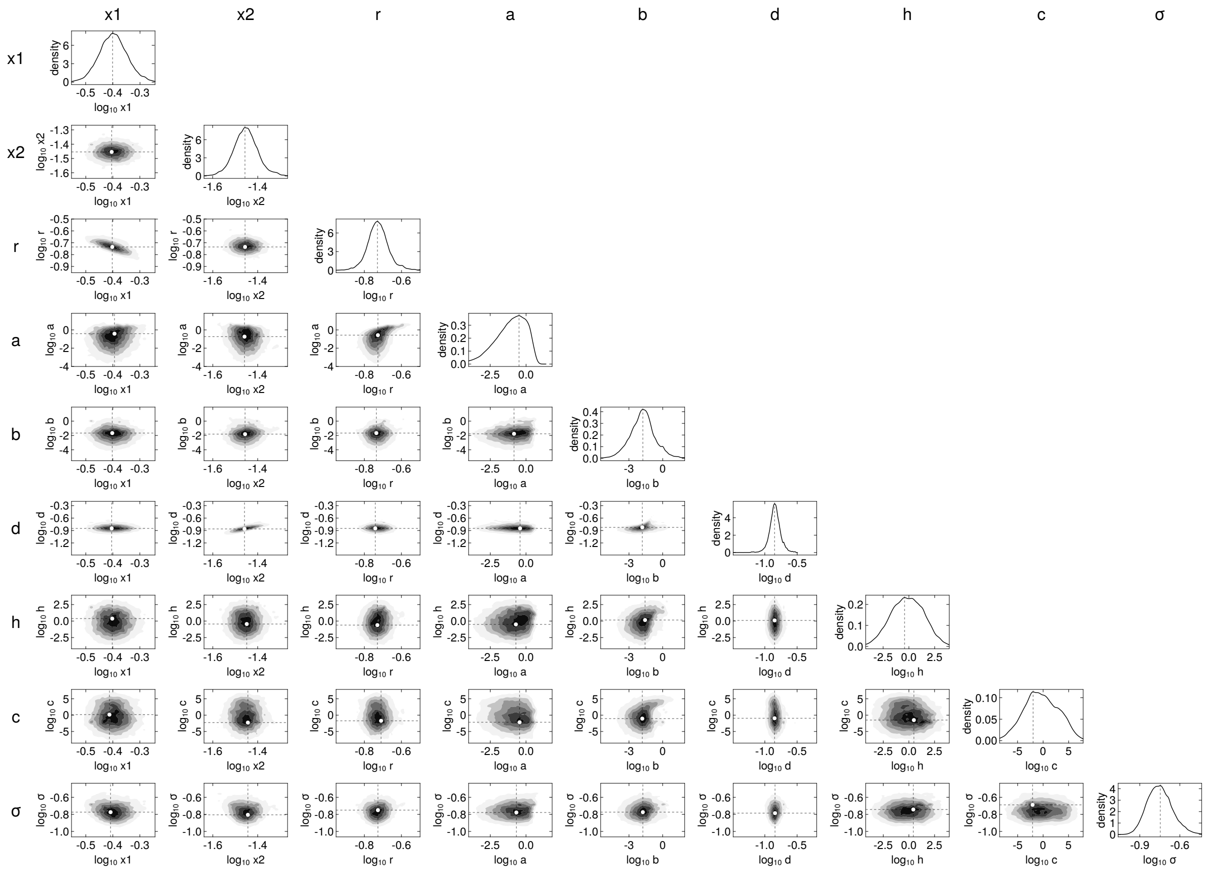


224

225

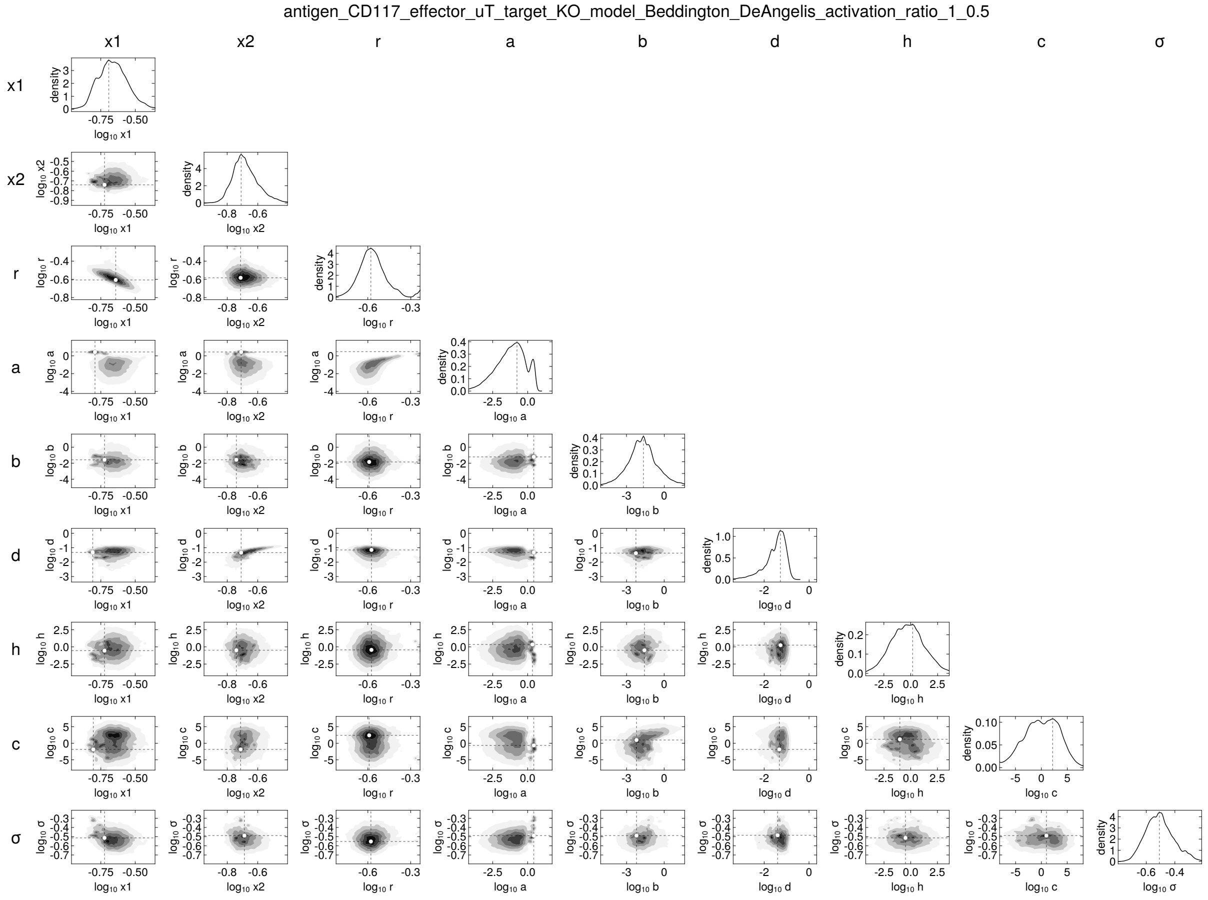
antigen_CD117_effector_uT_target_KO_model_Beddington_DeAngelis_activation_ratio_1_0.5
x1
x2
r
a
b
d
h
c
σ
density
log10 x1
log10 x2
log10 r
log10 a
log10 b
log10 d
log10 h
log10 c
log10 σ

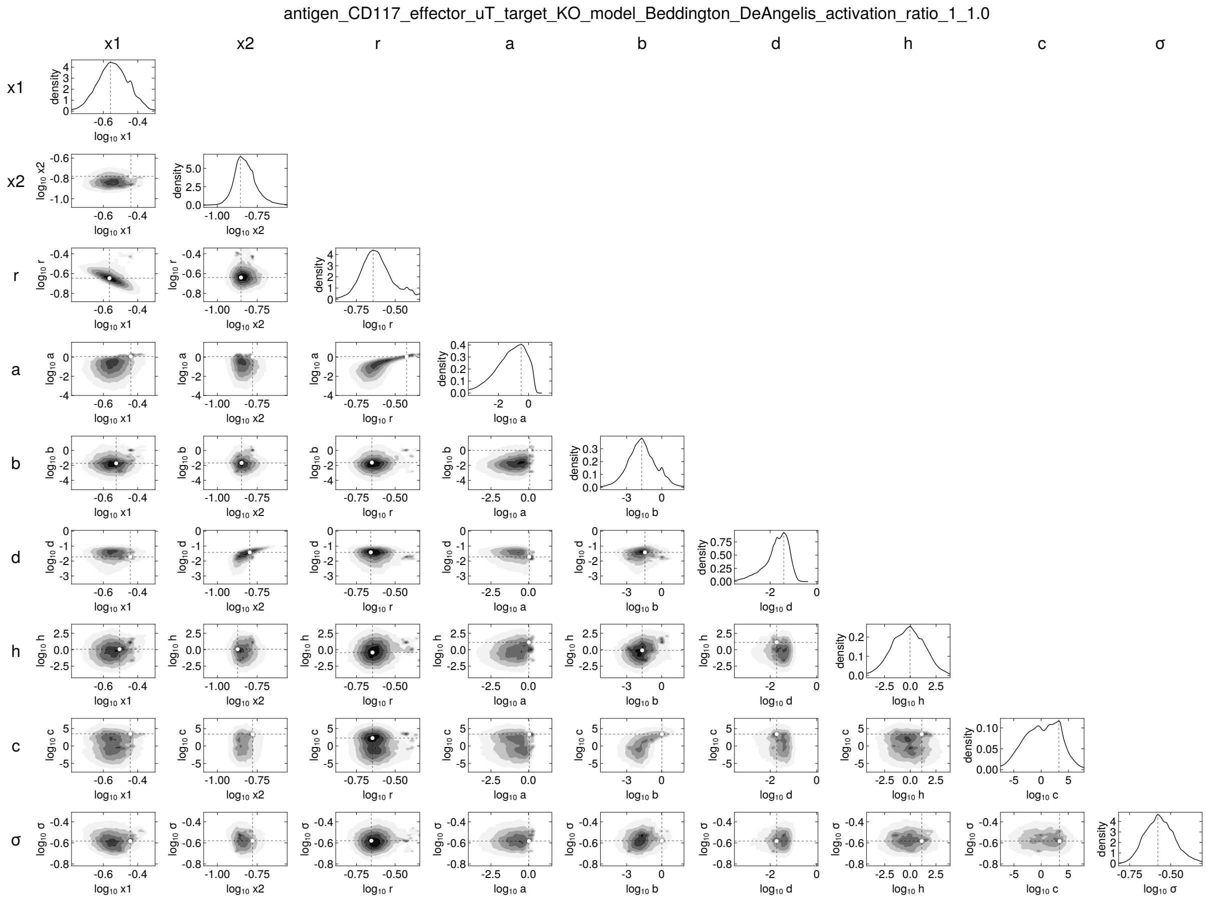
antigen_CD117_effector_uT_target_KO_model_Beddington_DeAngelis_activation_ratio_1_1.0
x1
x2
r
a
b
d
h
c
σ

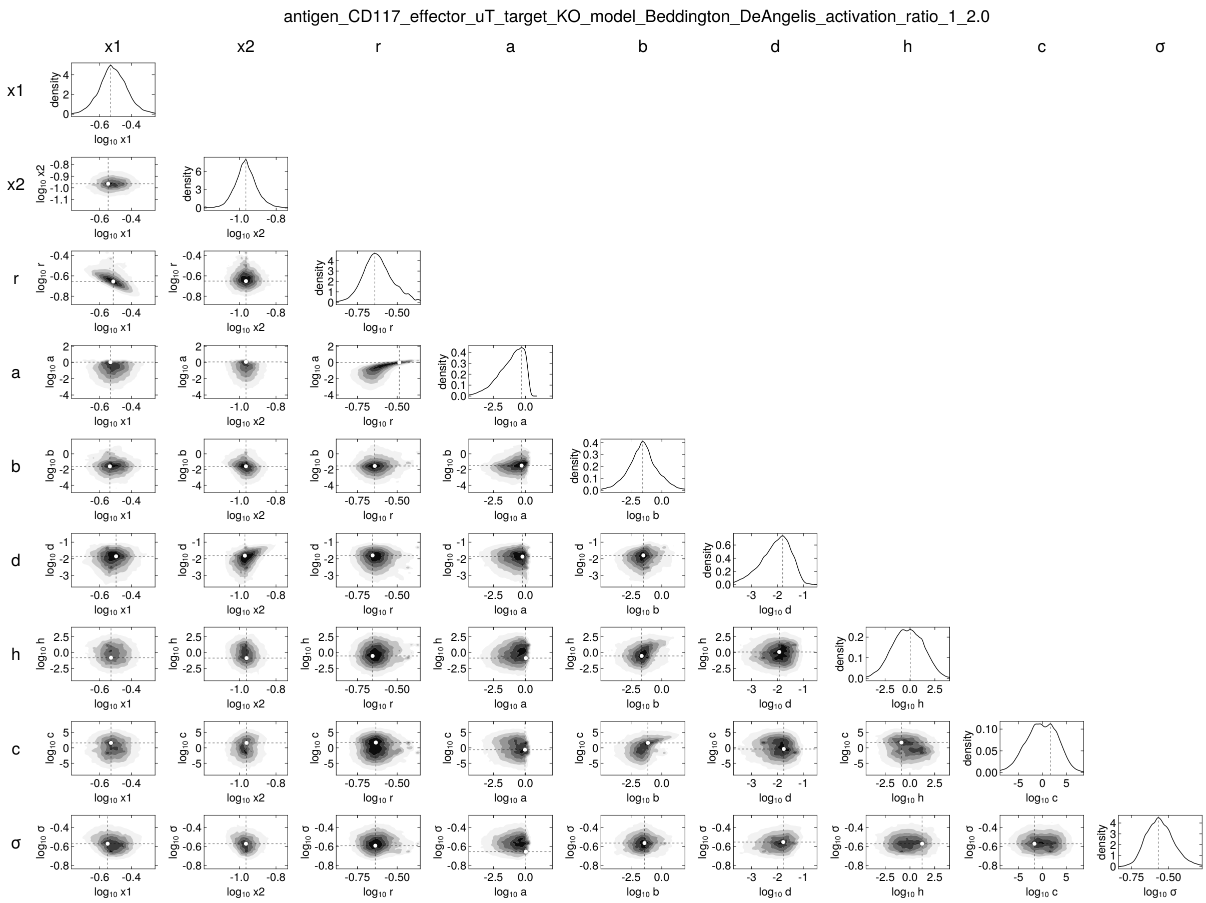
antigen_CD117_effector_uT_target_KO_model_Beddington_DeAngelis_activation_ratio_1_2.0
x1
x2
r
a
b
d
h
c
σ

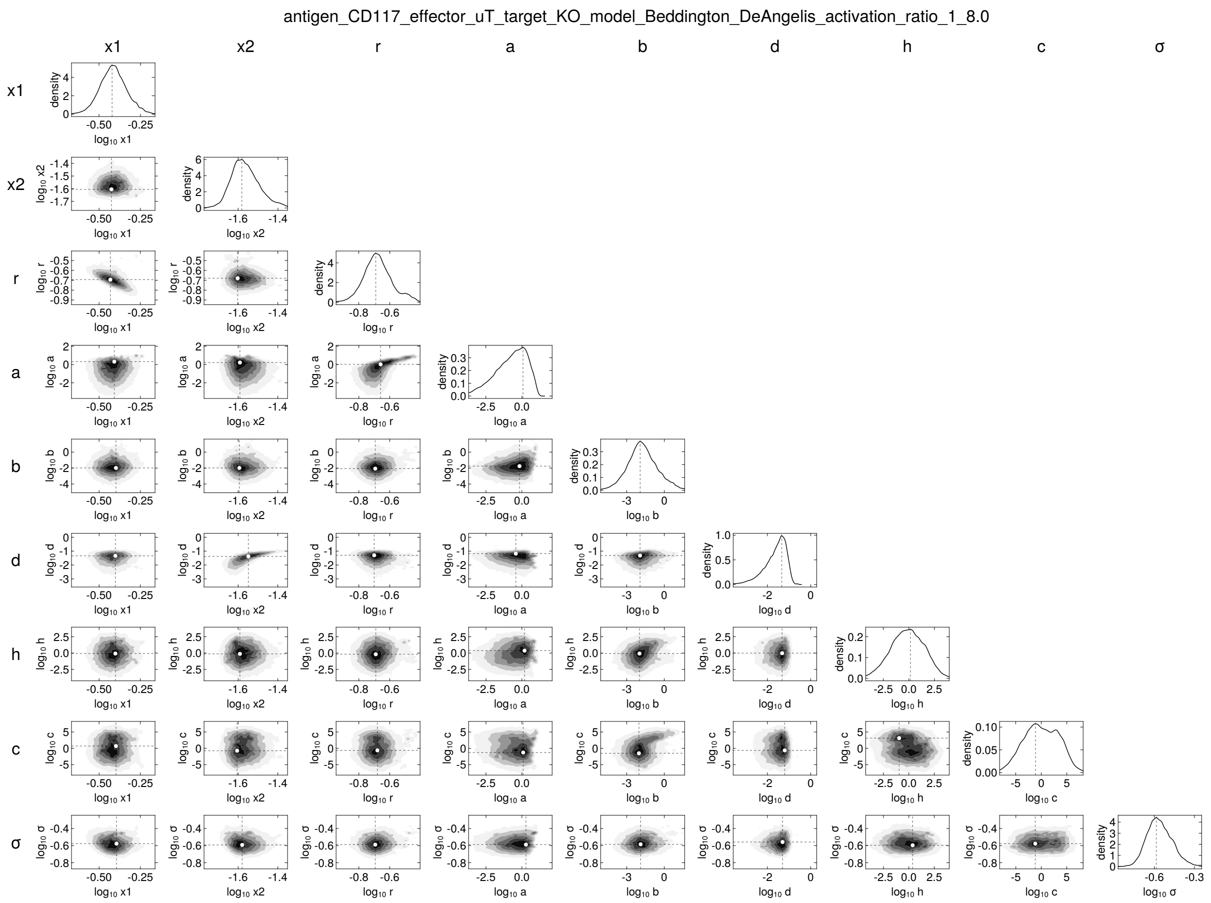
antigen_CD117_effector_uT_target_KO_model_Beddington_DeAngelis_activation_ratio_1_8.0
x1
x2
r
a
b
d
h
c
σ
density
$\log_{10}$ x1
$\log_{10}$ x2
$\log_{10}$ r
$\log_{10}$ a
$\log_{10}$ b
$\log_{10}$ d
$\log_{10}$ h
$\log_{10}$ c
$\log_{10}$ σ

antigen_CD117_effector_CART_target_WT_model_Beddington_DeAngelis_activation_ratio_1_0.5

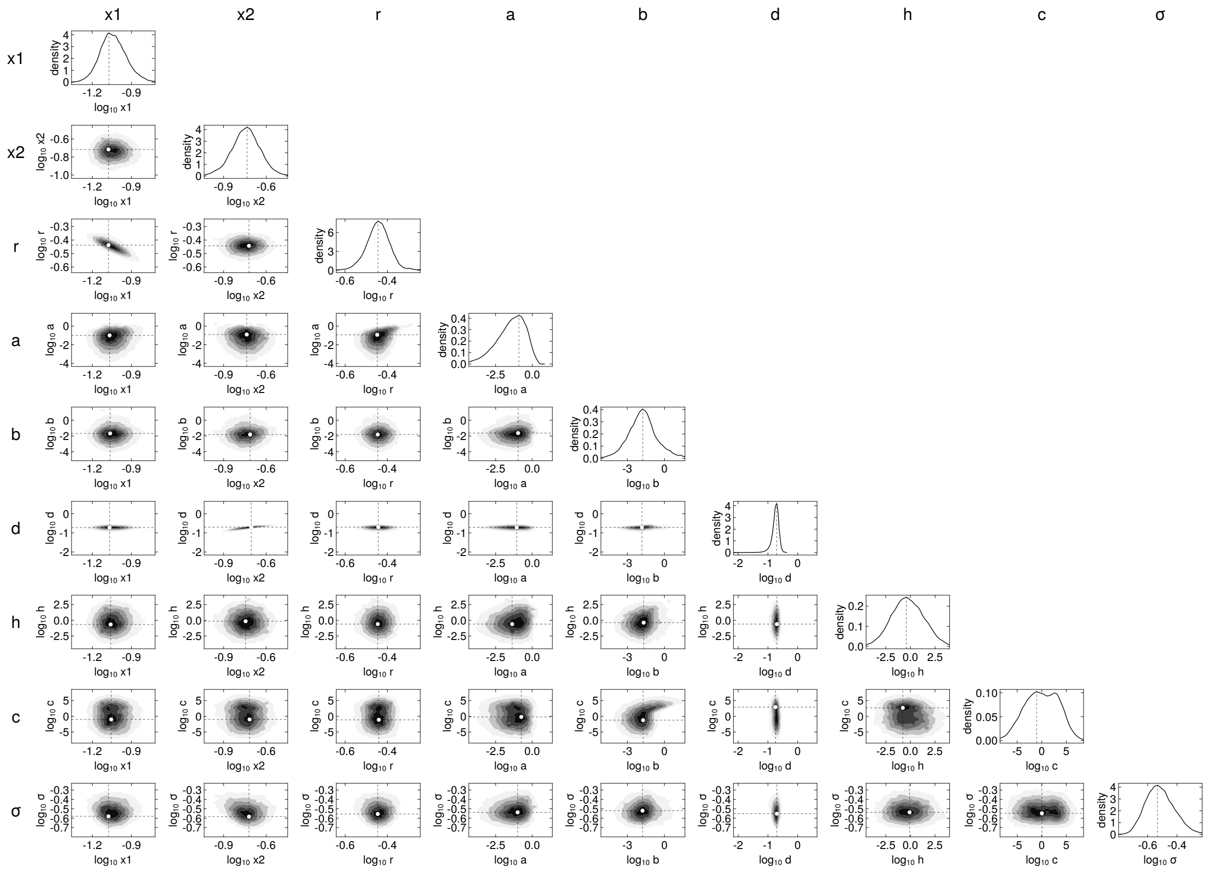

antigen_CD117_effector_CART_target_WT_model_Beddington_DeAngelis_activation_ratio_1_1.0

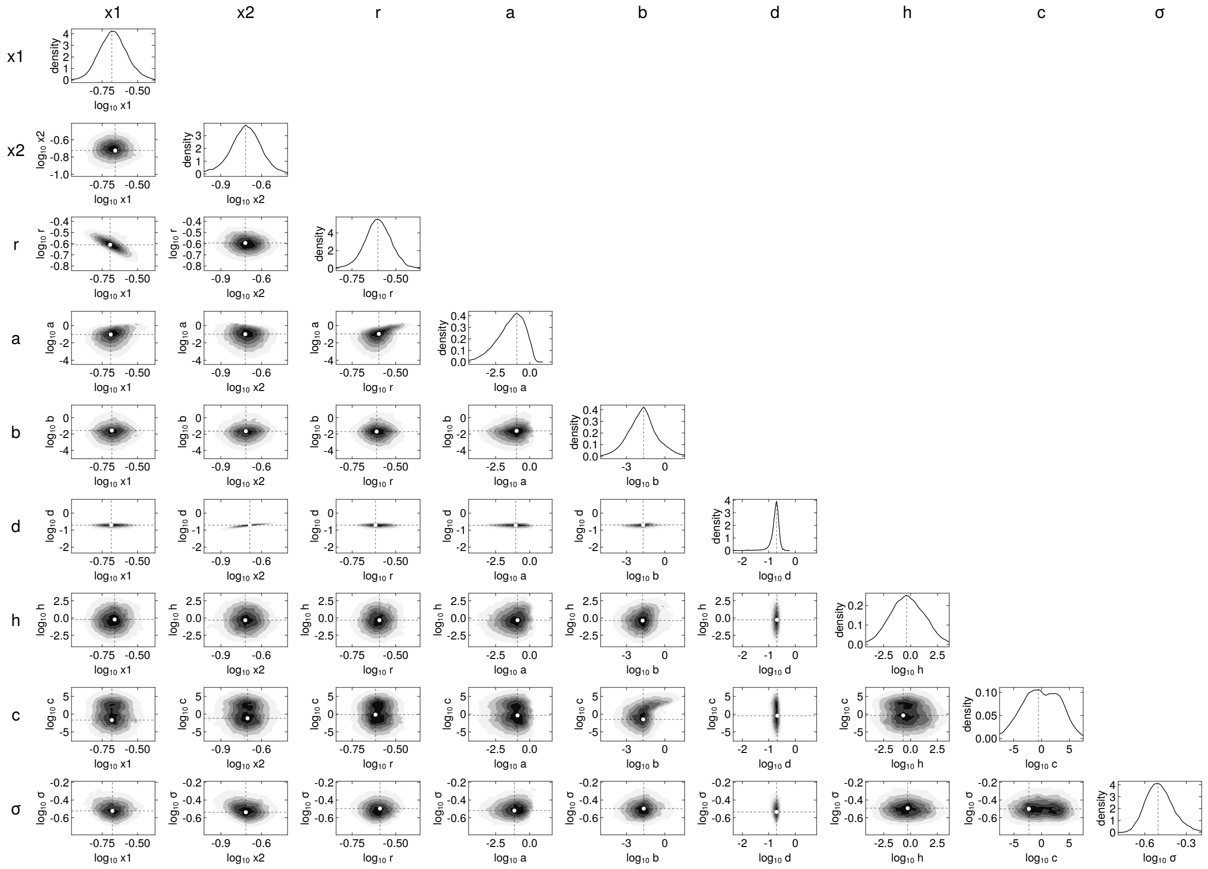


230

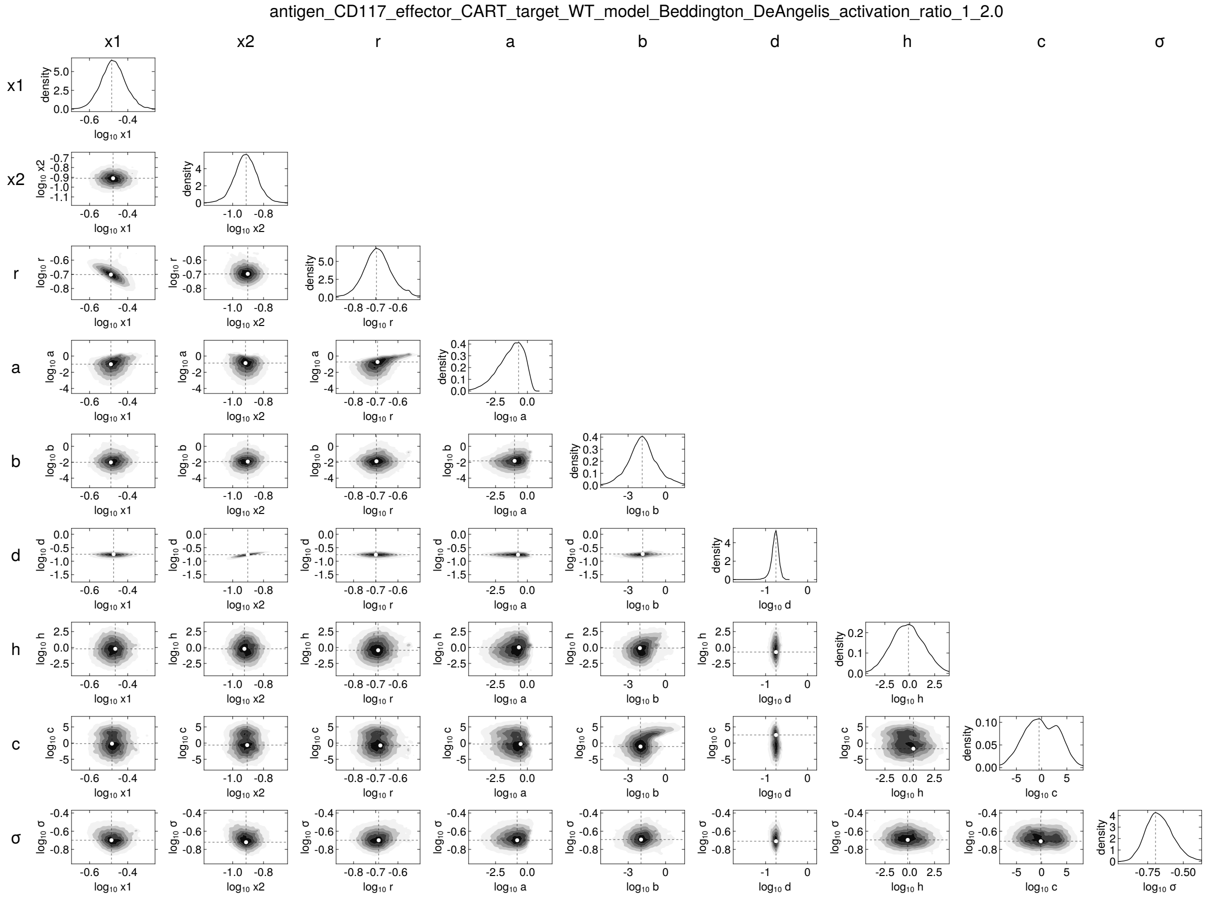
antigen_CD117_effector_CART_target_WT_model_Beddington_DeAngelis_activation_ratio_1_2.0
x1
x2
r
a
b
d
h
c
σ
density

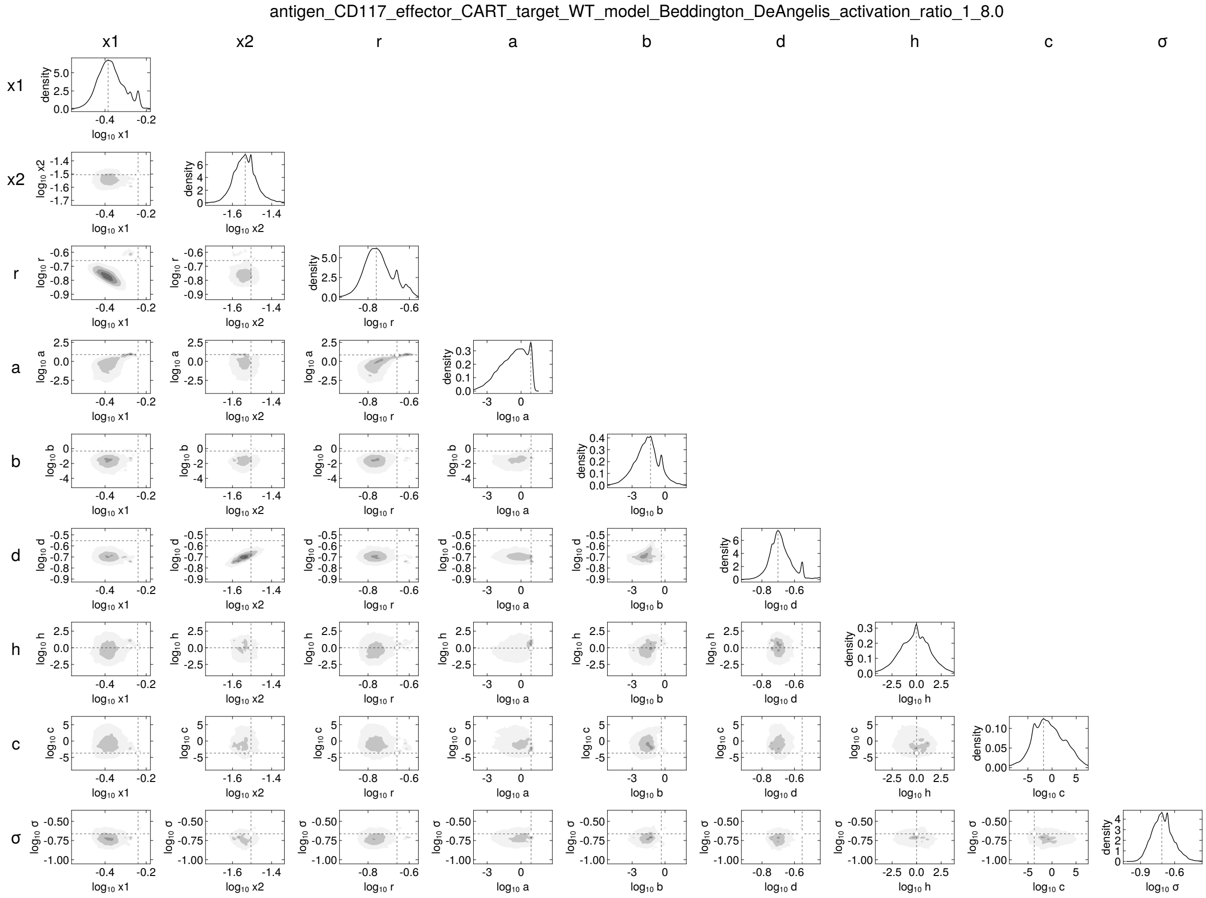
antigen_CD117_effector_CART_target_WT_model_Beddington_DeAngelis_activation_ratio_1_8.0
x1
x2
r
a
b
d
h
c
σ
density
$\log_{10}$ x1
$\log_{10}$ x2
$\log_{10}$ r
$\log_{10}$ a
$\log_{10}$ b
$\log_{10}$ d
$\log_{10}$ h
$\log_{10}$ c
$\log_{10}$ σ

232

antigen_CD117_effector_CART_target_KO_model_Beddington_DeAngelis_activation_ratio_1_0.5

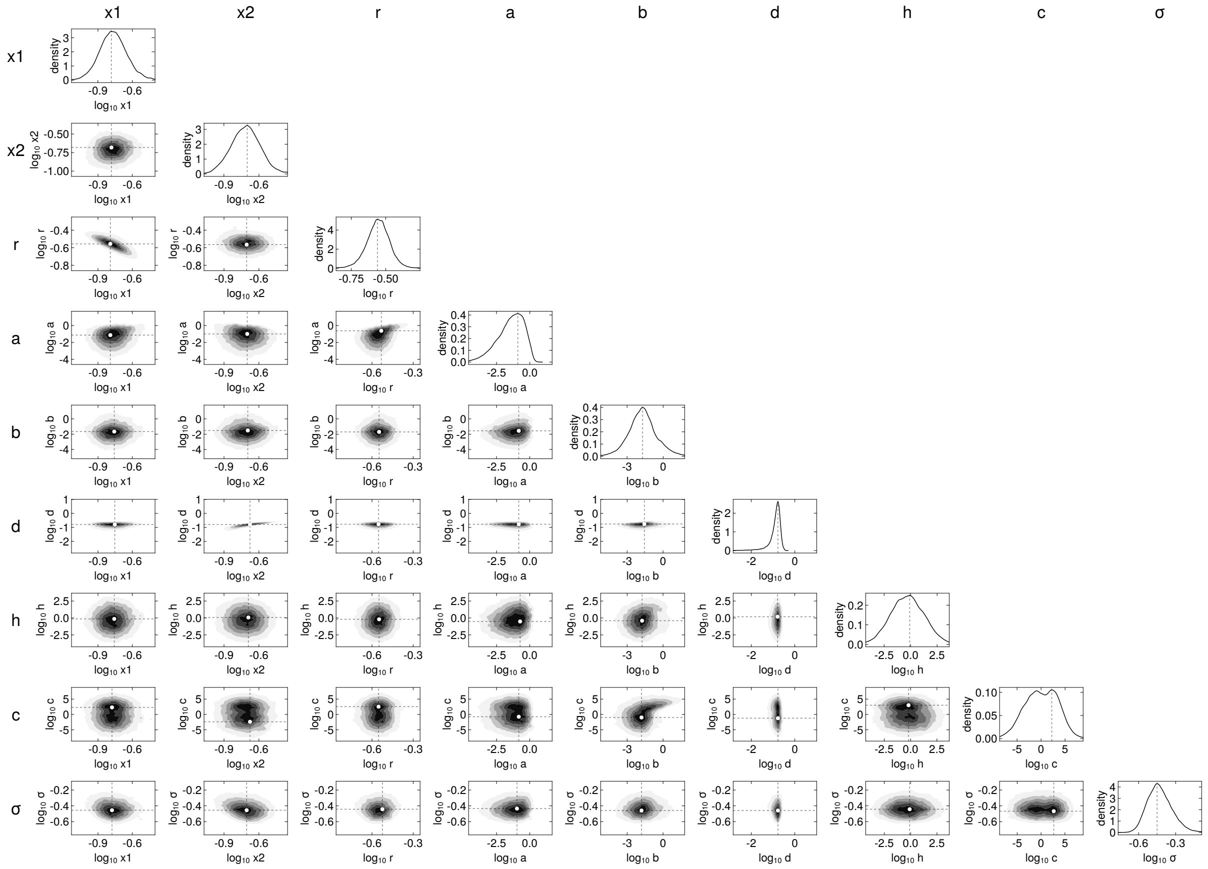

antigen_CD117_effector_CART_target_KO_model_Beddington_DeAngelis_activation_ratio_1_1.0

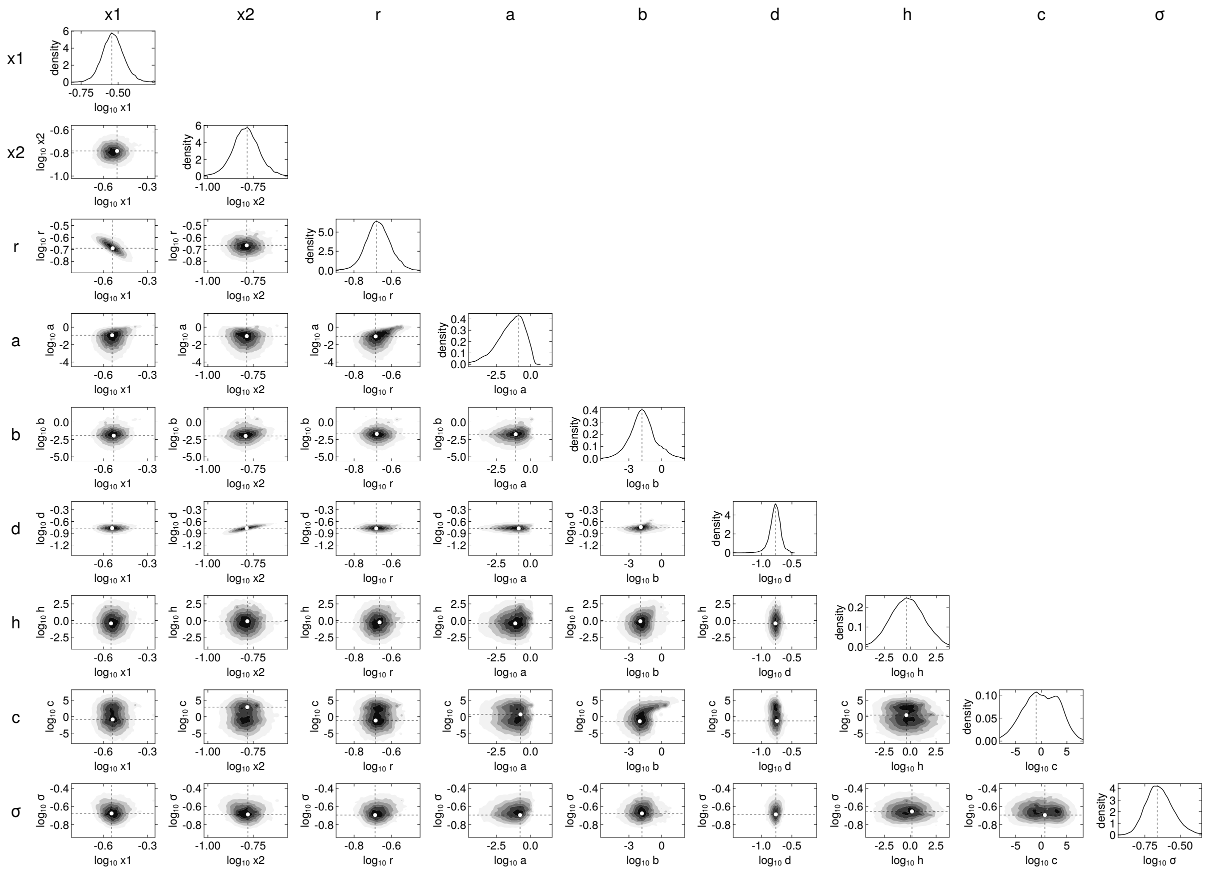


234

antigen_CD117_effector_CART_target_KO_model_Beddington_DeAngelis_activation_ratio_1_2.0

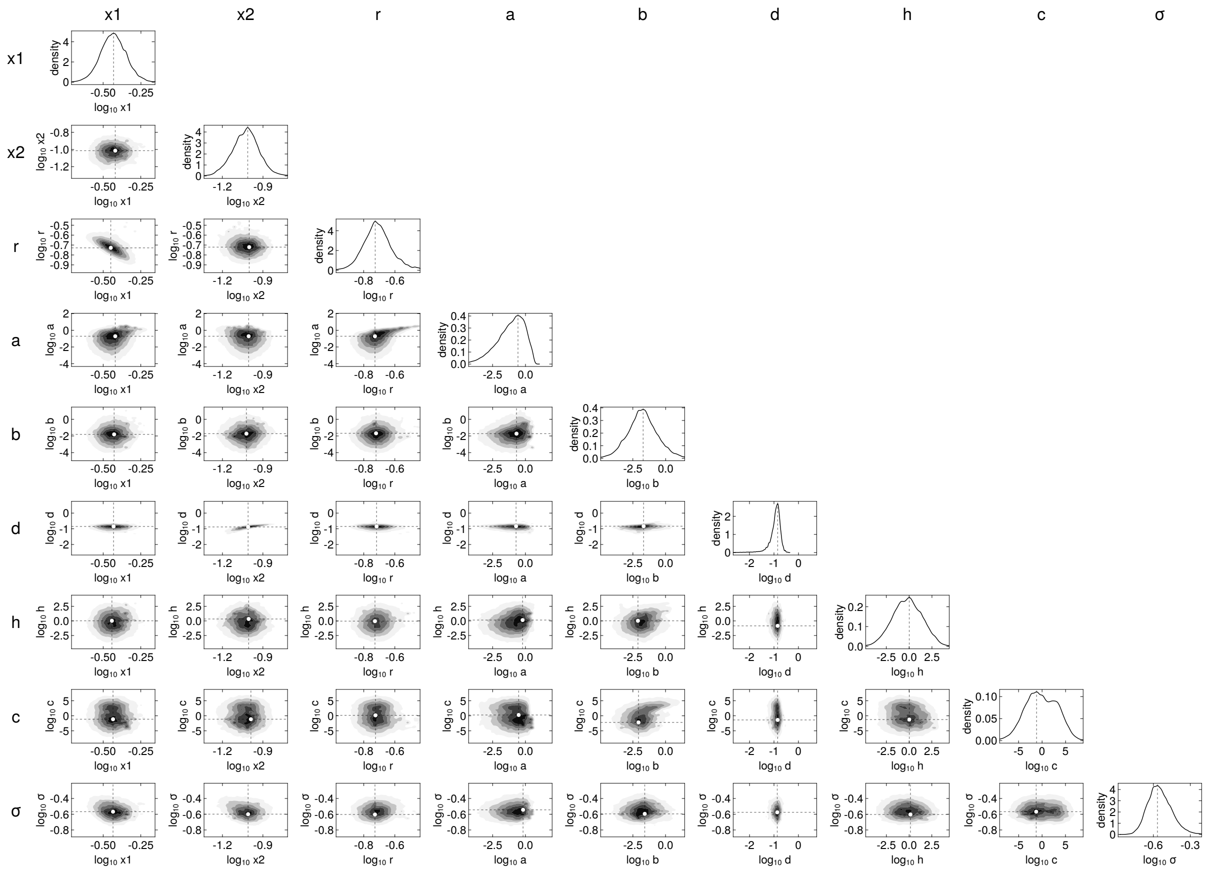

235

antigen_CD117_effector_CART_target_KO_model_Beddington_DeAngelis_activation_ratio_1_8.0

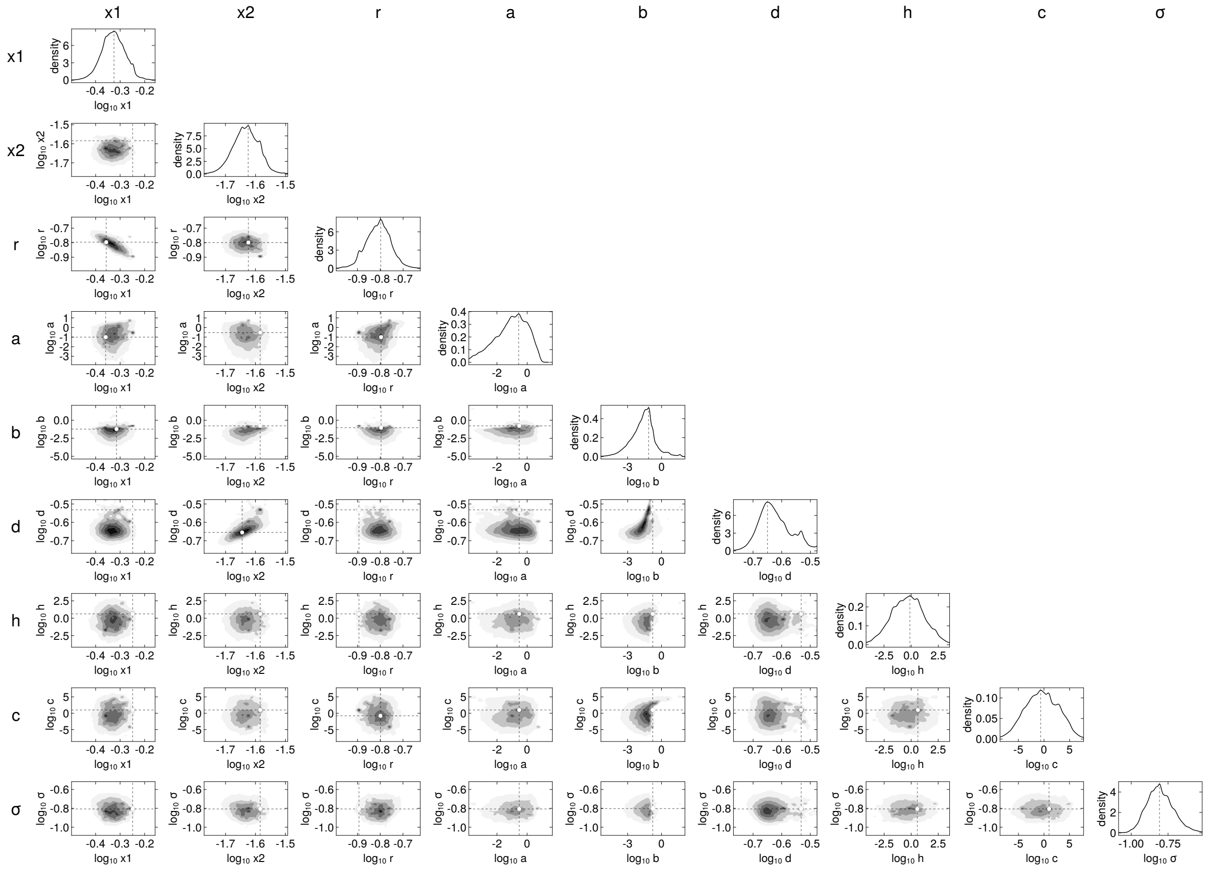


236

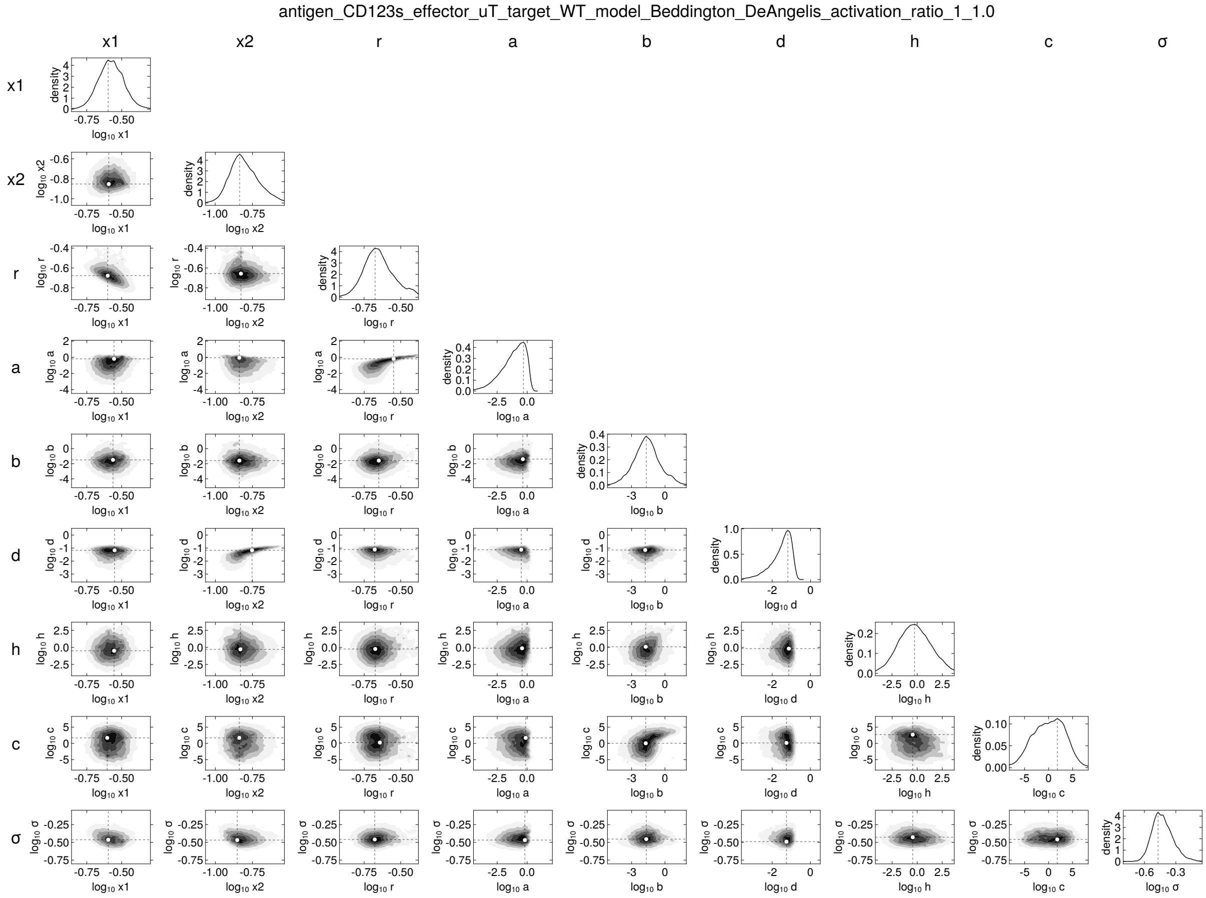
antigen_CD123s_effector_uT_target_WT_model_Beddington_DeAngelis_activation_ratio_1_1.0
x1
x2
r
a
b
d
h
c
σ

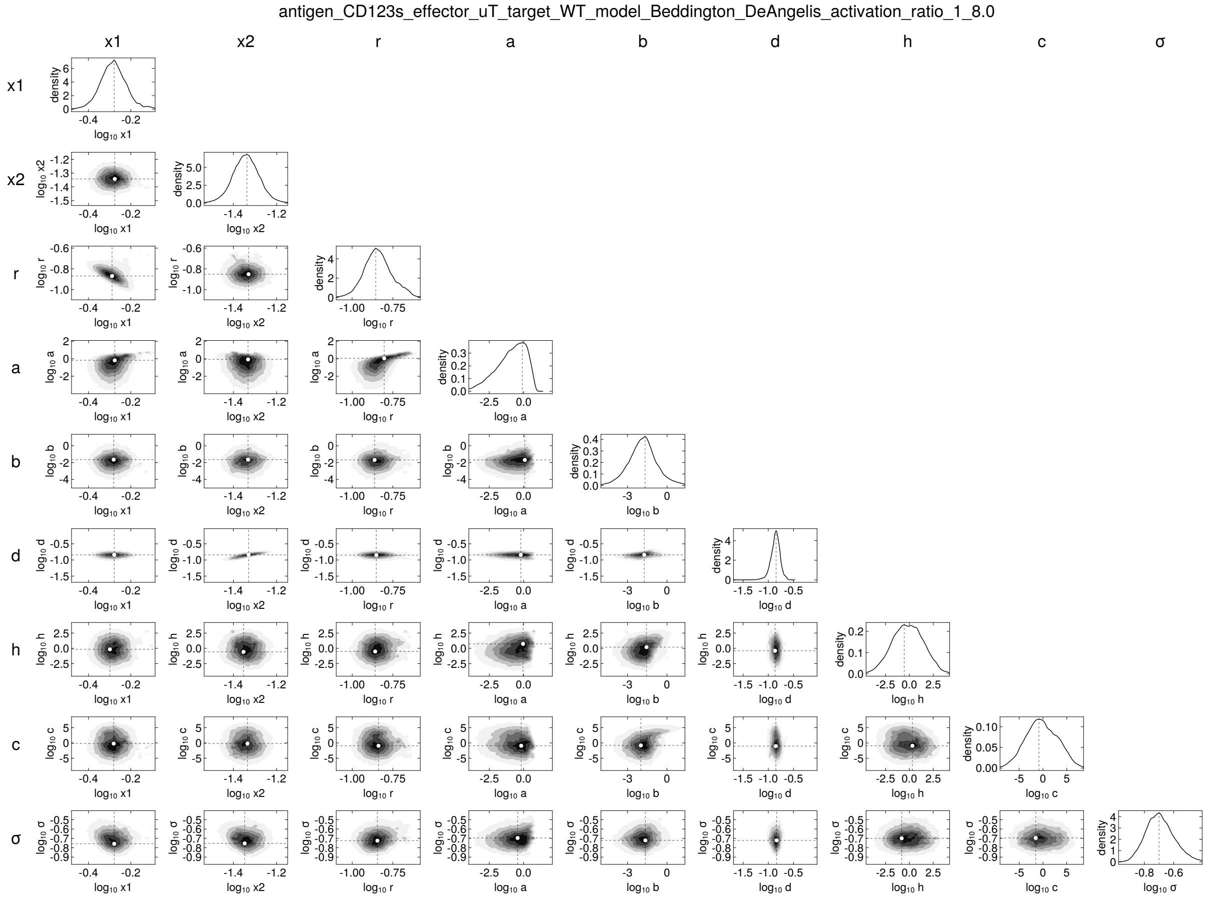
antigen_CD123s_effector_uT_target_WT_model_Beddington_DeAngelis_activation_ratio_1_8.0
x1
x2
r
a
b
d
h
c
σ

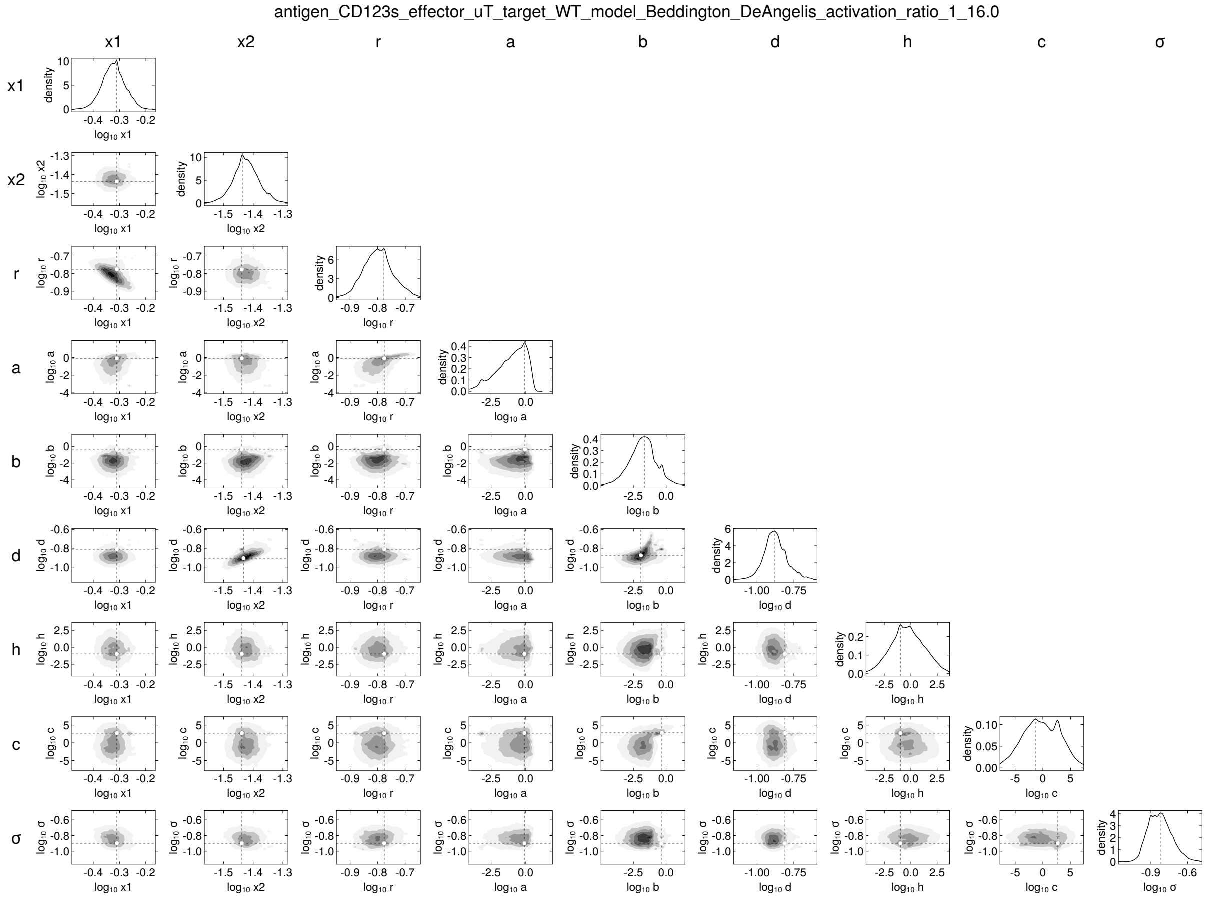
antigen_CD123s_effector_uT_target_WT_model_Beddington_DeAngelis_activation_ratio_1_16.0
x1
x2
r
a
b
d
h
c
σ
density
log10 x1
log10 x2
log10 r
log10 a
log10 b
log10 d
log10 h
log10 c
log10 σ

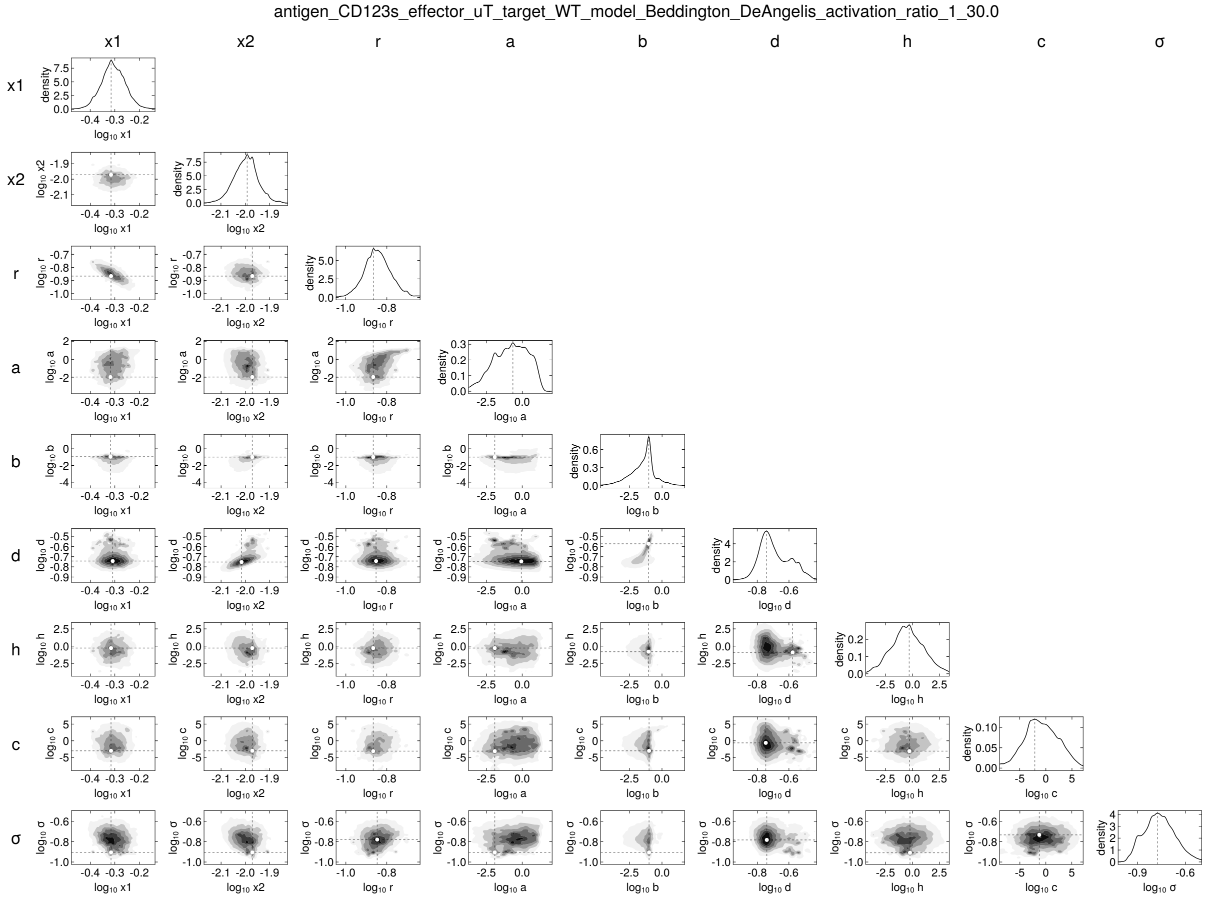
antigen_CD123s_effector_uT_target_WT_model_Beddington_DeAngelis_activation_ratio_1_30.0
x1
x2
r
a
b
d
h
c
σ


240

antigen_CD123s_effector_uT_target_KO_model_Beddington_DeAngelis_activation_ratio_1_1.0

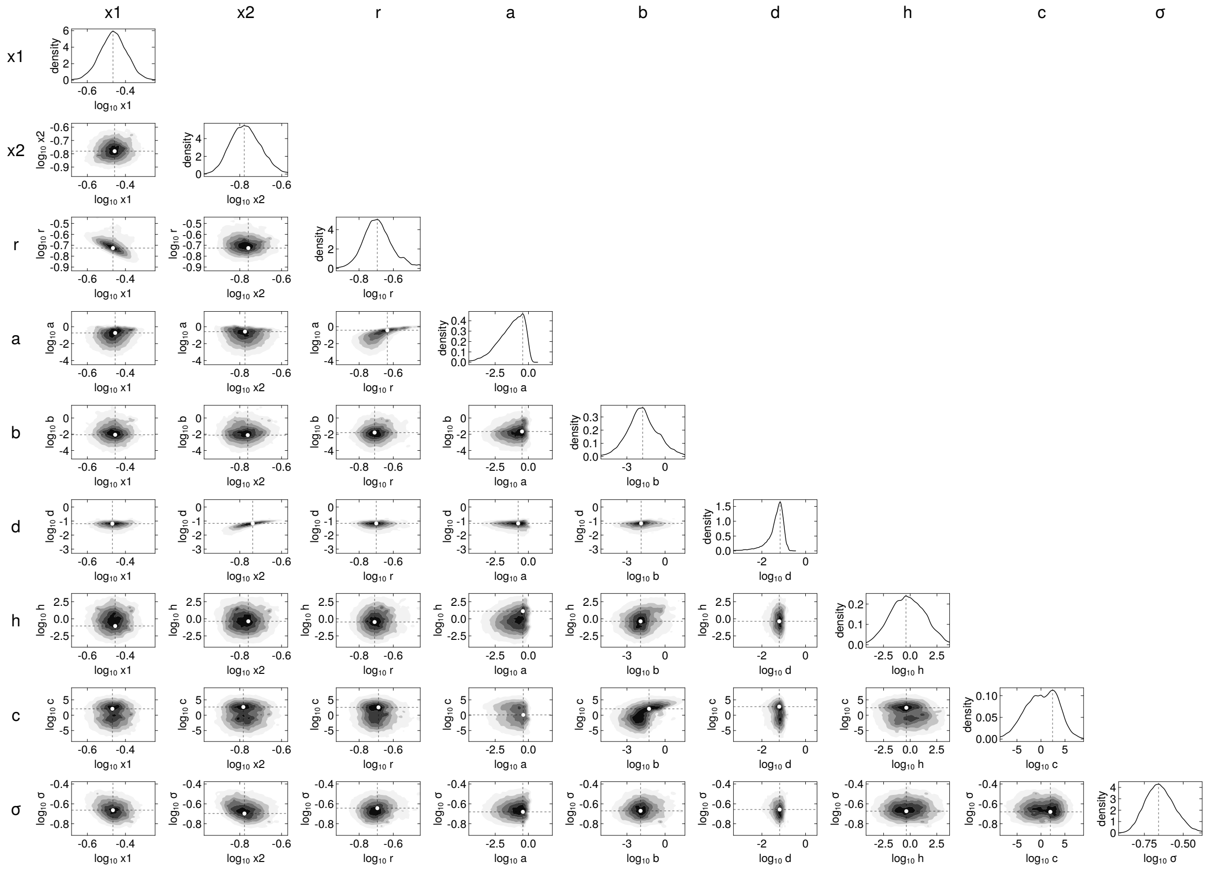

antigen_CD123s_effector_uT_target_KO_model_Beddington_DeAngelis_activation_ratio_1_8.0

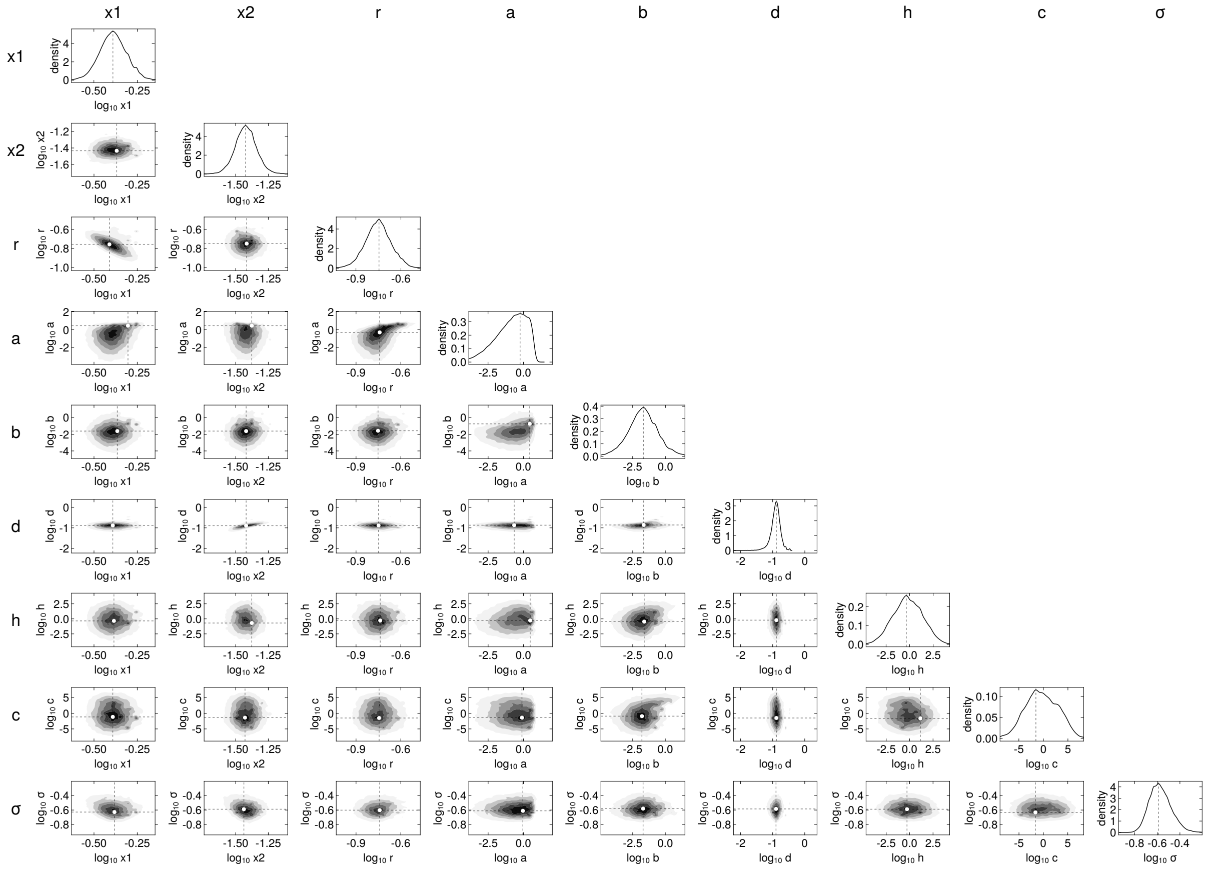

antigen_CD123s_effector_uT_target_KO_model_Beddington_DeAngelis_activation_ratio_1_16.0

243 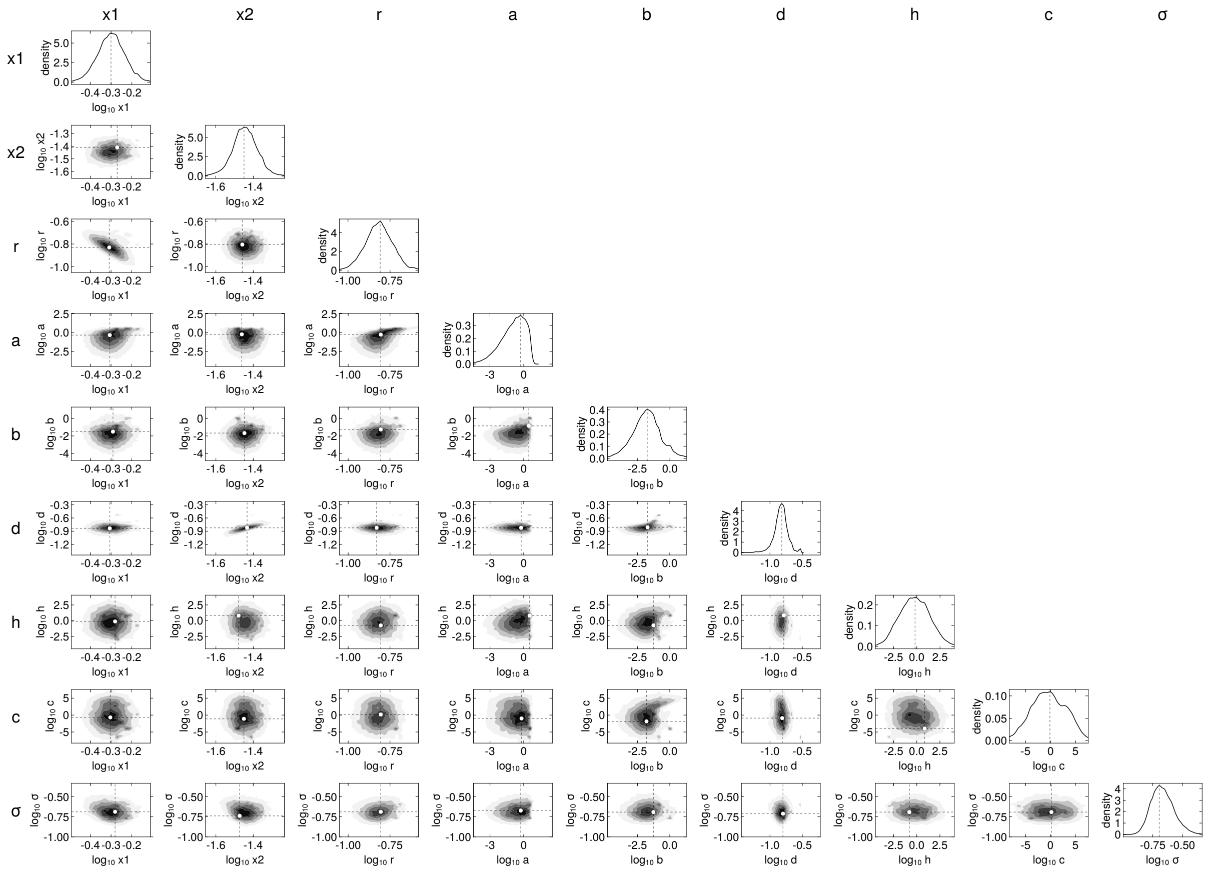

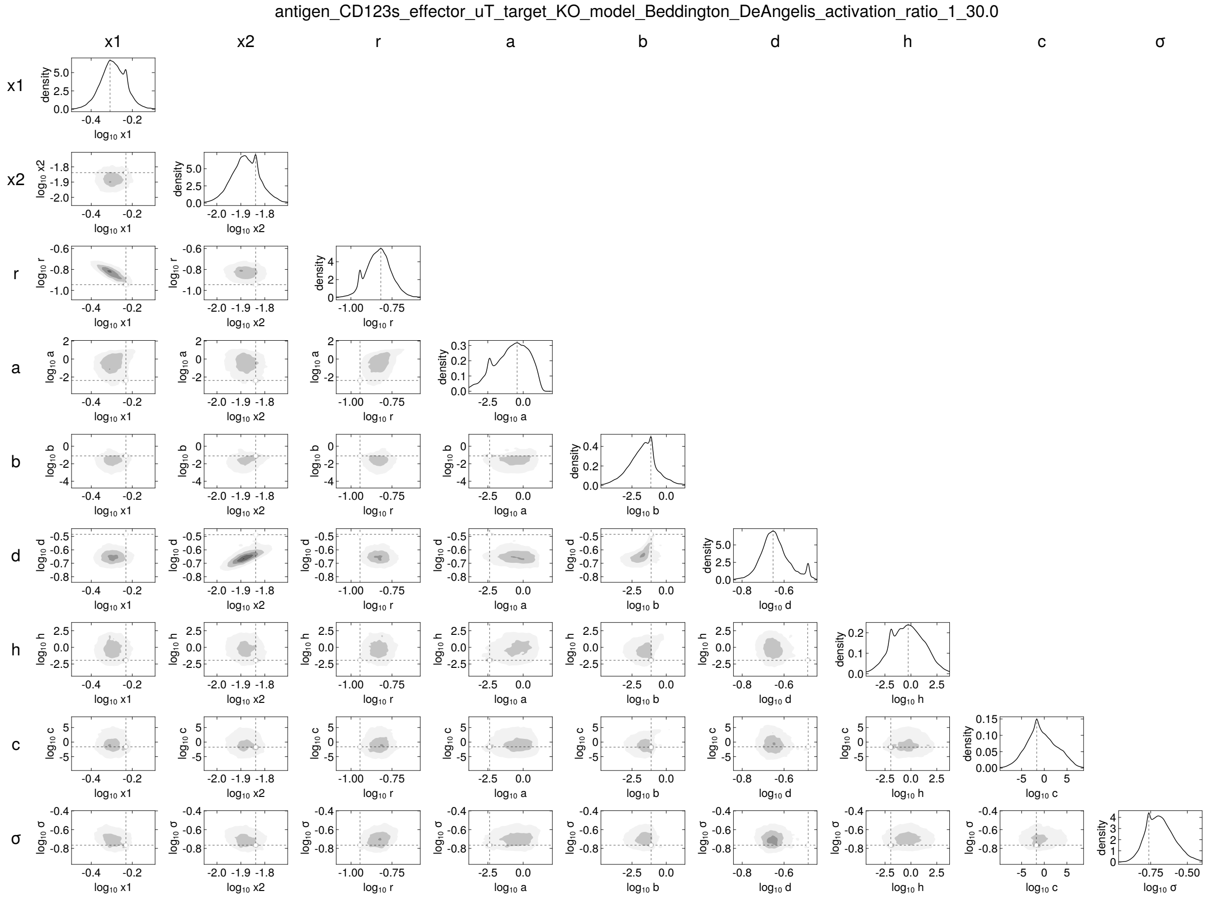
antigen_CD123s_effector_uT_target_KO_model_Beddington_DeAngelis_activation_ratio_1_30.0
x1
x2
r
a
b
d
h
c
σ
density

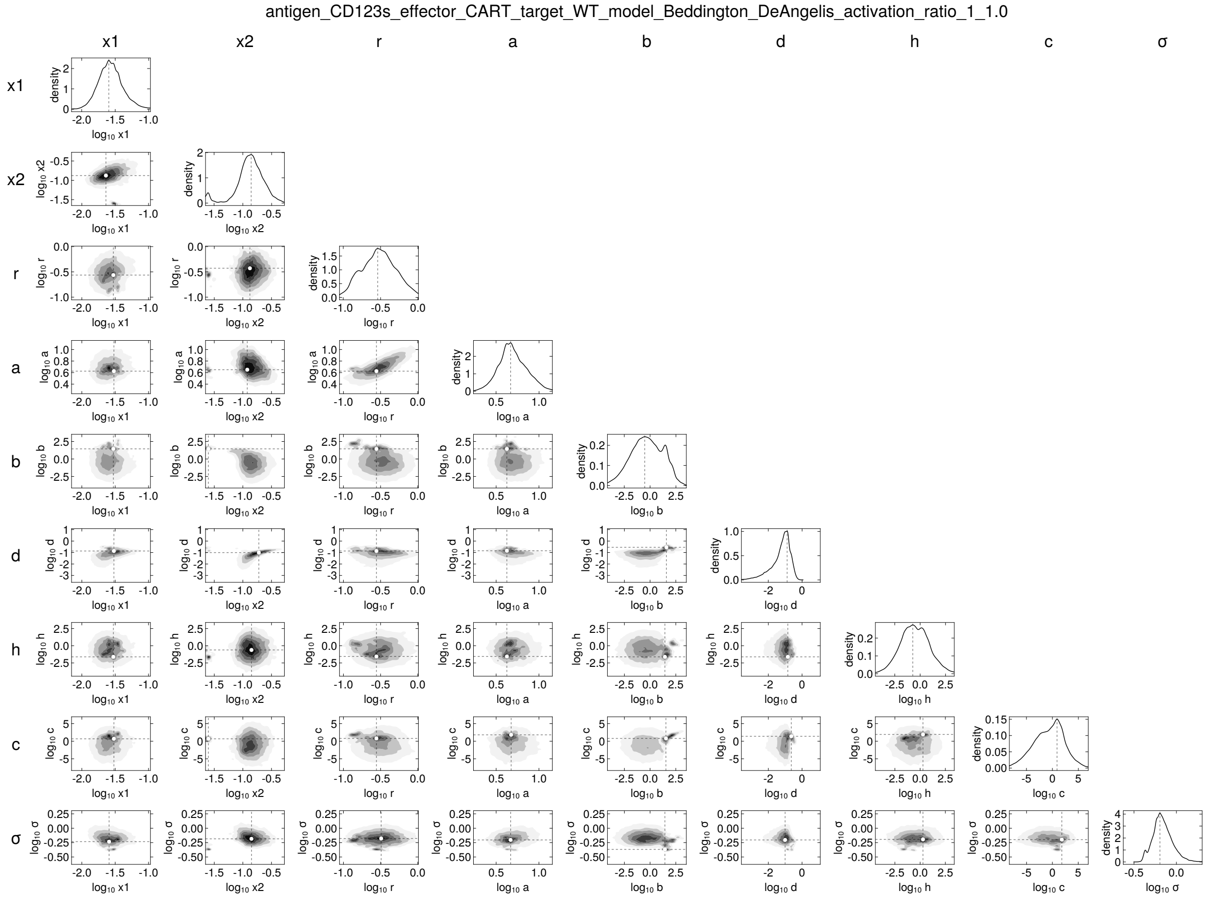
antigen_CD123s_effector_CART_target_WT_model_Beddington_DeAngelis_activation_ratio_1_1.0
x1
x2
r
a
b
d
h
c
σ
density
log10 x1
log10 x2
log10 r
log10 a
log10 b
log10 d
log10 h
log10 c
log10 σ

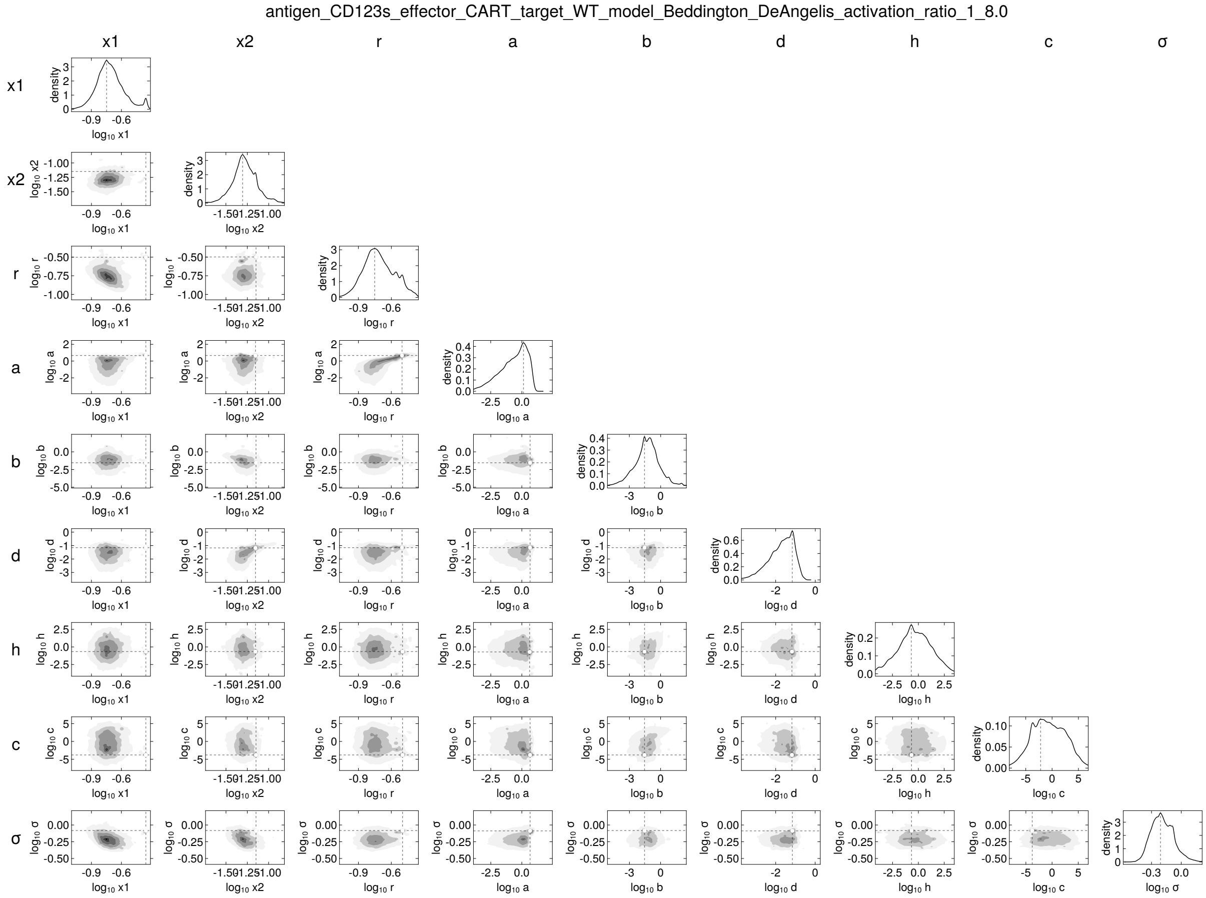
antigen_CD123s_effector_CART_target_WT_model_Beddington_DeAngelis_activation_ratio_1_8.0
x1
x2
r
a
b
d
h
c
σ

antigen_CD123s_effector_CART_target_WT_model_Beddington_DeAngelis_activation_ratio_1_16.0

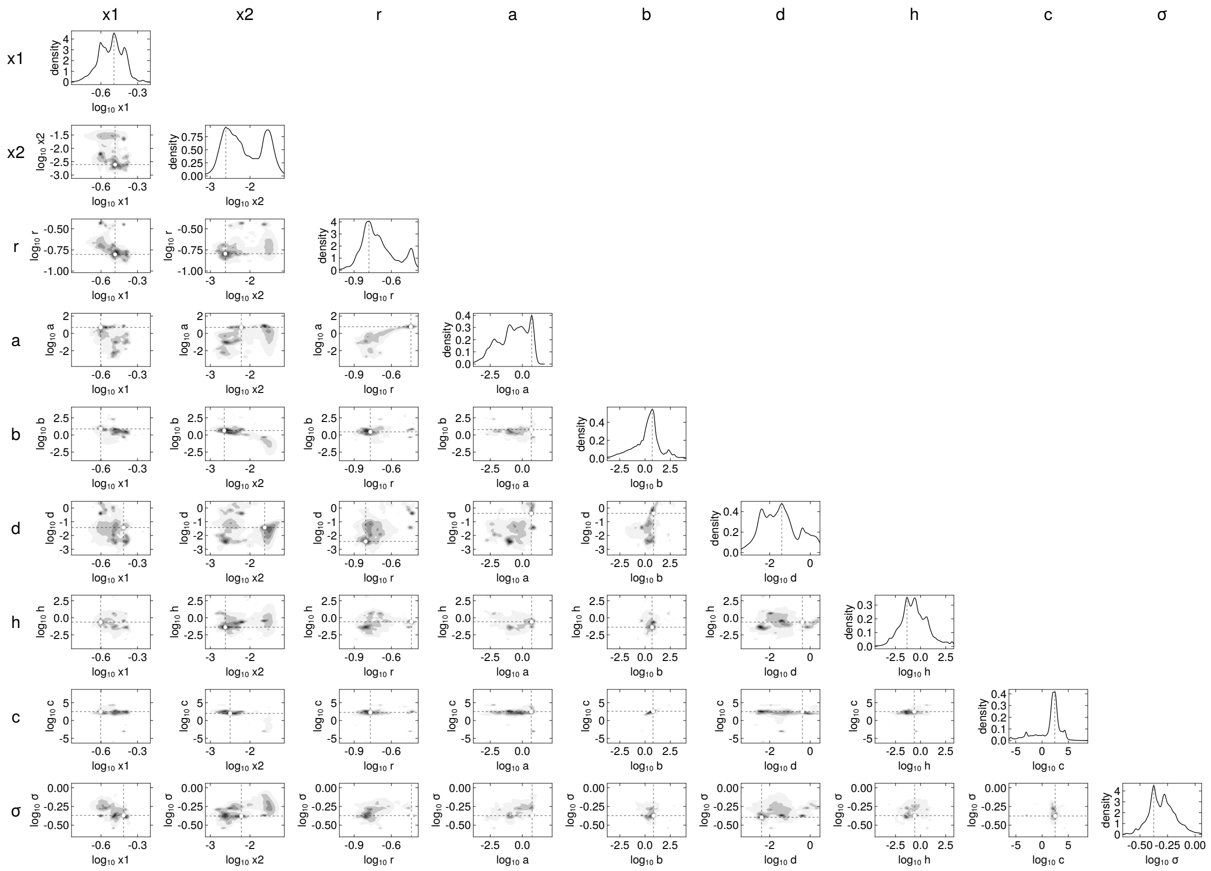


247

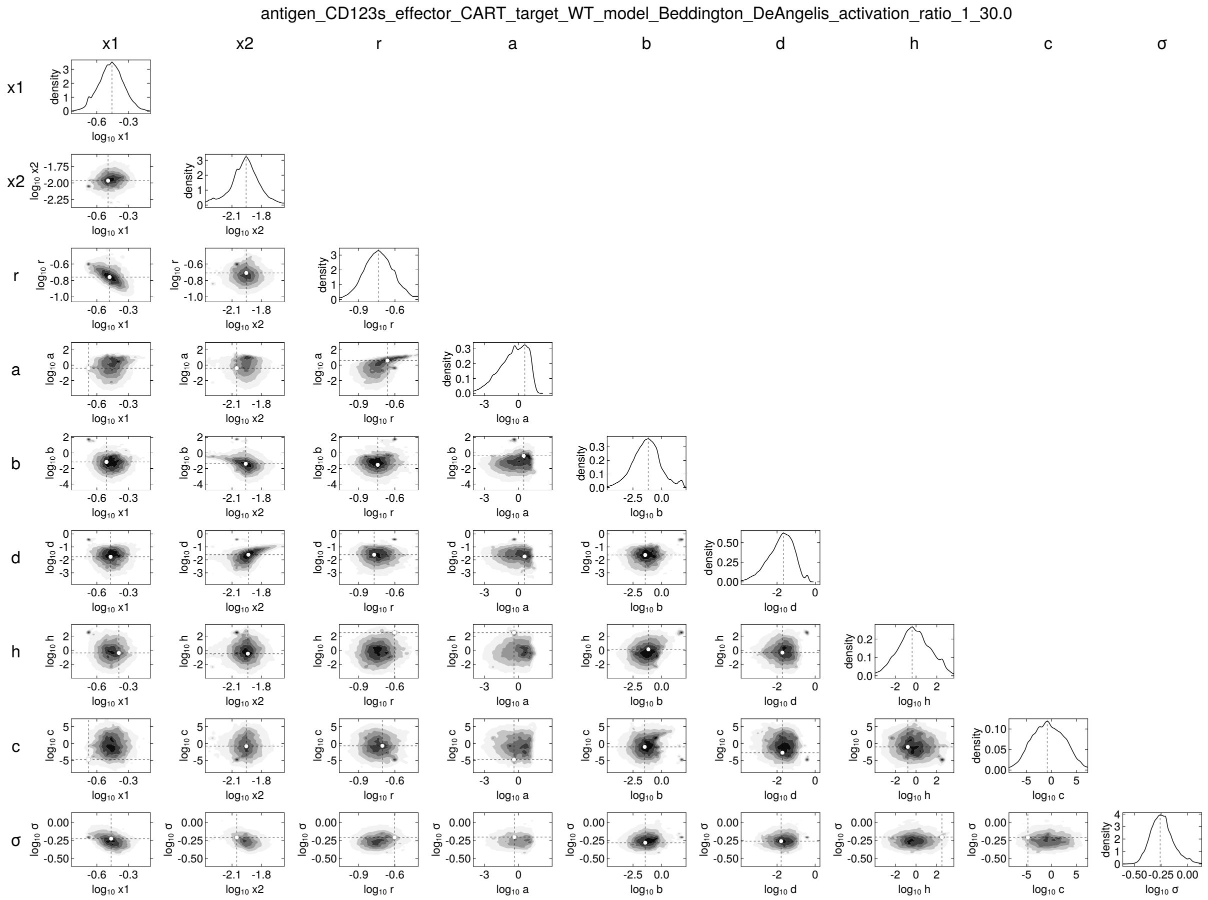
antigen_CD123s_effector_CART_target_WT_model_Beddington_DeAngelis_activation_ratio_1_30.0
x1
x2
r
a
b
d
h
c
σ
density
log10 x1
log10 x2
log10 r
log10 a
log10 b
log10 d
log10 h
log10 c
log10 σ

248

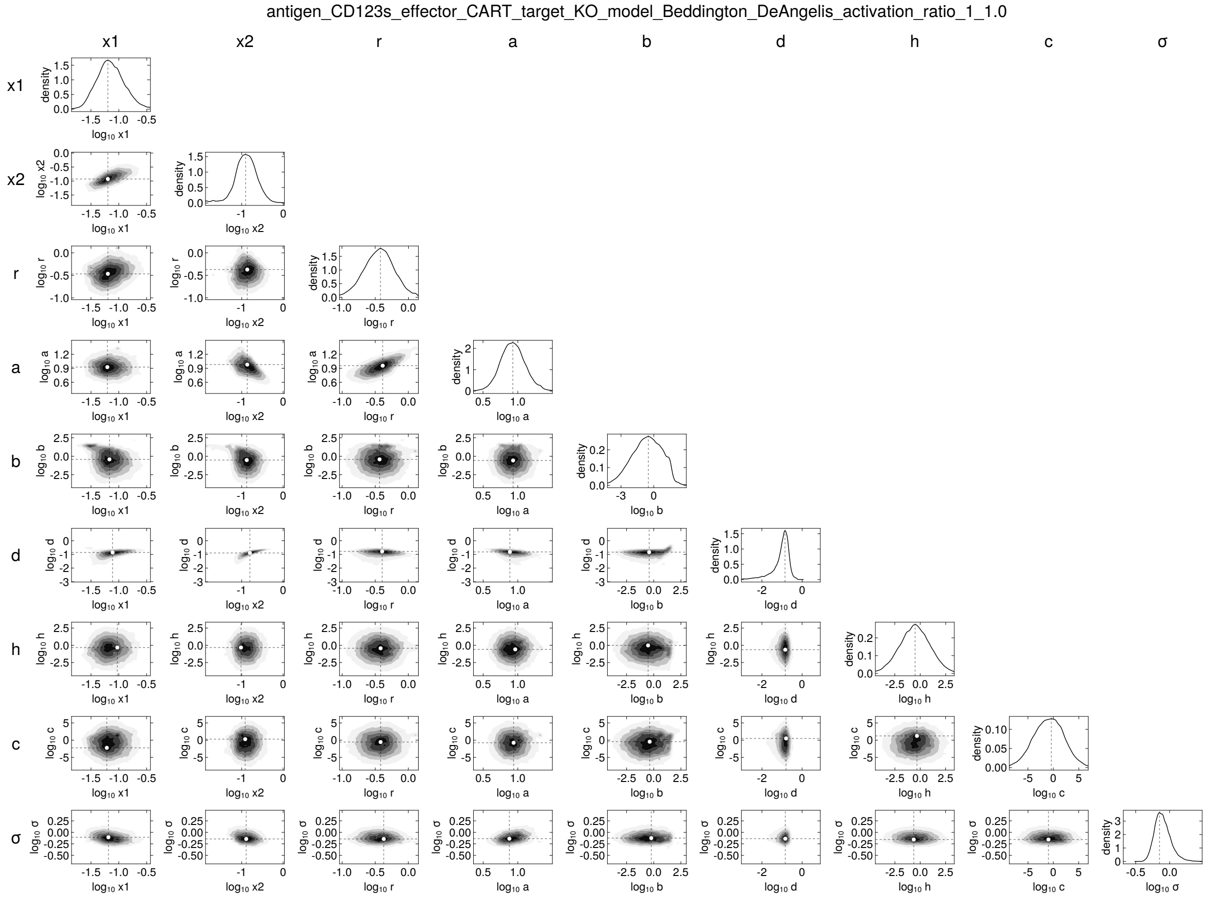

antigen_CD123s_effector_CART_target_KO_model_Beddington_DeAngelis_activation_ratio_1_1.0
x1
x2
r
a
b
d
h
c
σ

antigen_CD123s_effector_CART_target_KO_model_Beddington_DeAngelis_activation_ratio_1_8.0

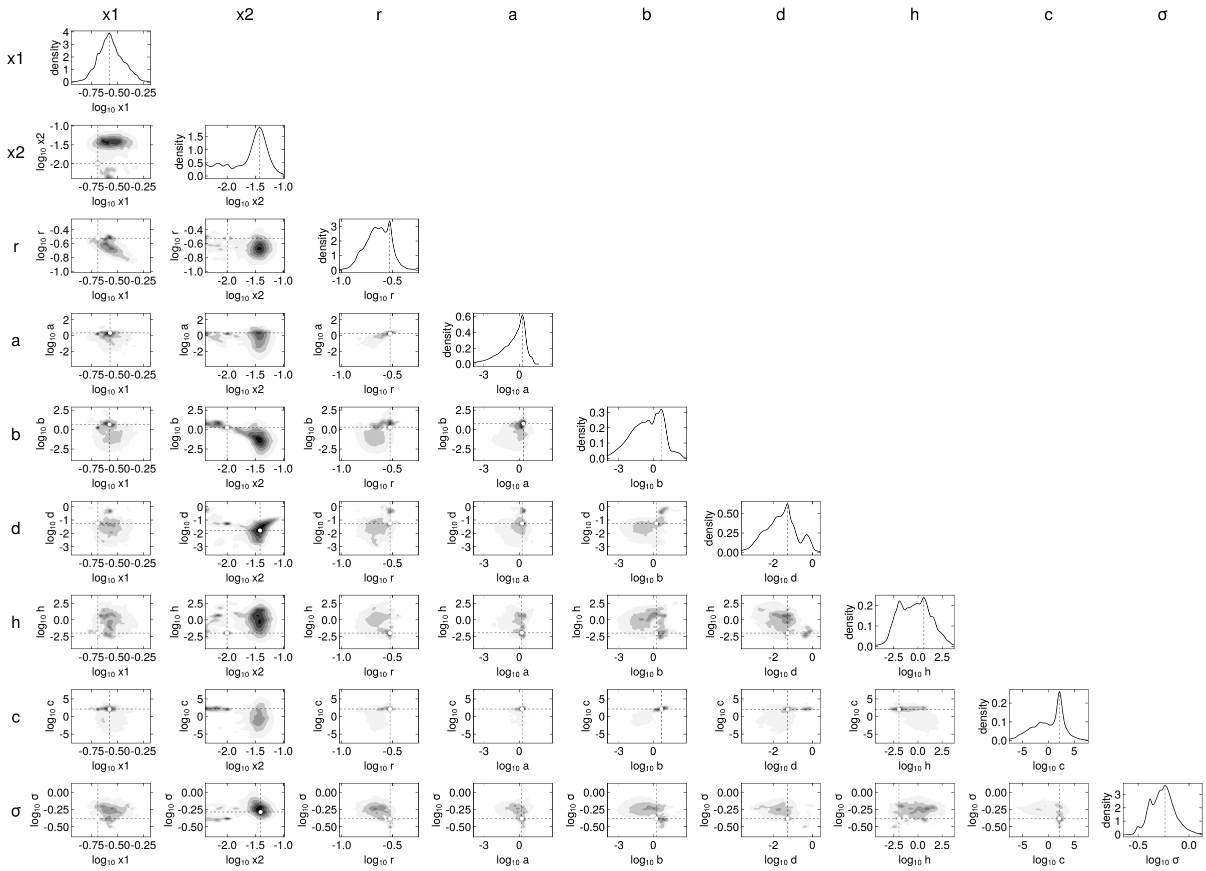


250

antigen_CD123s_effector_CART_target_KO_model_Beddington_DeAngelis_activation_ratio_1_16.0

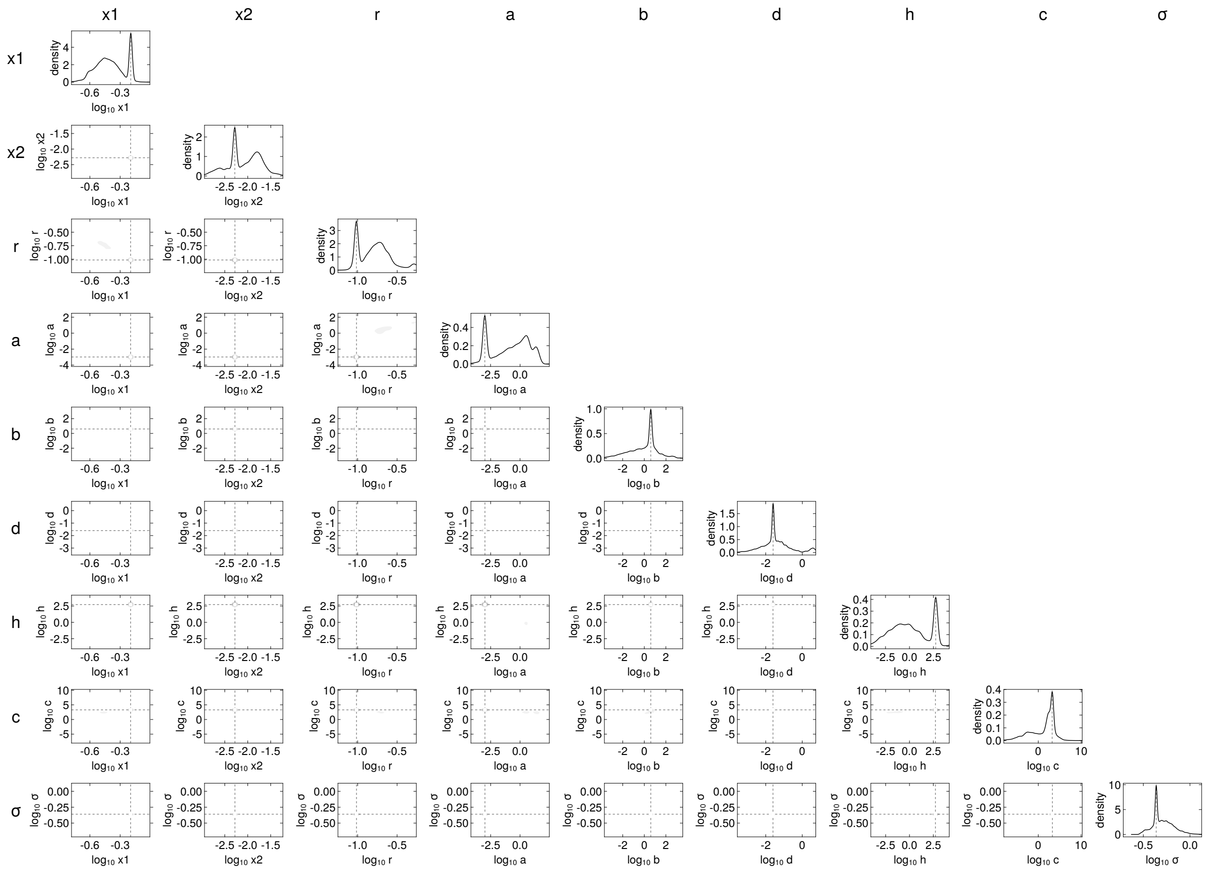


251

antigen_CD123s_effector_CART_target_KO_model_Beddington_DeAngelis_activation_ratio_1_30.0

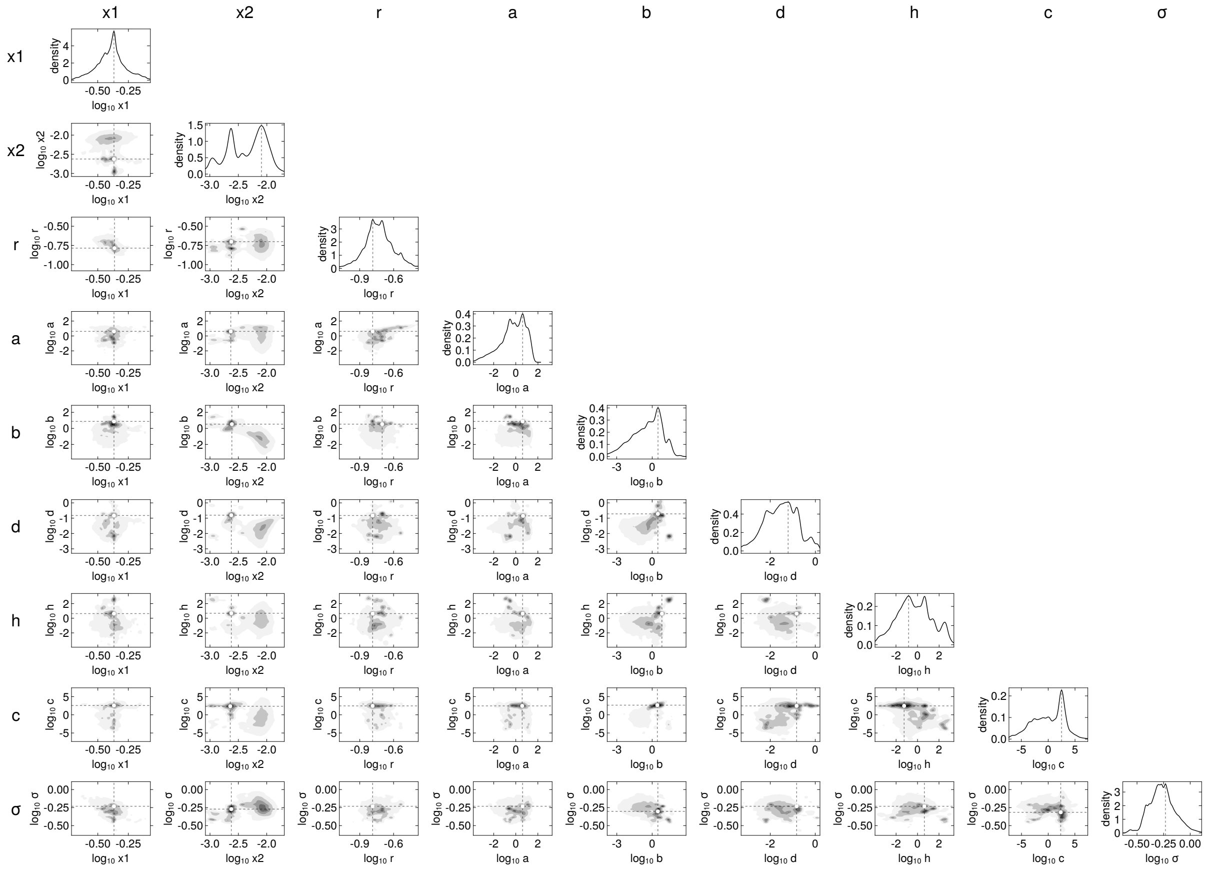


252

antigen_CD123w_effector_uT_target_WT_model_Beddington_DeAngelis_activation_ratio_1_0.5

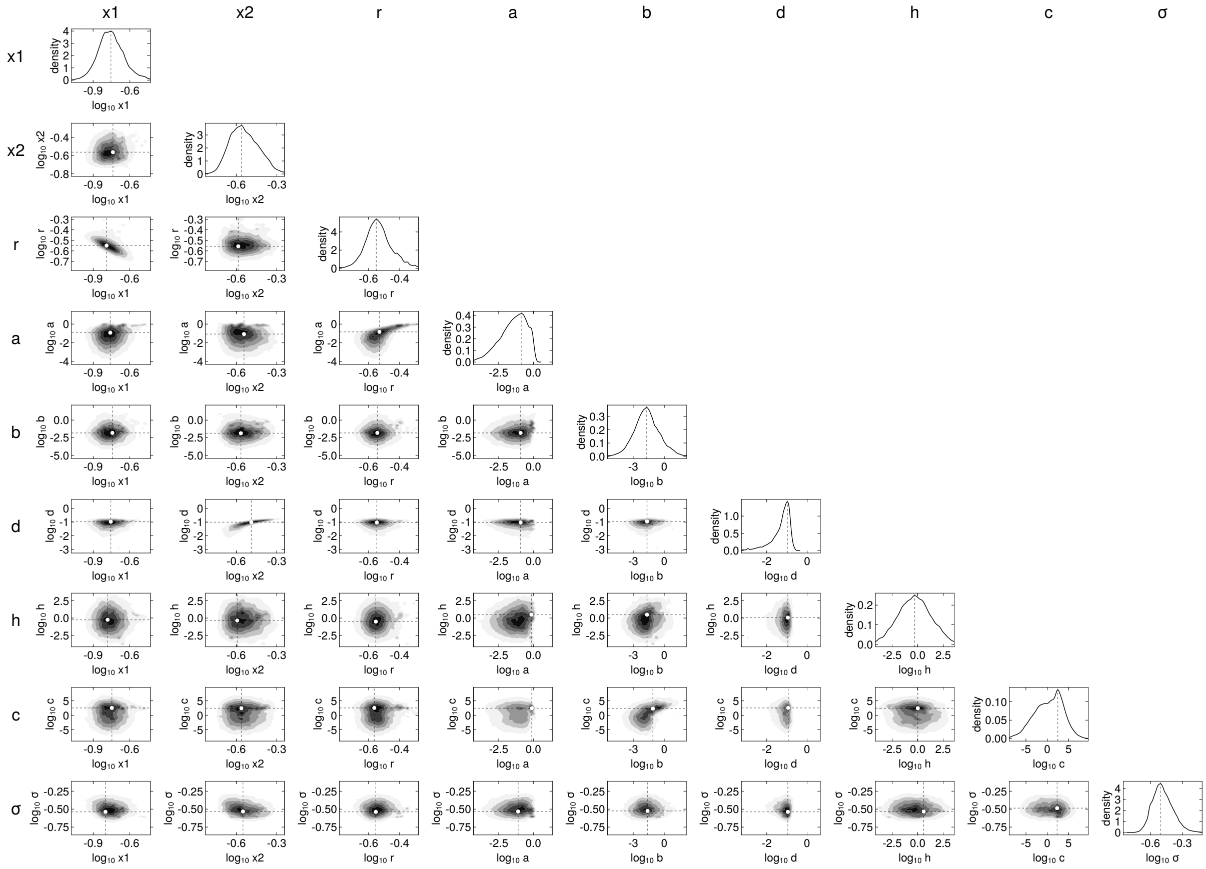

253

antigen_CD123w_effector_uT_target_WT_model_Beddington_DeAngelis_activation_ratio_1_1.0

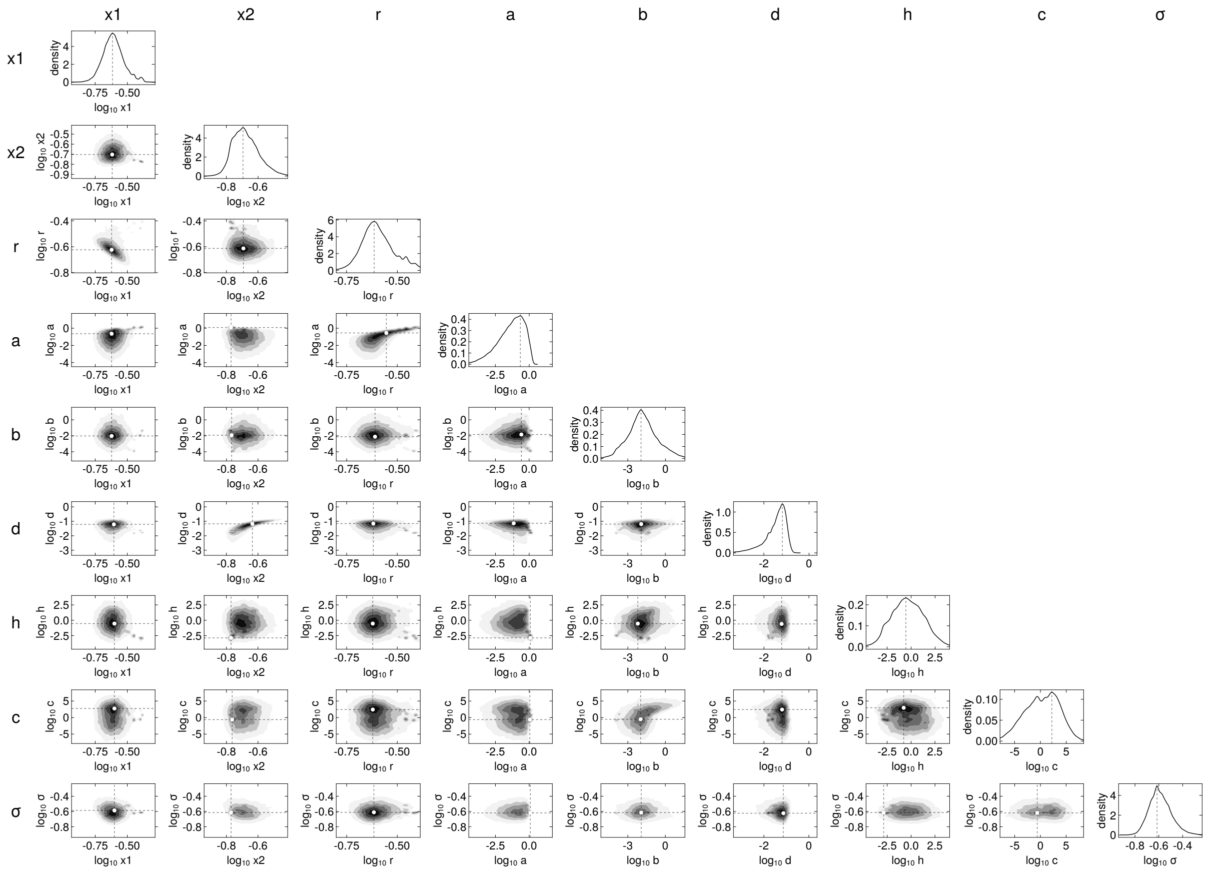


254

antigen_CD123w_effector_uT_target_WT_model_Beddington_DeAngelis_activation_ratio_1_2.0

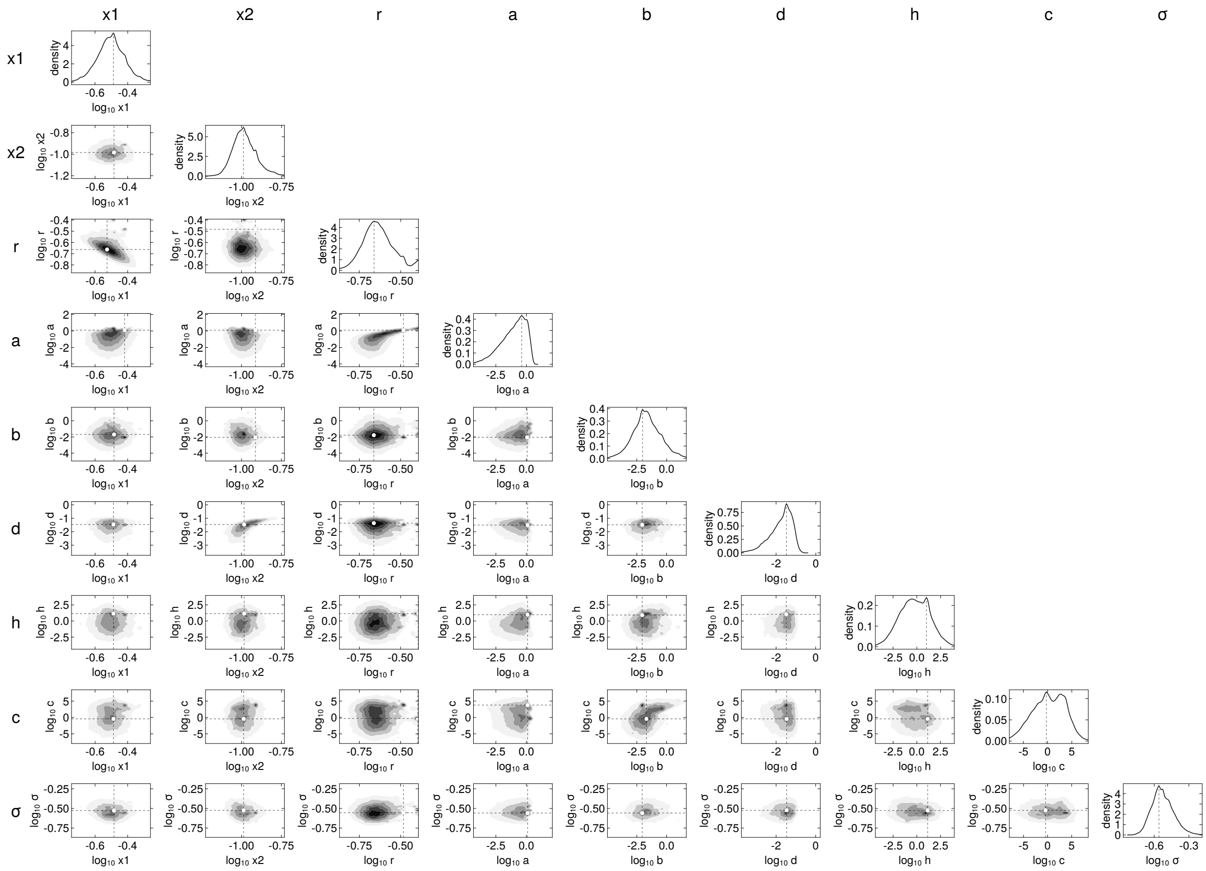


255

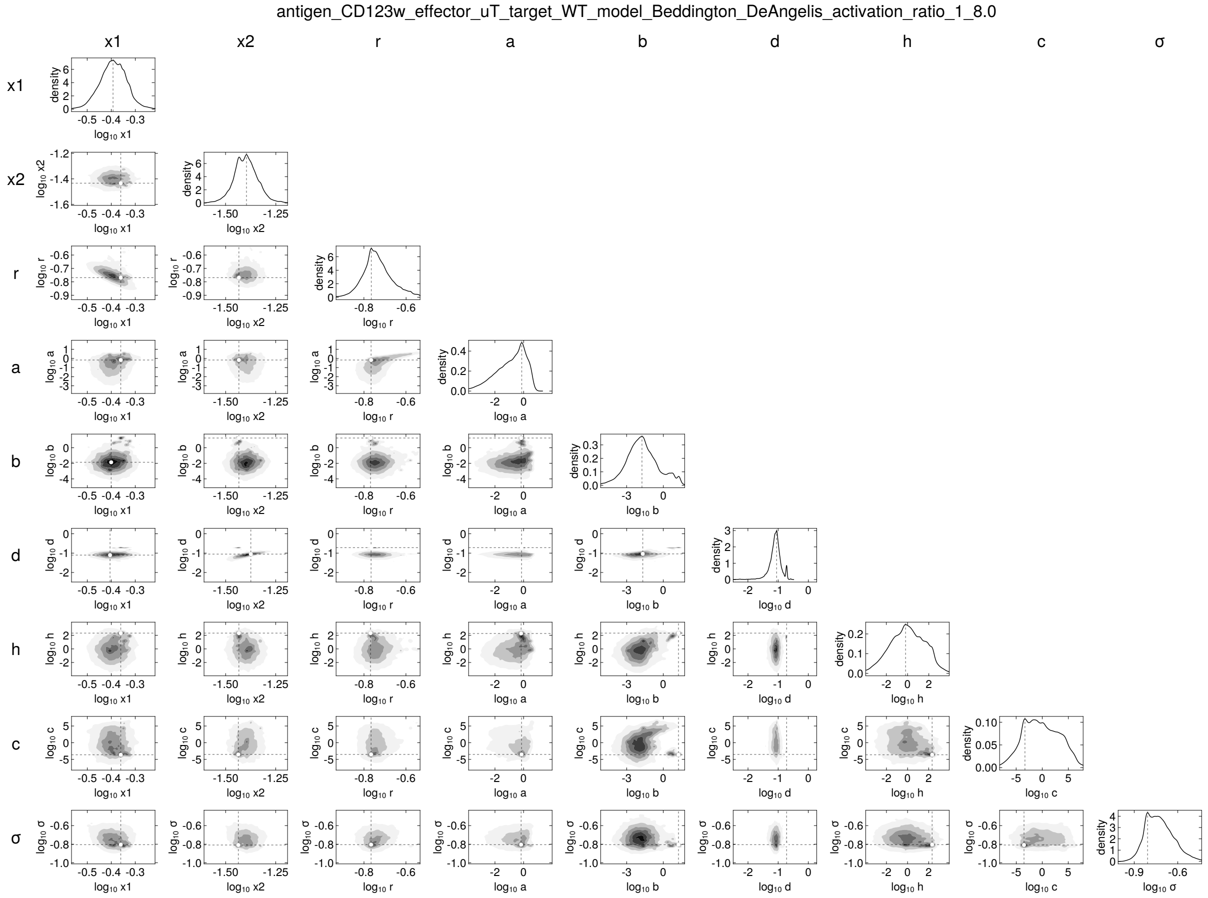
antigen_CD123w_effector_uT_target_WT_model_Beddington_DeAngelis_activation_ratio_1_8.0
x1
x2
r
a
b
d
h
c
σ
density
log10 x1
log10 x2
log10 r
log10 a
log10 b
log10 d
log10 h
log10 c
log10 σ

256

antigen_CD123w_effector_uT_target_KO_model_Beddington_DeAngelis_activation_ratio_1_0.5

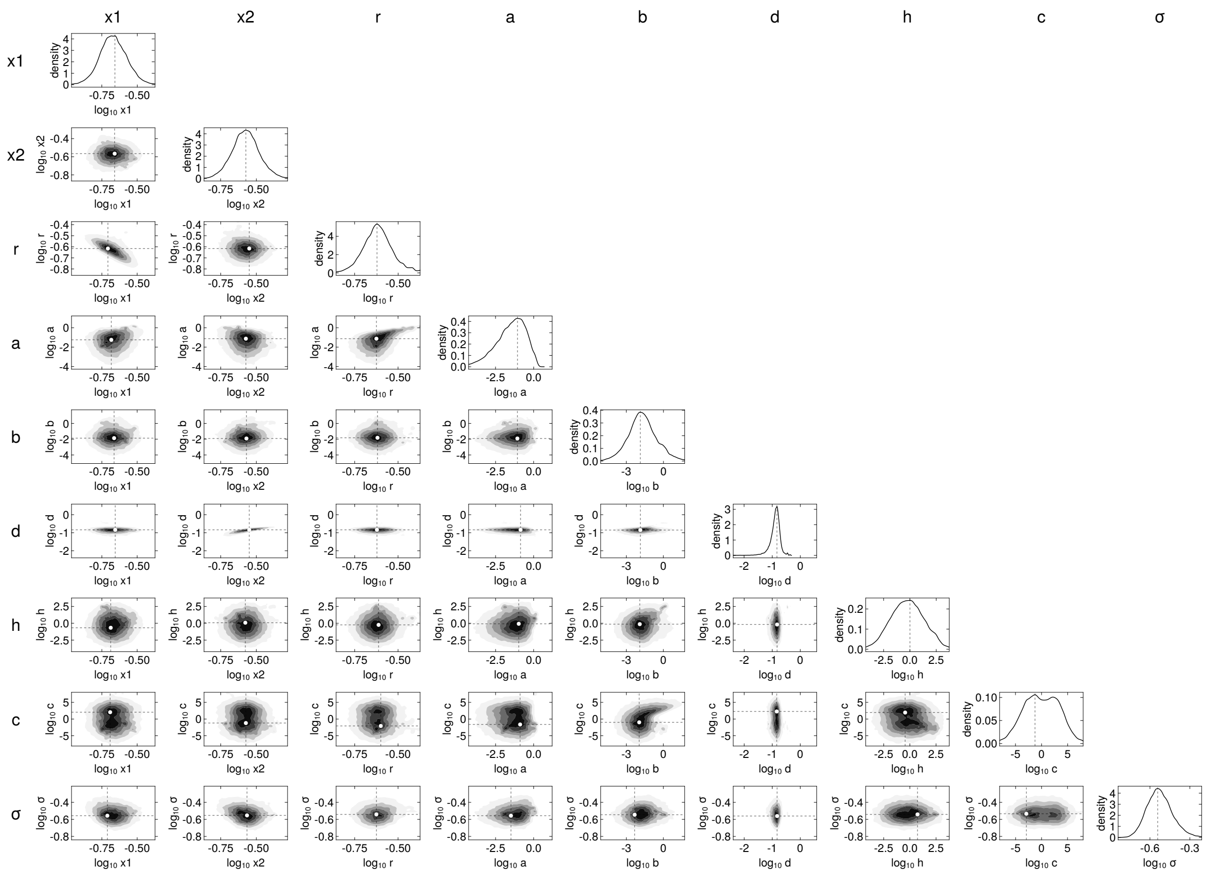

257

antigen_CD123w_effector_uT_target_KO_model_Beddington_DeAngelis_activation_ratio_1_1.0

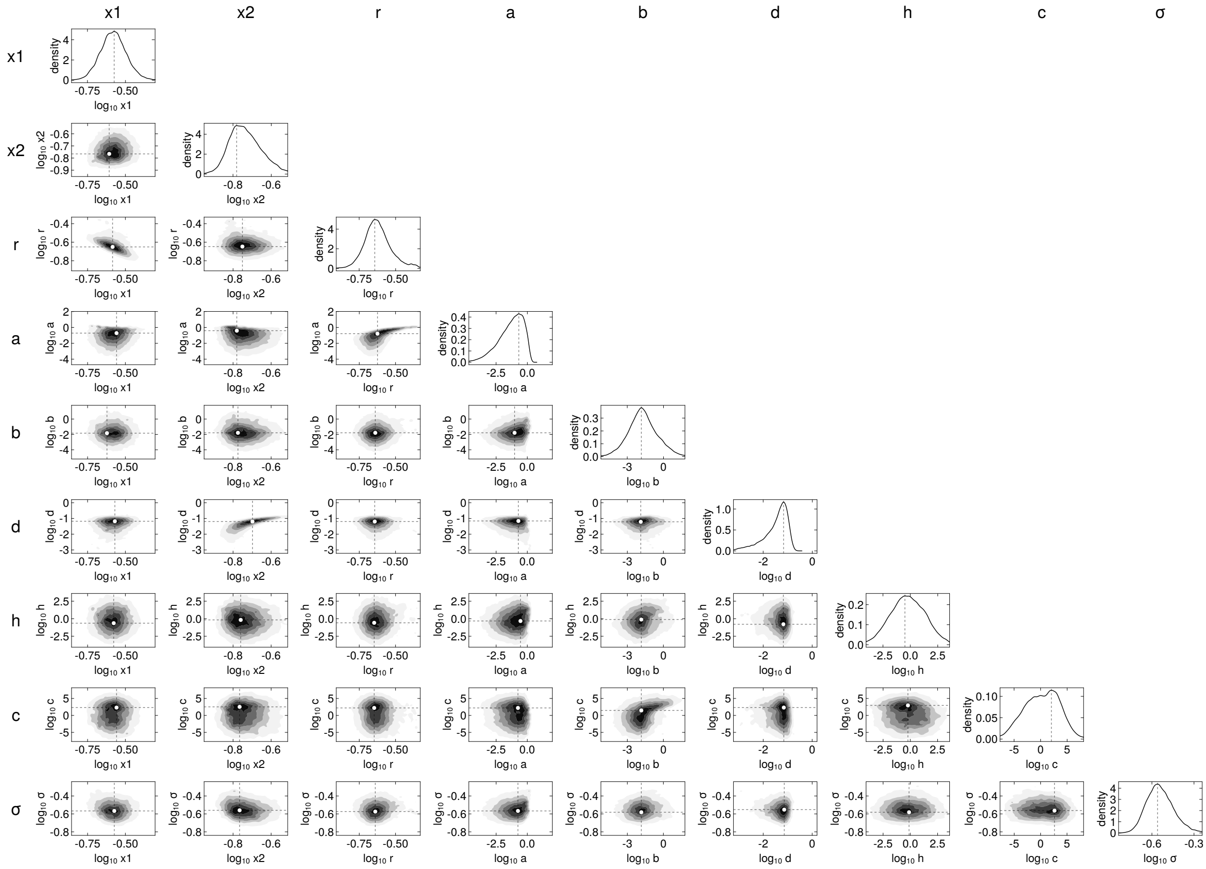


258

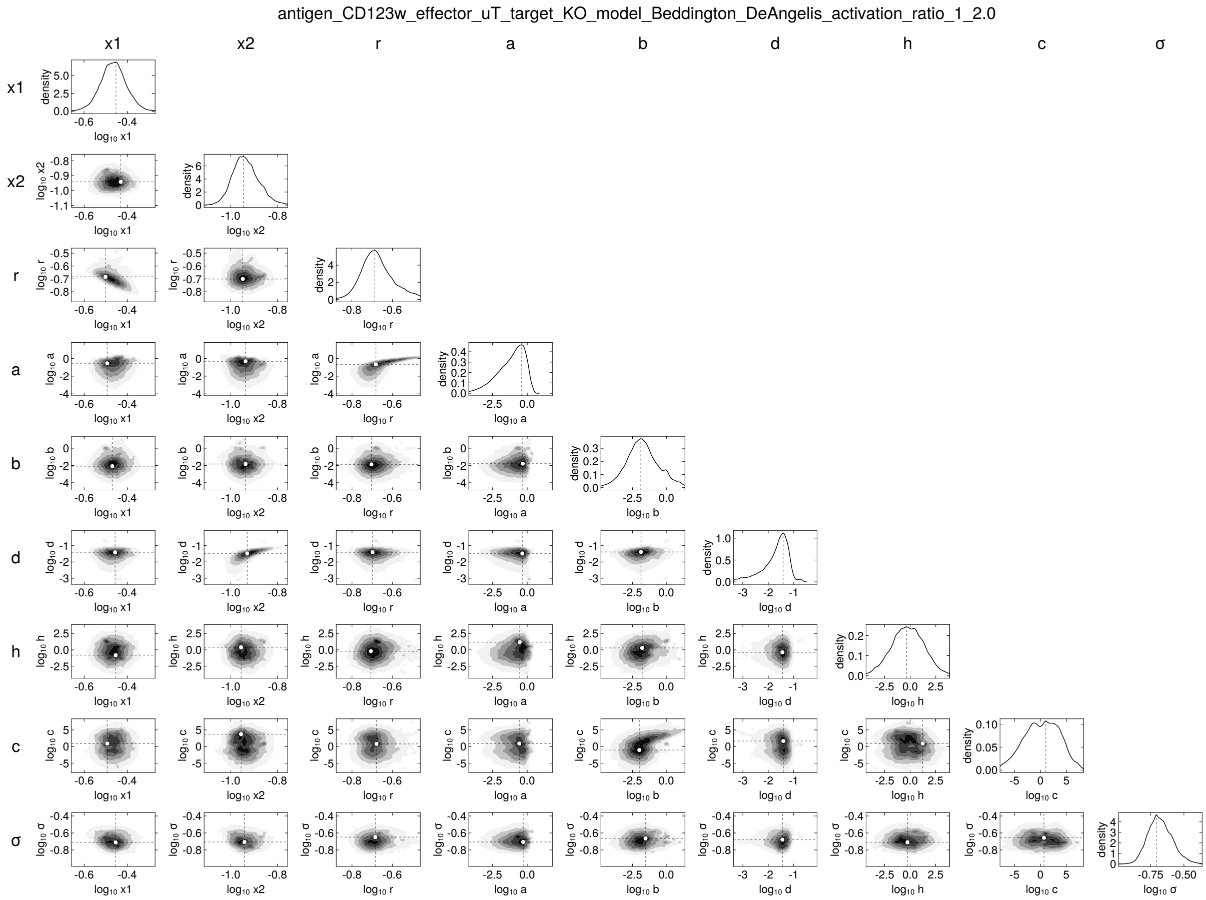
antigen_CD123w_effector_uT_target_KO_model_Beddington_DeAngelis_activation_ratio_1_2.0
x1
x2
r
a
b
d
h
c
σ

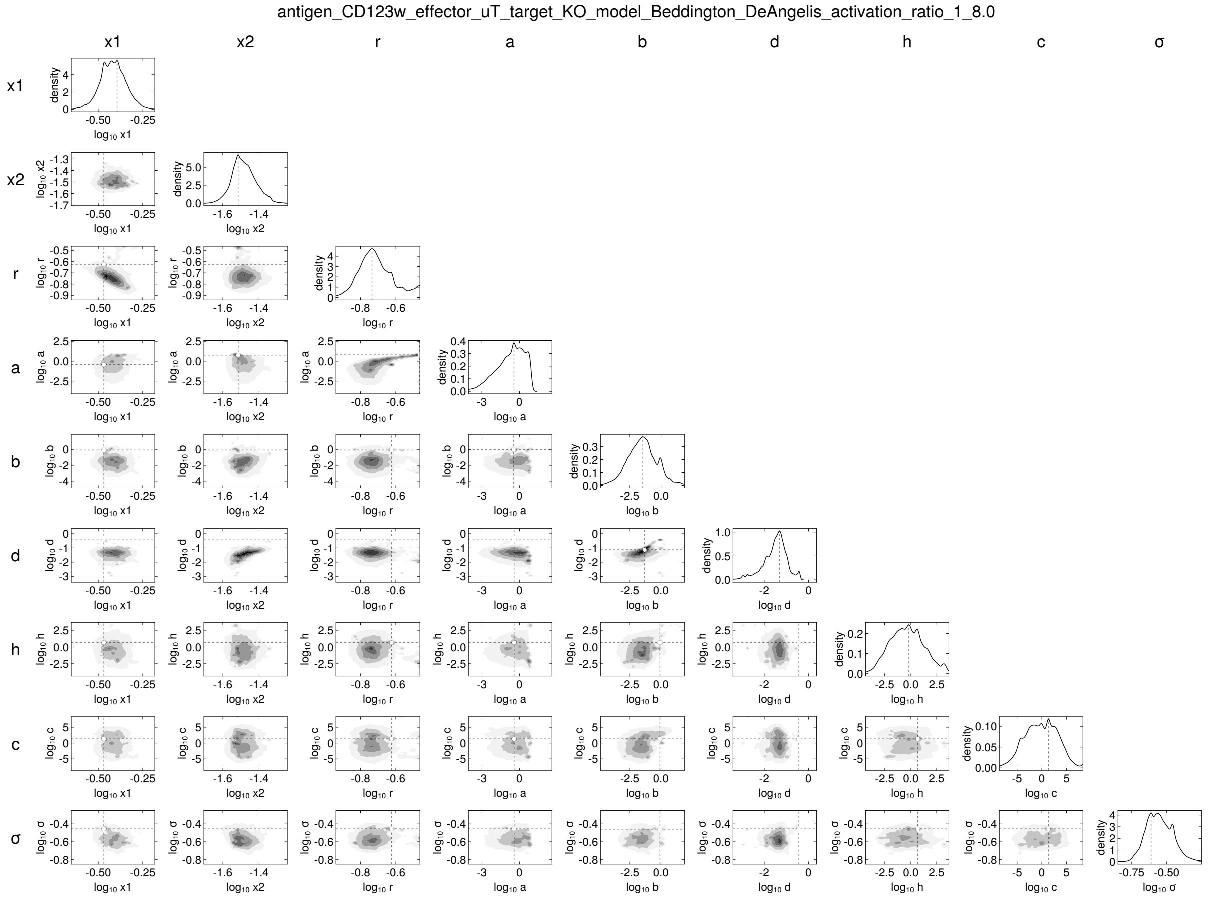
antigen_CD123w_effector_uT_target_KO_model_Beddington_DeAngelis_activation_ratio_1_8.0
x1
x2
r
a
b
d
h
c
σ
density
log10 x1
log10 x2
log10 r
log10 a
log10 b
log10 d
log10 h
log10 c
log10 σ

260

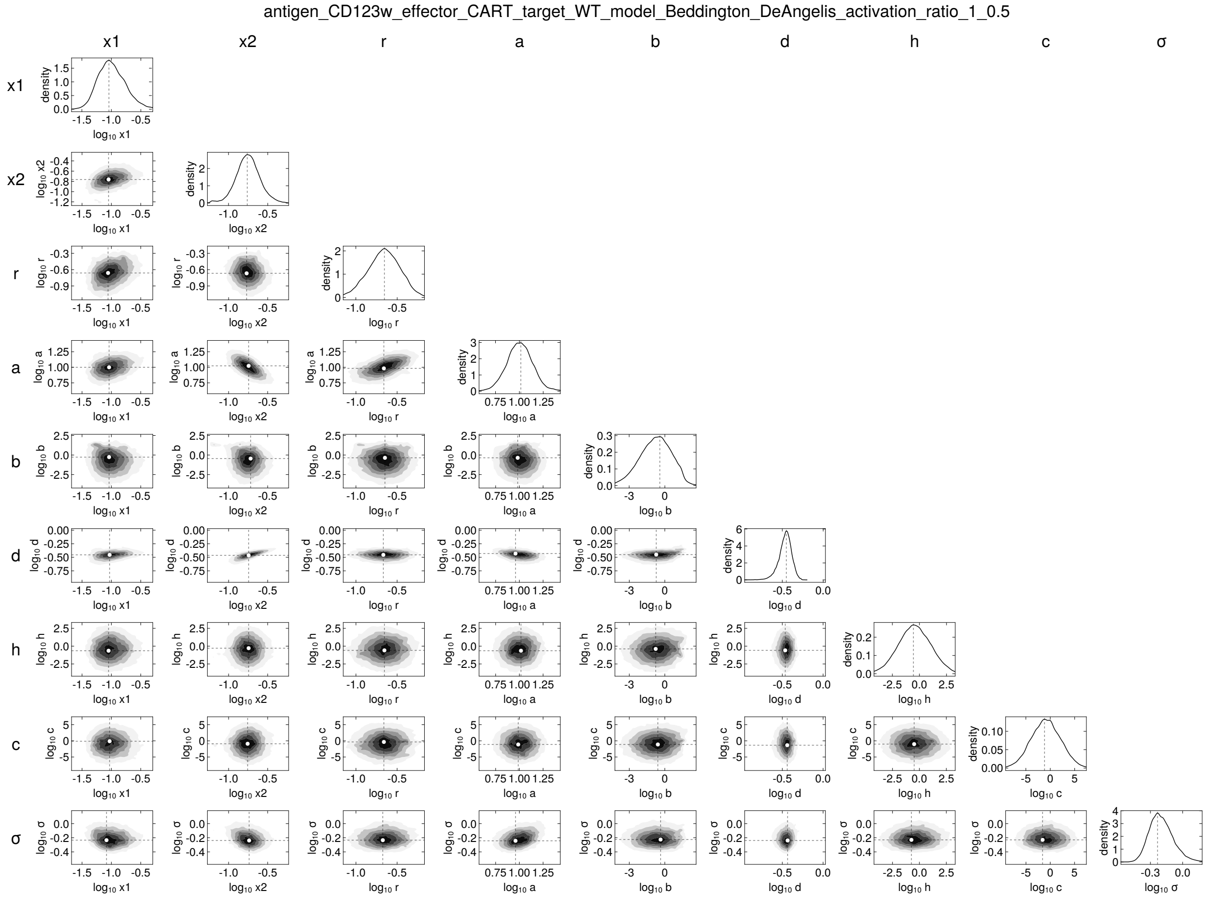
antigen_CD123w_effector_CART_target_WT_model_Beddington_DeAngelis_activation_ratio_1_0.5
x1
x2
r
a
b
d
h
c
σ

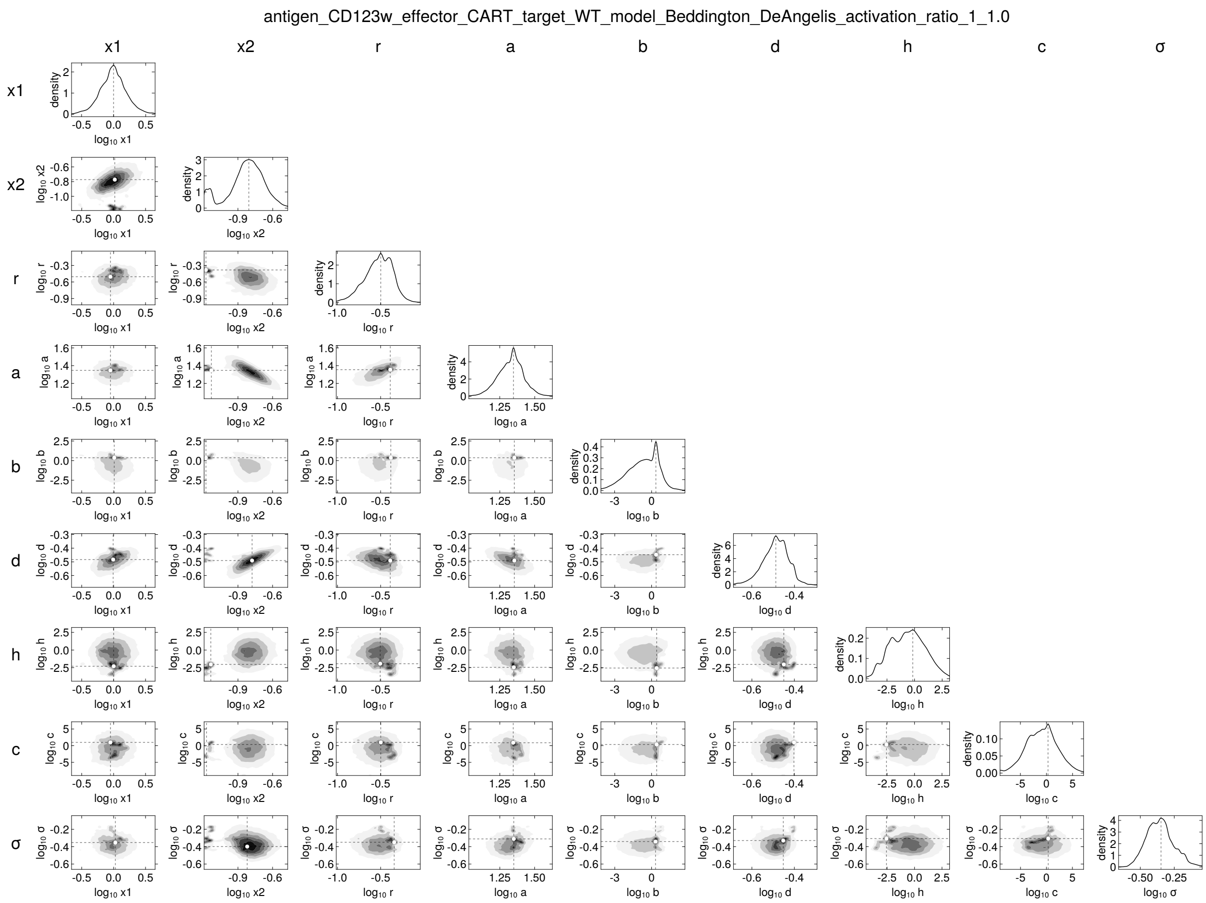

antigen_CD123w_effector_CART_target_WT_model_Beddington_DeAngelis_activation_ratio_1_1.0
x1
x2
r
a
b
d
h
c
σ
density
log10 x1
log10 x2
log10 r
log10 a
log10 b
log10 d
log10 h
log10 c
log10 σ

antigen_CD123w_effector_CART_target_WT_model_Beddington_DeAngelis_activation_ratio_1_2.0

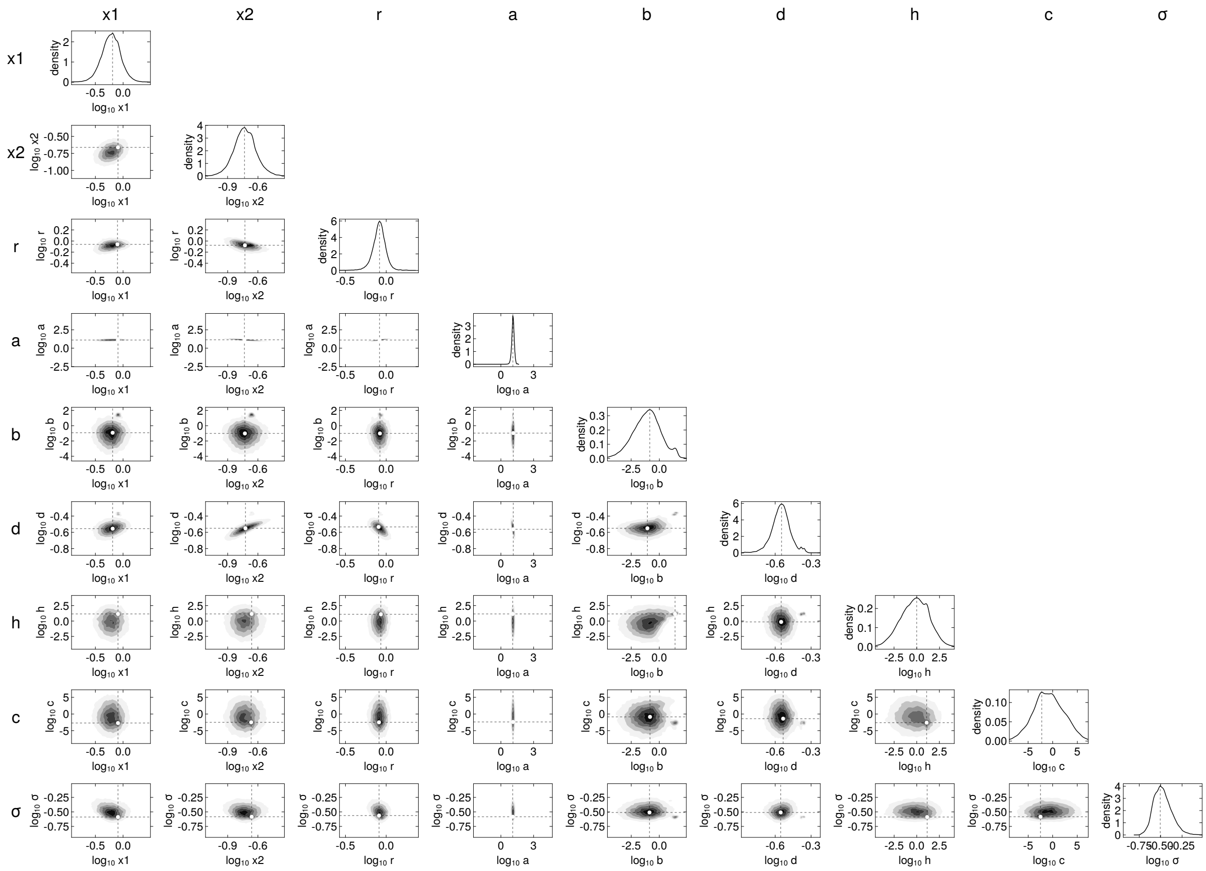


263

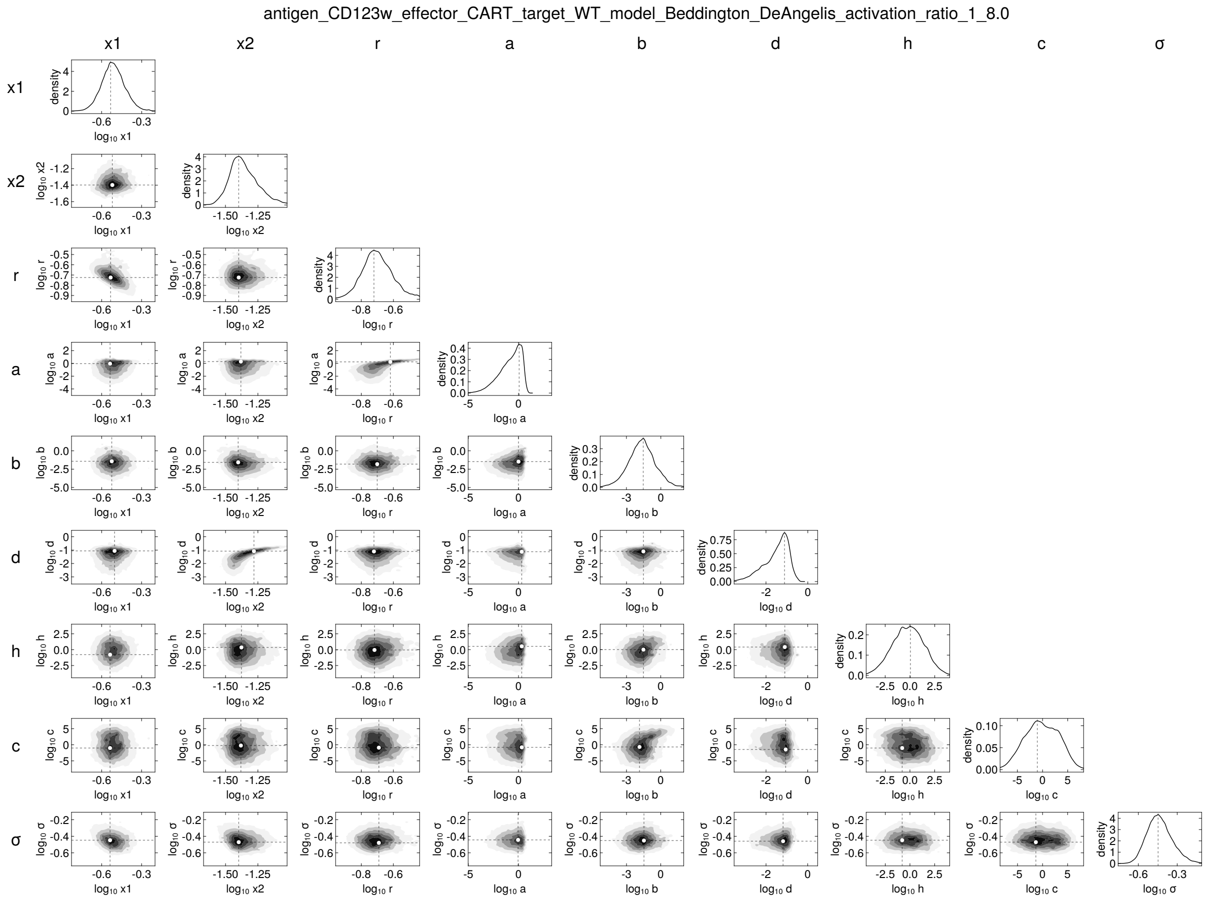
antigen_CD123w_effector_CART_target_WT_model_Beddington_DeAngelis_activation_ratio_1_8.0
x1
x2
r
a
b
d
h
c
σ

antigen_CD123w_effector_CART_target_KO_model_Beddington_DeAngelis_activation_ratio_1_0.5

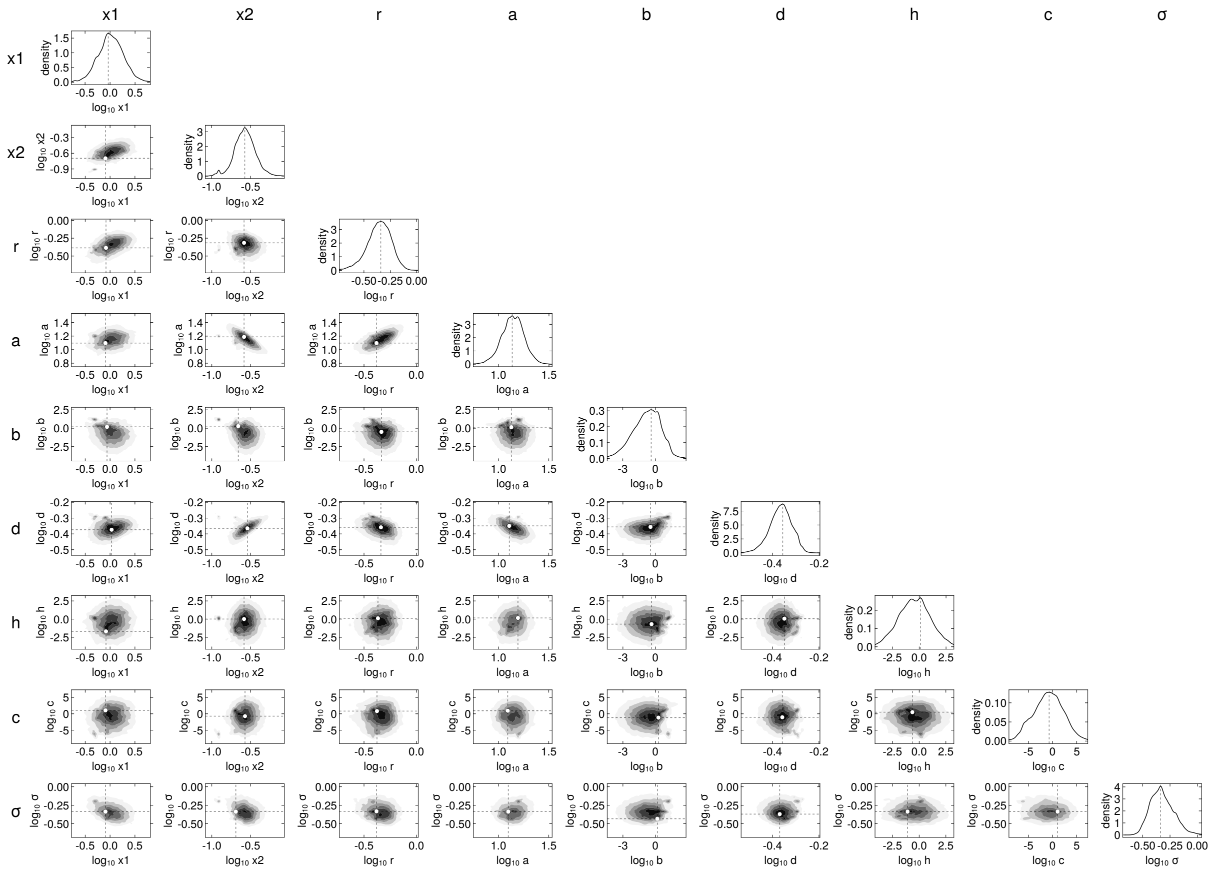

antigen_CD123w_effector_CART_target_KO_model_Beddington_DeAngelis_activation_ratio_1_1.0

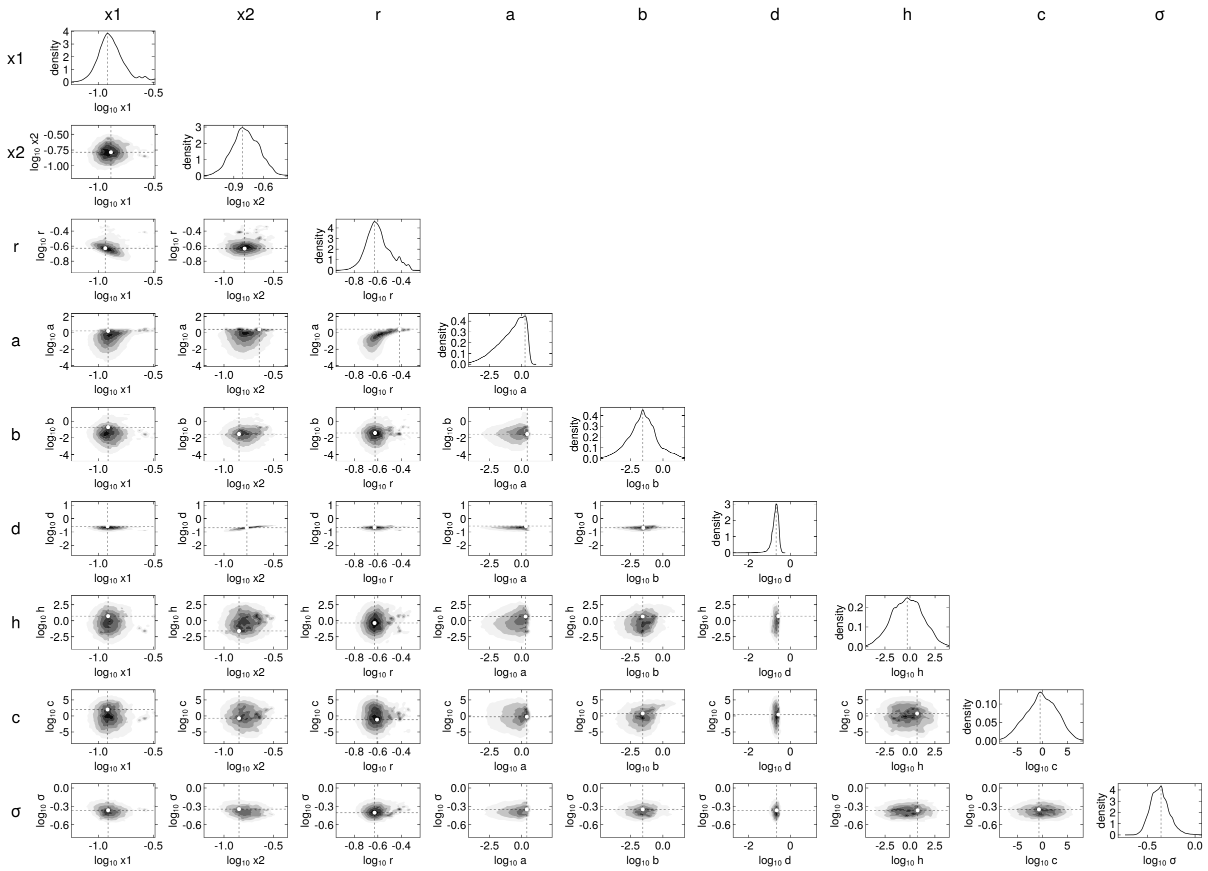


266

antigen_CD123w_effector_CART_target_KO_model_Beddington_DeAngelis_activation_ratio_1_2.0

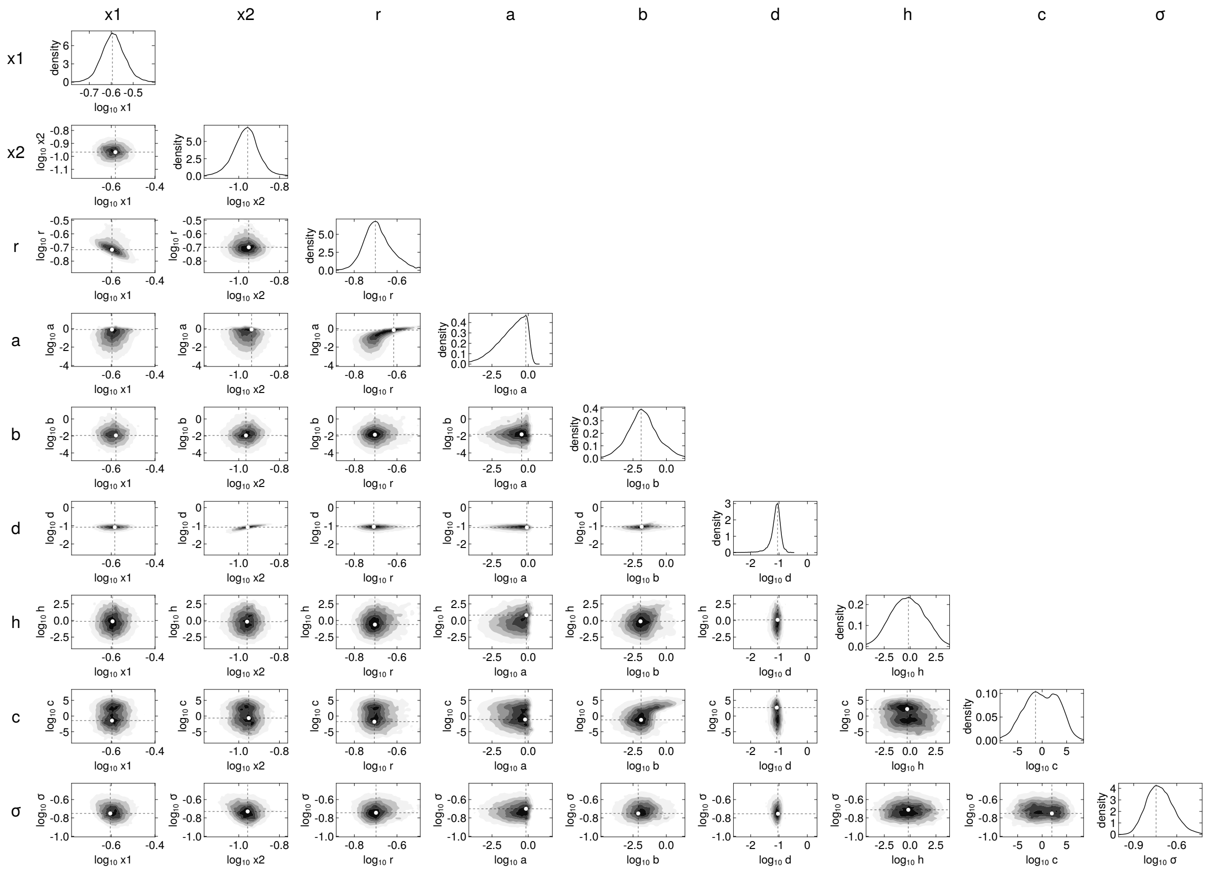


267

antigen_CD123w_effector_CART_target_KO_model_Beddington_DeAngelis_activation_ratio_1_8.0

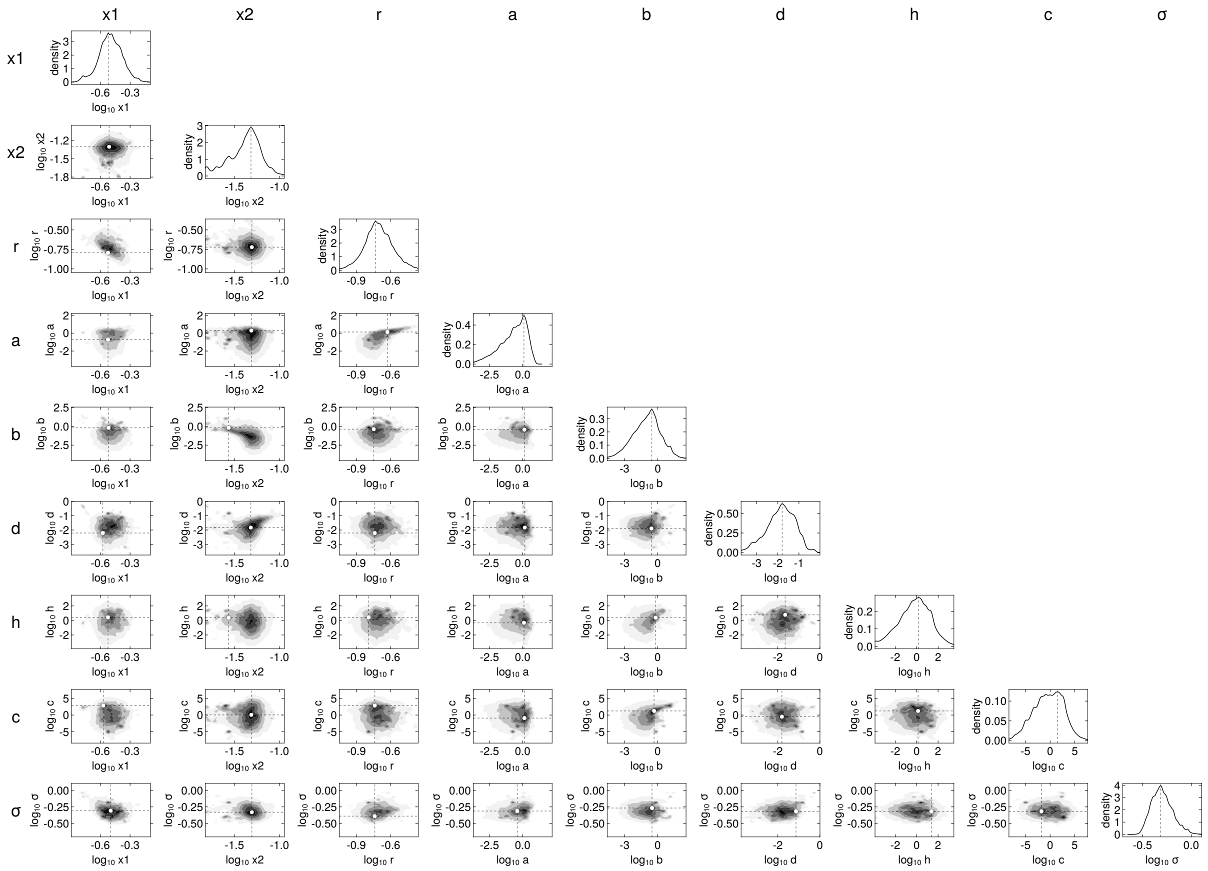

268

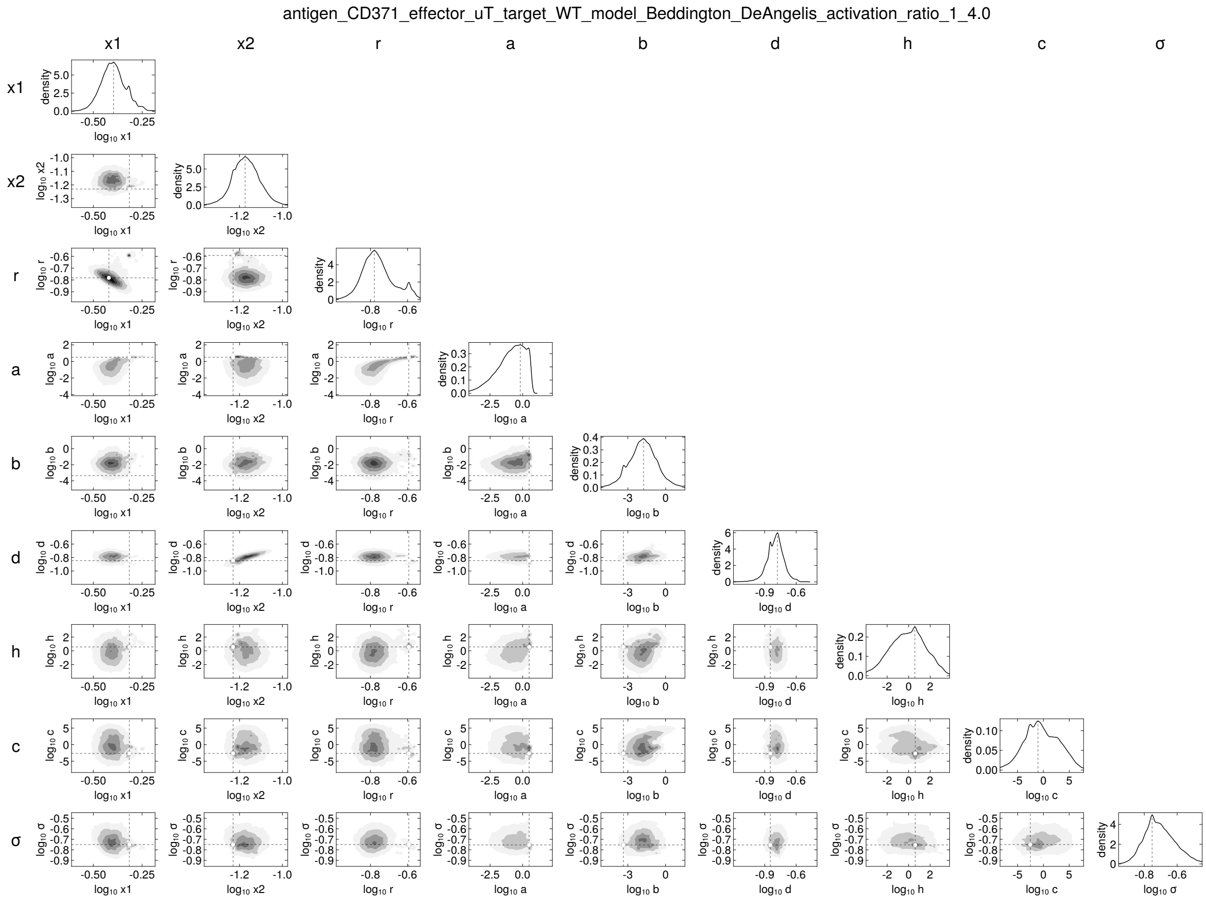
antigen_CD371_effector_uT_target_WT_model_Beddington_DeAngelis_activation_ratio_1_4.0
x1
x2
r
a
b
d
h
c
σ
density
log10 x1
log10 x2
log10 r
log10 a
log10 b
log10 d
log10 h
log10 c
log10 σ

antigen_CD371_effector_uT_target_WT_model_Beddington_DeAngelis_activation_ratio_1_16.0

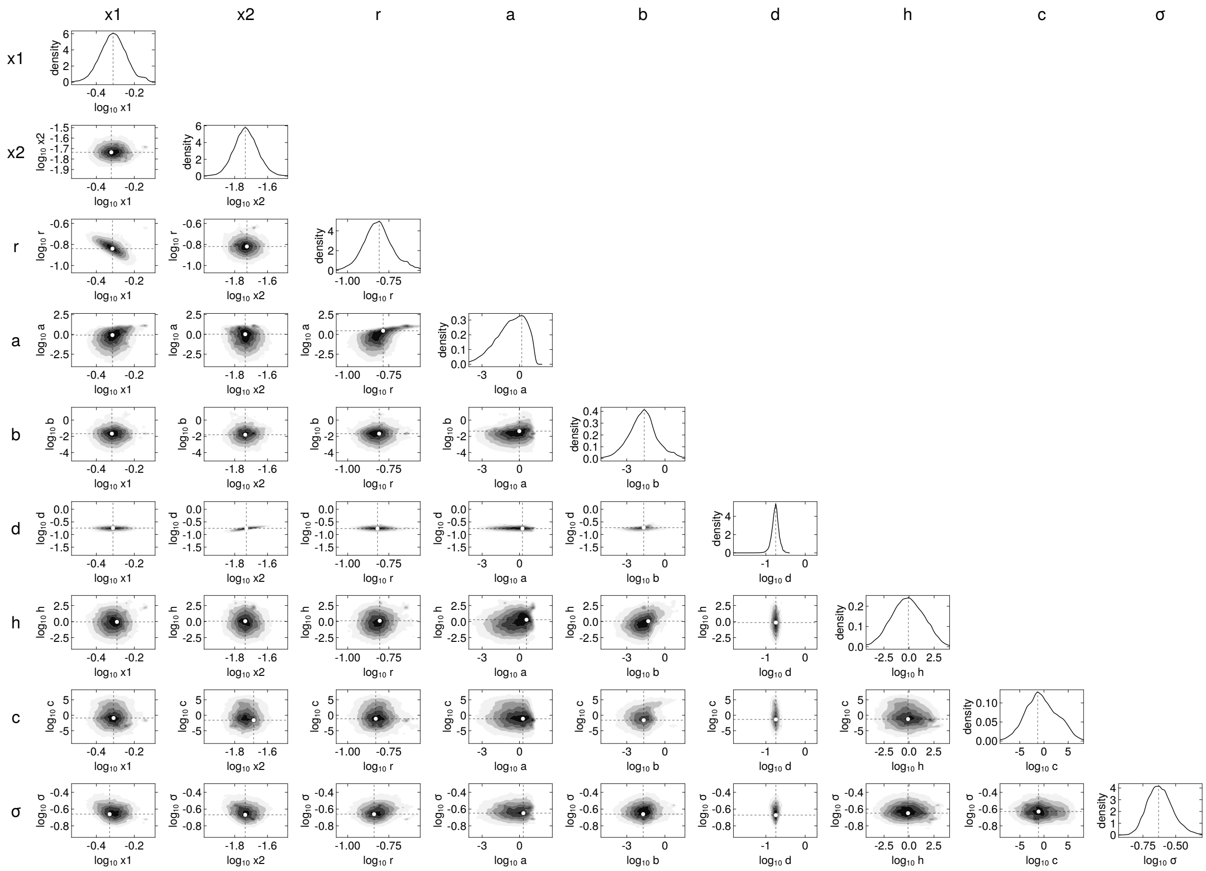


270

antigen_CD371_effector_uT_target_WT_model_Beddington_DeAngelis_activation_ratio_1_40.0

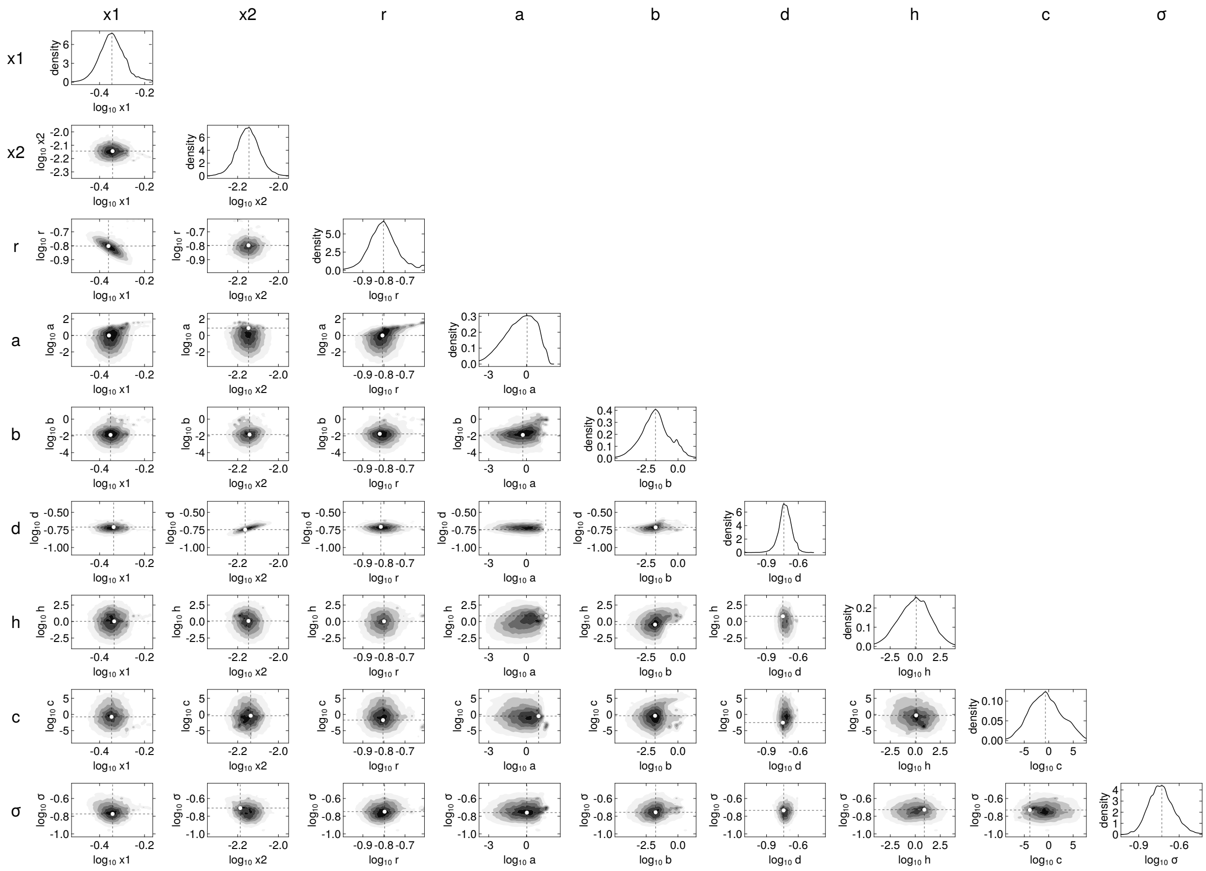


271

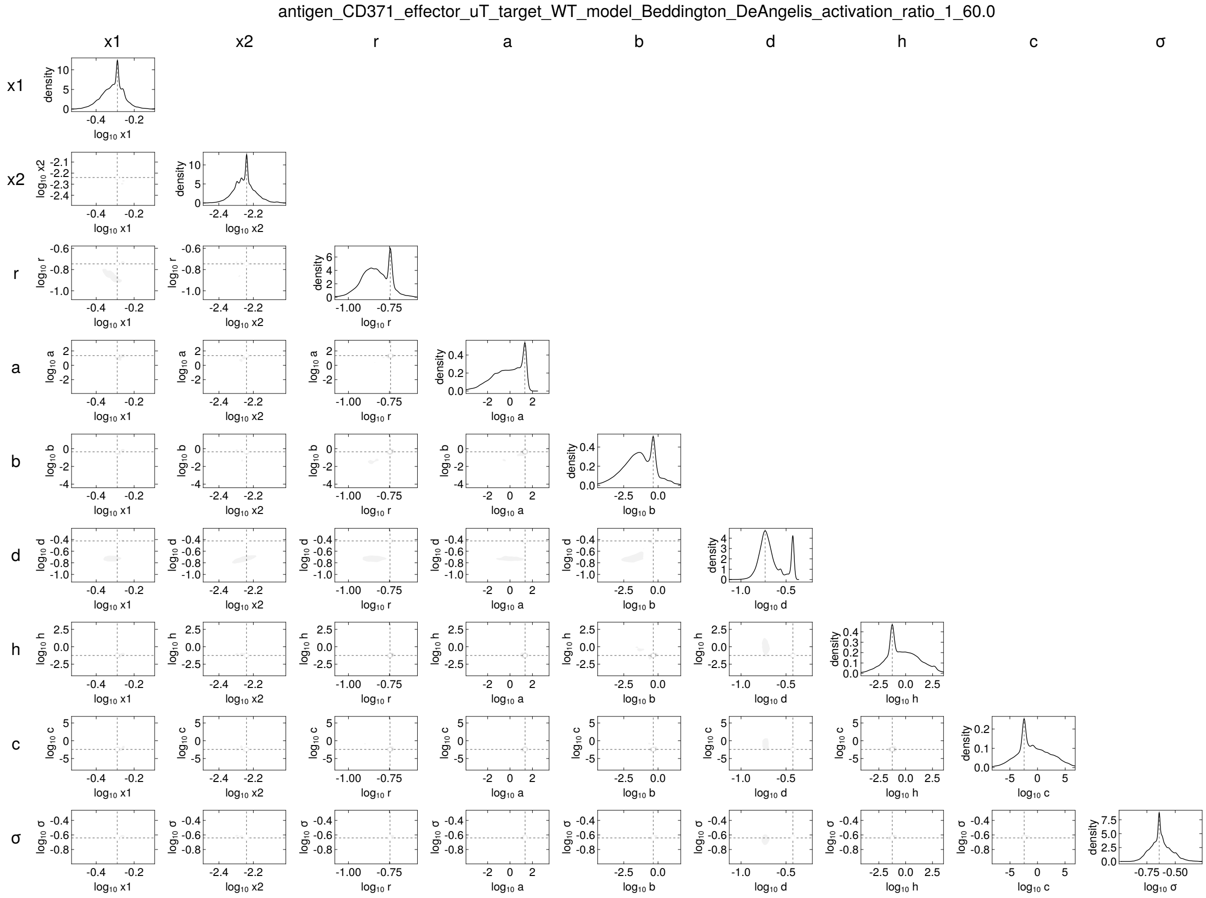
antigen_CD371_effector_uT_target_WT_model_Beddington_DeAngelis_activation_ratio_1_60.0
x1
x2
r
a
b
d
h
c
σ
density
log10 x1
log10 x2
log10 r
log10 a
log10 b
log10 d
log10 h
log10 c
log10 σ

272

antigen_CD371_effector_uT_target_KO_model_Beddington_DeAngelis_activation_ratio_1_4.0

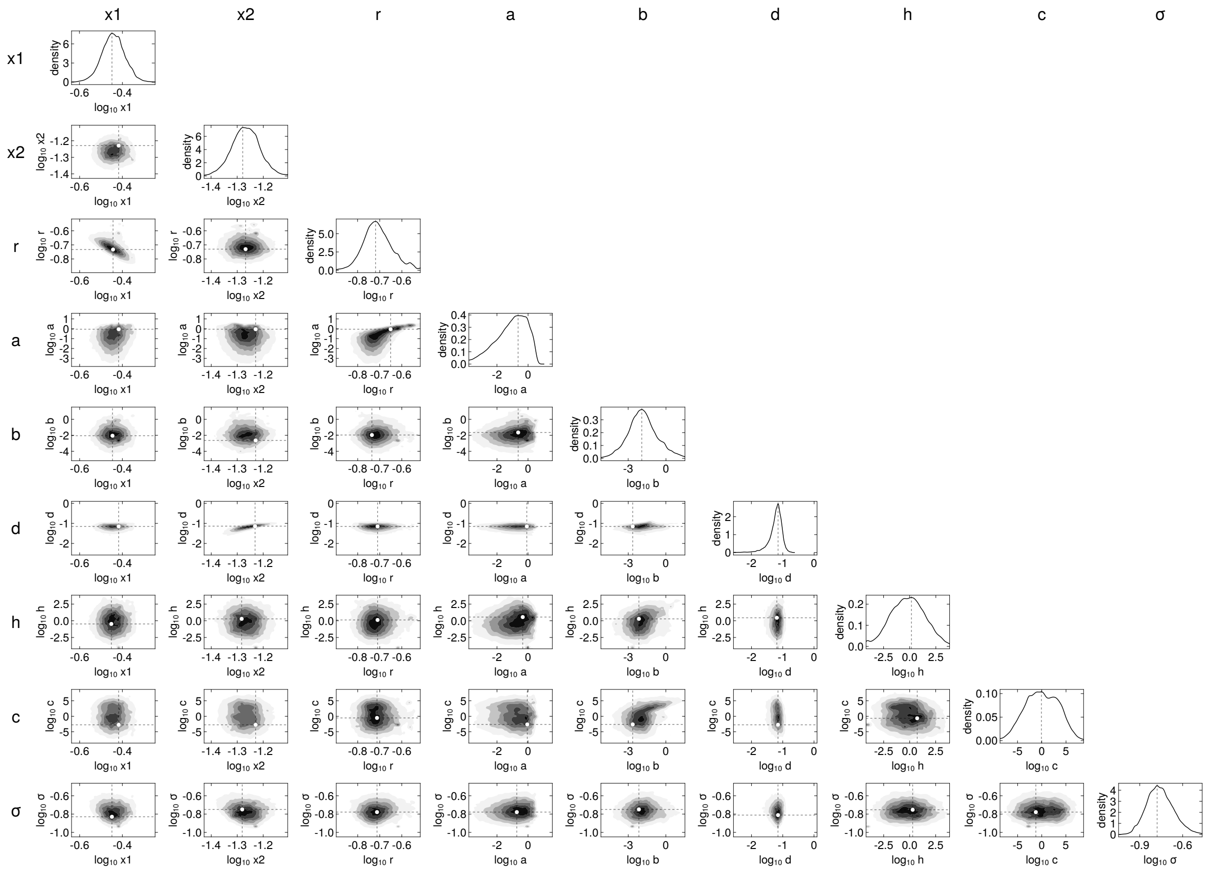


273

antigen_CD371_effector_uT_target_KO_model_Beddington_DeAngelis_activation_ratio_1_16.0

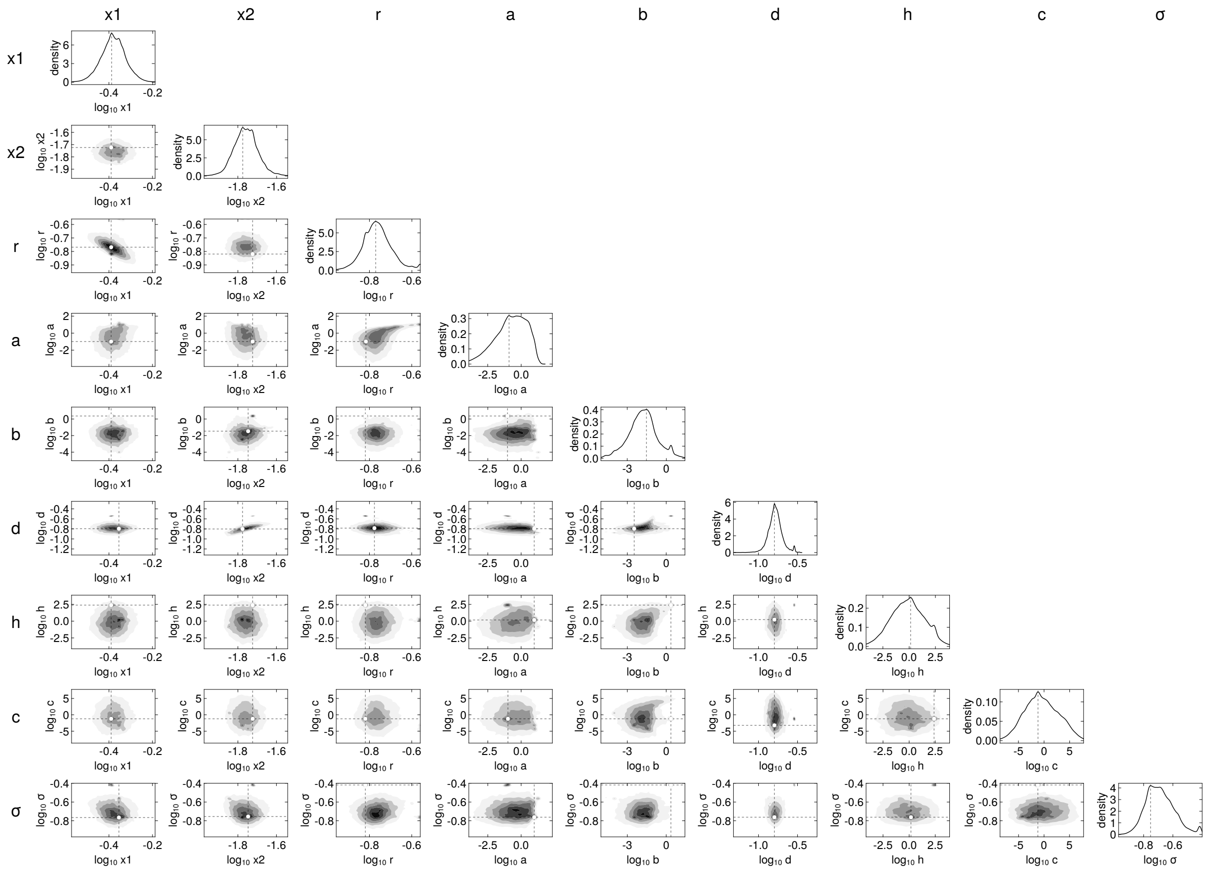


274

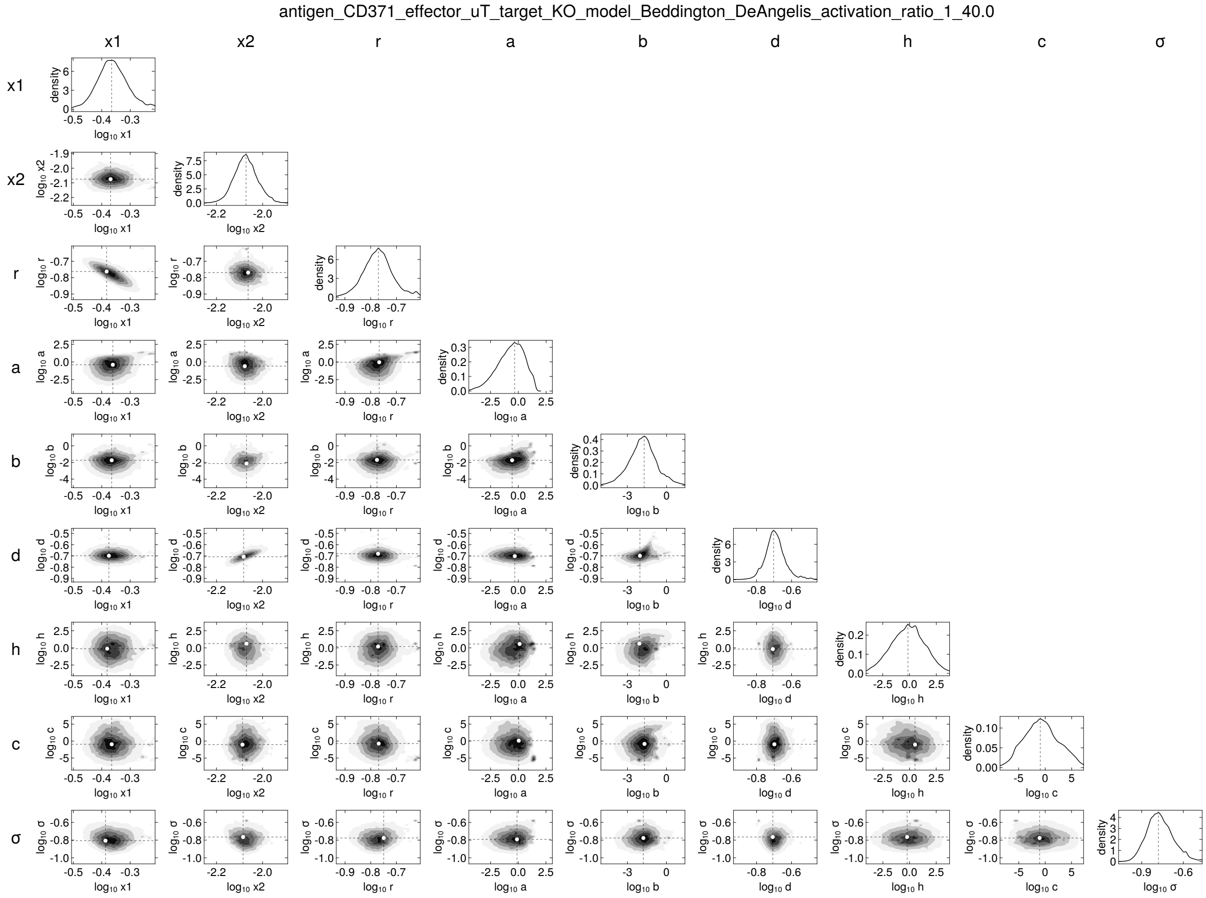
antigen_CD371_effector_uT_target_KO_model_Beddington_DeAngelis_activation_ratio_1_40.0
x1
x2
r
a
b
d
h
c
σ
density
log10 x1
log10 x2
log10 r
log10 a
log10 b
log10 d
log10 h
log10 c
log10 σ

275

antigen_CD371_effector_uT_target_KO_model_Beddington_DeAngelis_activation_ratio_1_60.0

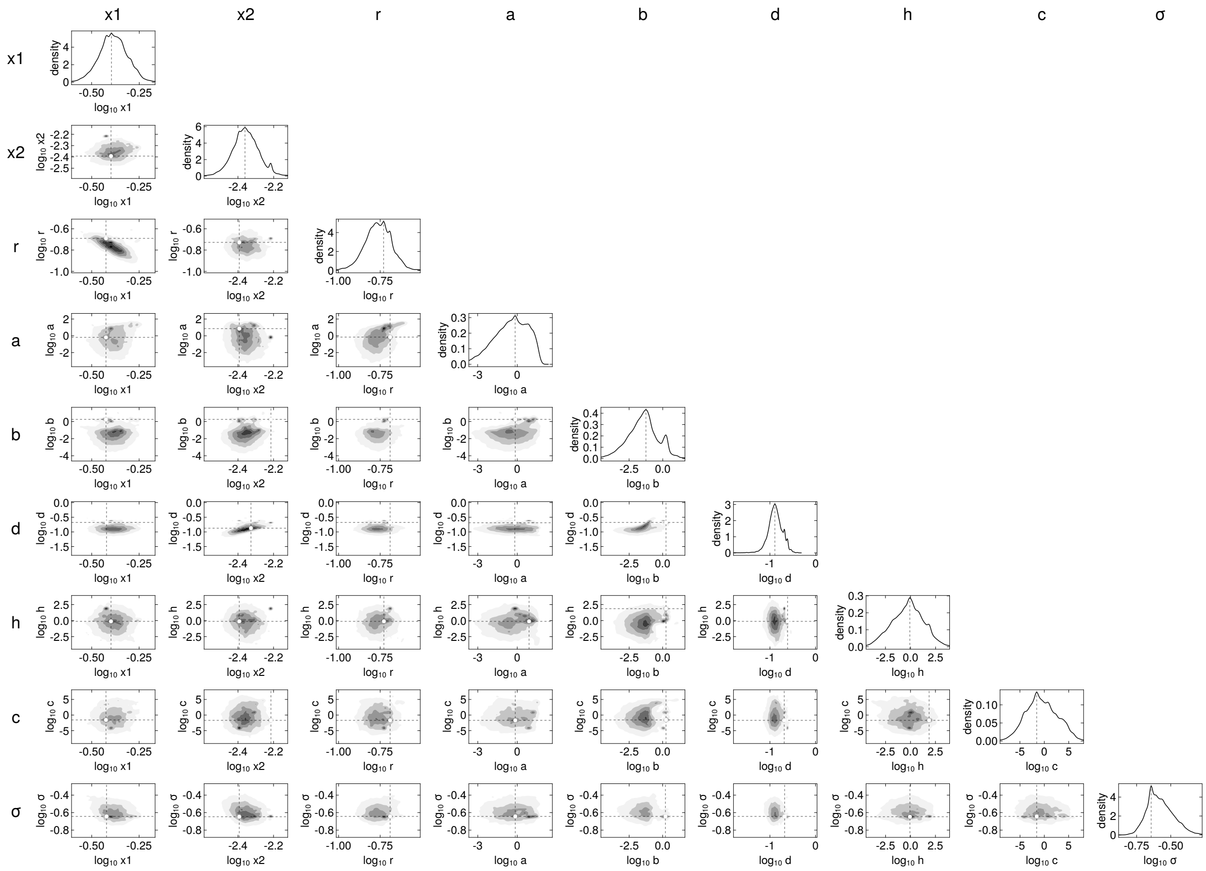

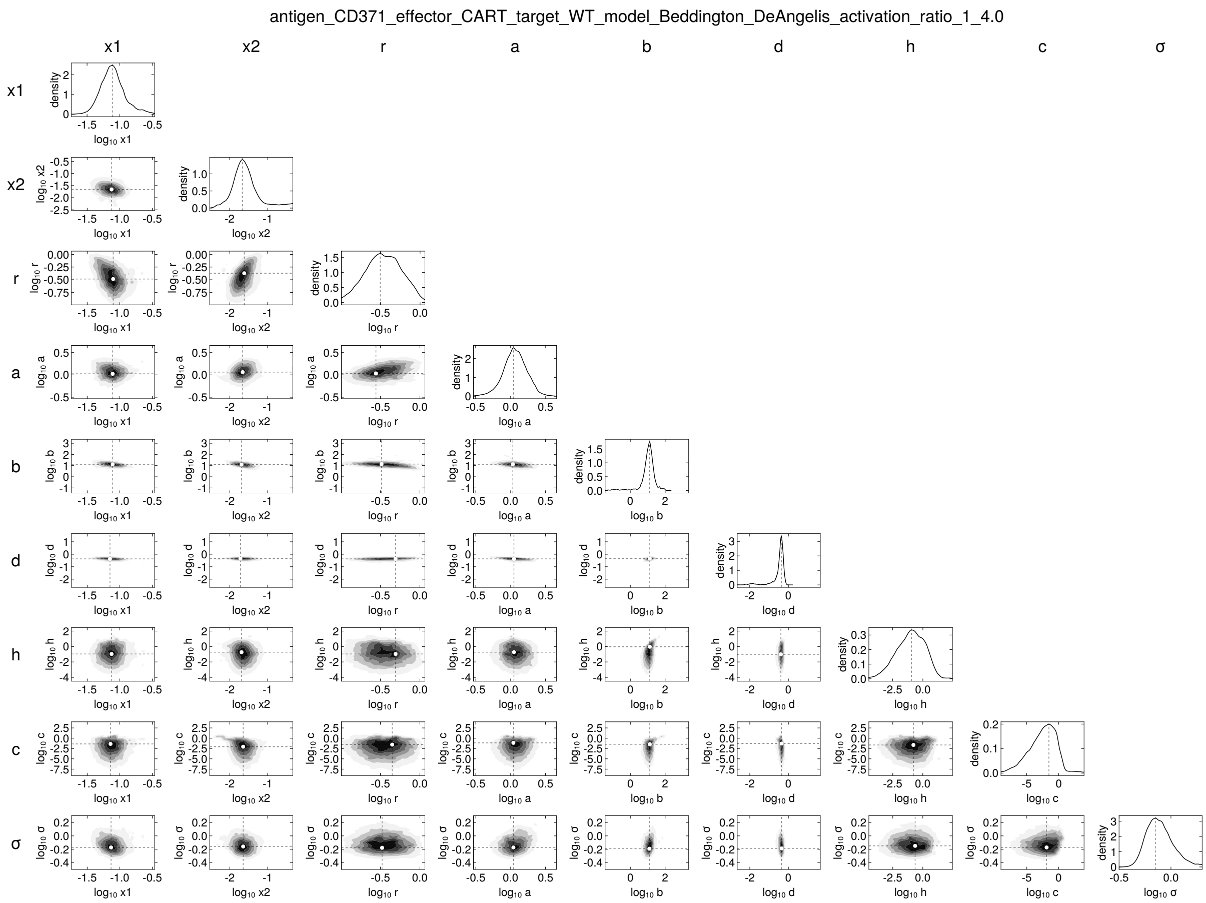
antigen_CD371_effector_CART_target_WT_model_Beddington_DeAngelis_activation_ratio_1_4.0
x1
x2
r
a
b
d
h
c
σ
density
log10 x1
log10 x2
log10 r
log10 a
log10 b
log10 d
log10 h
log10 c
log10 σ

antigen_CD371_effector_CART_target_WT_model_Beddington_DeAngelis_activation_ratio_1_16.0

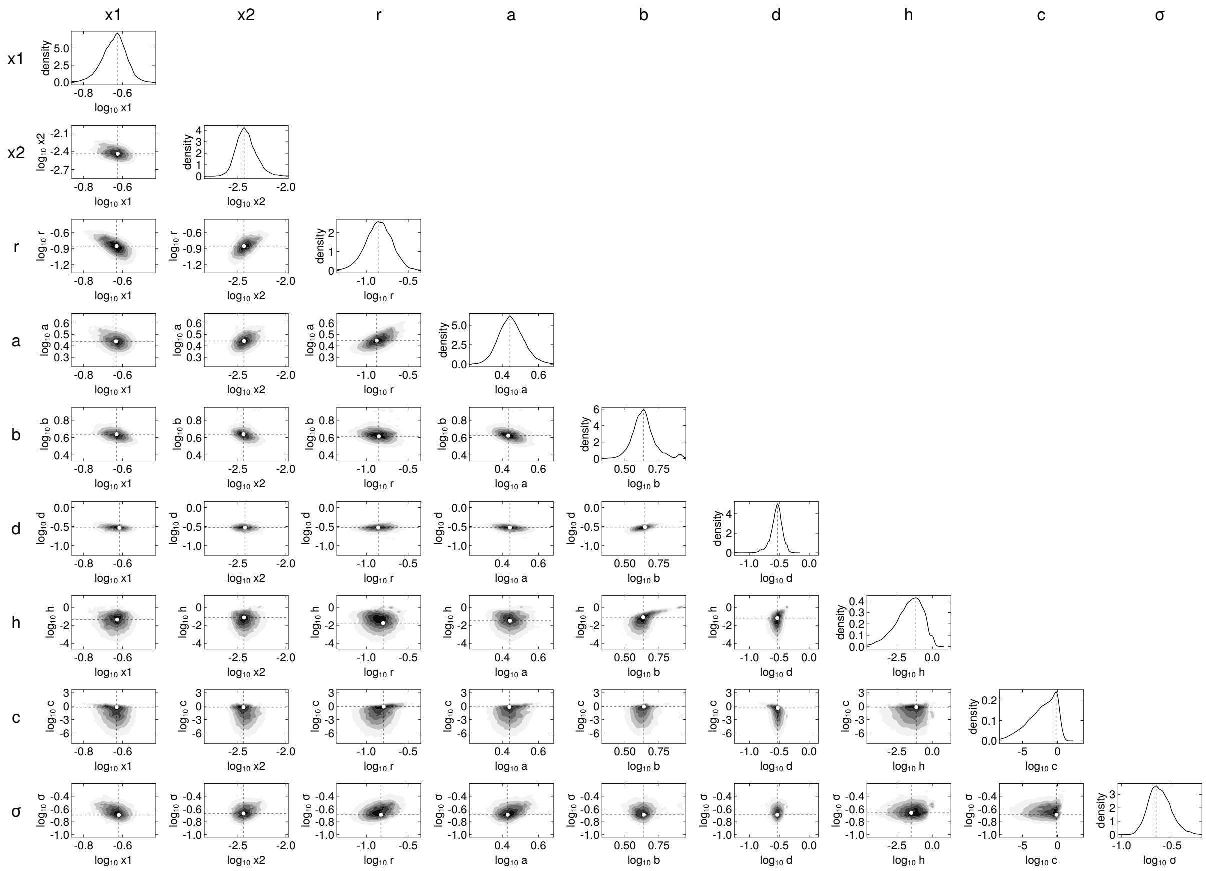


278

antigen_CD371_effector_CART_target_WT_model_Beddington_DeAngelis_activation_ratio_1_40.0

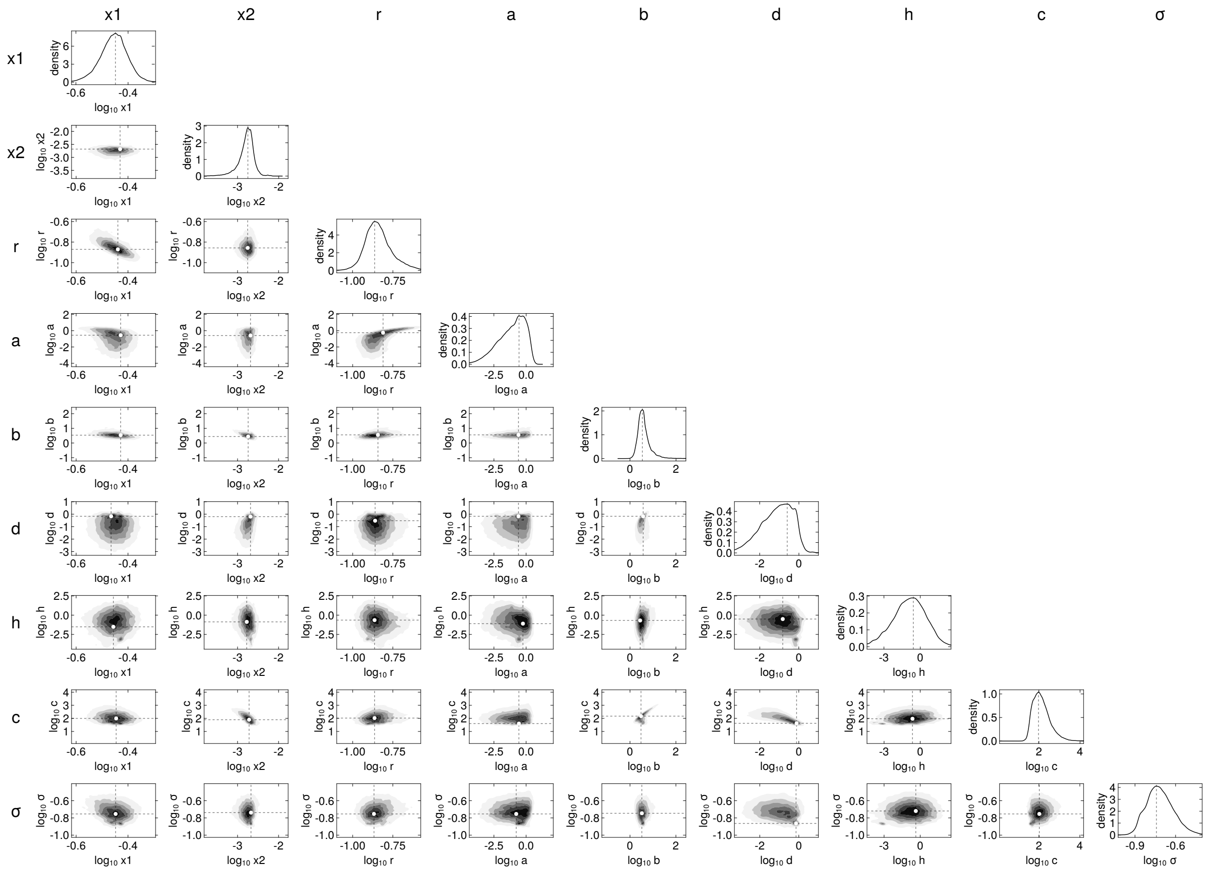


279

antigen_CD371_effector_CART_target_WT_model_Beddington_DeAngelis_activation_ratio_1_60.0

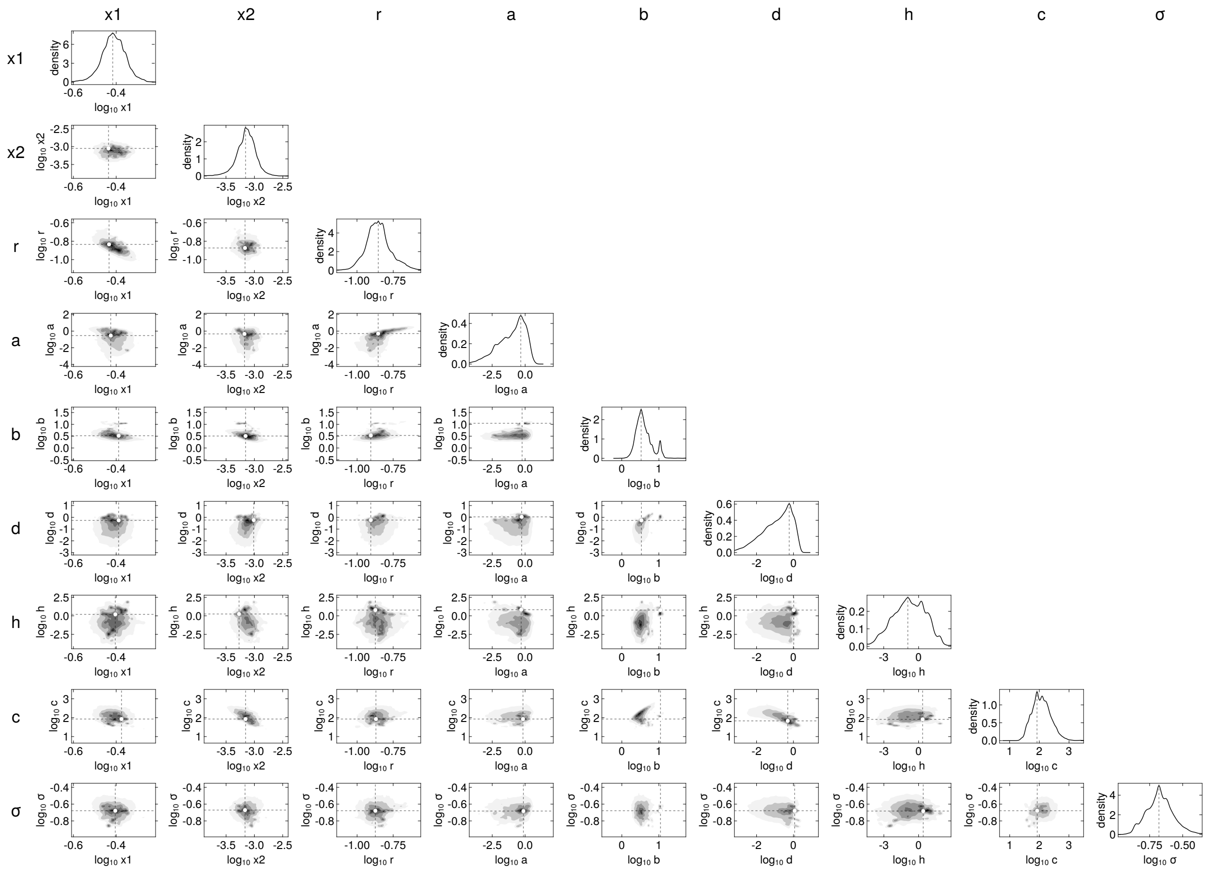


280

antigen_CD371_effector_CART_target_KO_model_Beddington_DeAngelis_activation_ratio_1_4.0

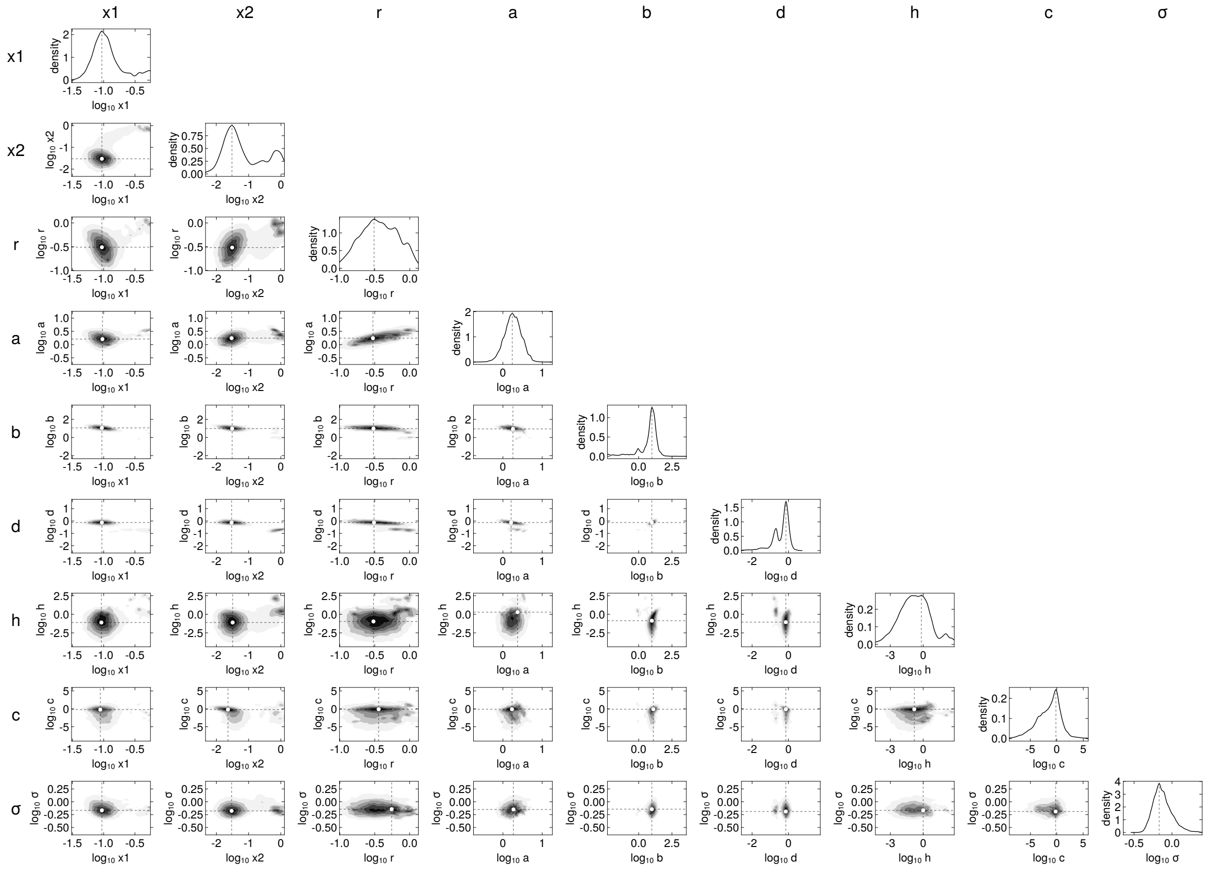


281

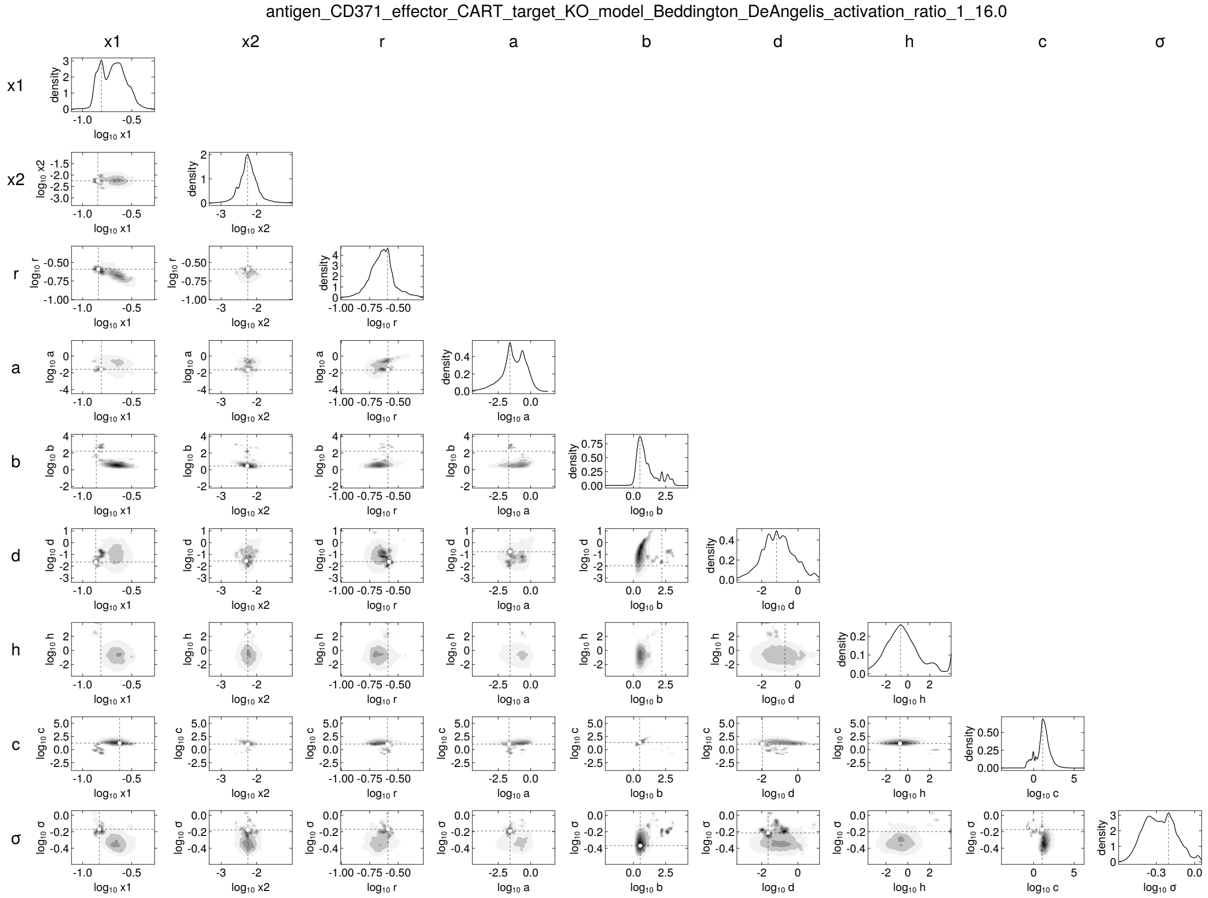
antigen_CD371_effector_CART_target_KO_model_Beddington_DeAngelis_activation_ratio_1_16.0
x1
x2
r
a
b
d
h
c
σ
density
log10 x1
log10 x2
log10 r
log10 a
log10 b
log10 d
log10 h
log10 c
log10 σ

282

antigen_CD371_effector_CART_target_KO_model_Beddington_DeAngelis_activation_ratio_1_40.0

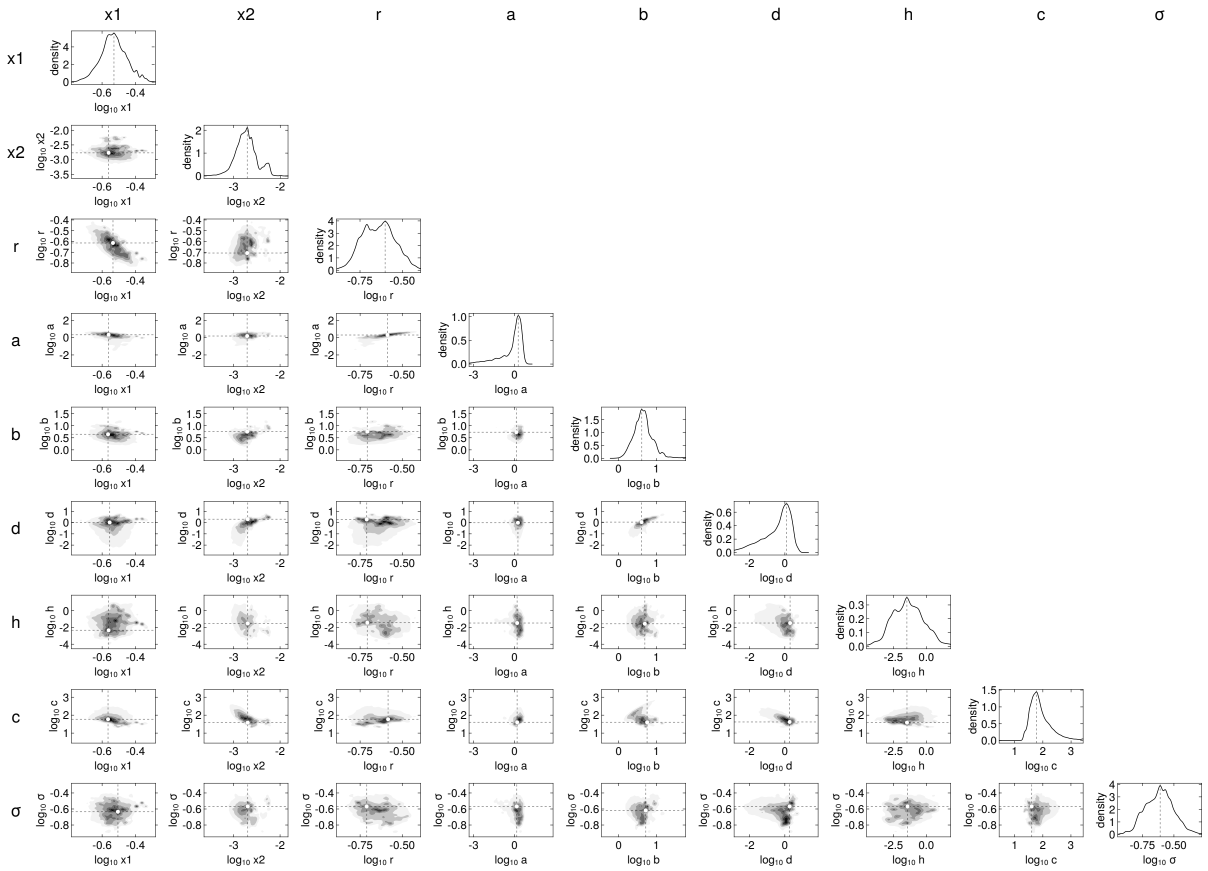

283

antigen_CD371_effector_CART_target_KO_model_Beddington_DeAngelis_activation_ratio_1_60.0

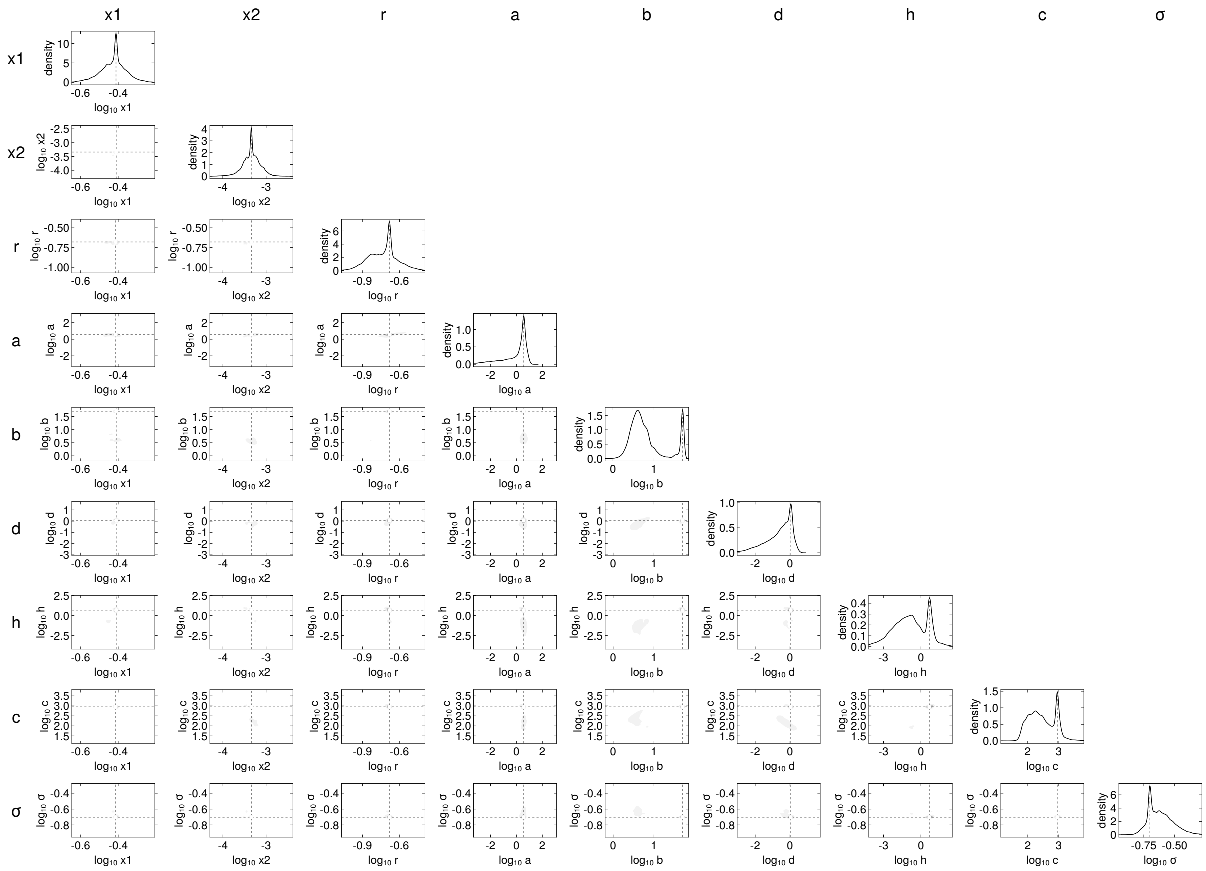
